# Further biochemical profiling of *Hypholoma fasciculare* metabolome reveals its chemogenetic diversity

**DOI:** 10.1101/2020.05.28.122176

**Authors:** Suhad A.A. Al-Salihi, Ian Bull, Raghad A. Al-Salhi, Paul J. Gates, Kifah Salih, Andy M. Bailey, Gary D. Foster

## Abstract

There is a desperate need in continuing the search for natural products with novel mechanism to battle the constant increase of microbial drug resistance. Previously mushroom forming fungi were neglected as a source of novel antibiotics, due to the difficulties associated with their culture preparation and genetic tractability. However, modern fungal molecular and synthetic biology tools, renewed the interest in exploring mushroom fungi for novel therapeutics. The aim of this study was to have a comprehensive picture of nine basidiomycetes secondary metabolites (SM), screen their biological and chemical properties to describe the genetic pathways associated with their production. *H. fasciculare* revealed to be highly active antagonistic species, with antimicrobial activity against three different microorganisms - *Bacillus subtilis, Escherichia coli* and *Saccharomyces cerevisiae*-. Extensive genomic comparison and chemical analysis using analytical chromatography, led to the characterisation of more than 15 variant biosynthetic gene clusters and the first identification of a potent antibacterial metabolite-3, 5-dichloromethoxy benzoic acid (3, 5-D)-in this species, for which a biosynthetic gene cluster was predicted. This work demonstrates the great potential of mushroom forming fungi as a reservoir of bioactive natural products which are currently unexplored, and that access to their genomic data and structural diversity natural products via utilizing modern computational analysis and efficient chemical methods, could accelerate the development and applications of such distinct molecules in both pharmaceutical and agrochemical industry.

## Introduction

Mushroom forming fungi are recognised to produce a plethora of special compounds to defeat competitive organisms coexist in their ecosystem. These are usually known as specialised secondary metabolites or natural products which often have bioactive properties such as antimicrobial, antitumor and insecticide amongst others (1). However, historically most secondary metabolites have been derived from bacteria and ascomycetes fungi, overlooking basidiomycetes as a potential repository of novel natural products. There are reports in the literature that only 1 in 5 of currently used antibiotics are developed from fungi (2). This is mainly due to complications involved in their laboratory cultivation. Compared to other groups of fungi, mushroom forming basidiomycetes grow in a slow rate and reproduce using dikaryotic cell type, making their genetic tractability investigations more sophisticated. It was therefore predicted that mushroom forming basidiomycetes may produce certain novel compounds that are yet to be exploited (3). Although, basidiomycetes are known to produce certain mycotoxins with significant biological effects in human and agriculture, such as psilocybin from *Psilocybe spp*. and strobulin from *Strobilurus tenacellus* (4 & 5), the main group of natural products that have been isolated from basidiomycetes fungi are; terpenoid and halogenated compounds. These involves tetrachlorinated phenols, illudanes, sterpurnes, and illudalanes, all of which are largely produced by fungi with one exception of the illudalanes which are also produced by some plants (6 & 7).

Given the large number of untapped Agaricales species, there is likely to find promising potentials within such group of organisms. *H. fasciculare* (previously known as *Naematoloma fasciculare*), is an inedible (in most Europe) wood decay mushroom forming basidiomycetes, which was recognised as sulphur tuft, due to its way of growing (tight cluster tufts) and the bright sulphur-yellow colour of its cap. Several field studies have shown the capability of this fungus in controlling other wood decaying organism’s colonization that are coexist its niche (8). Although, *H. fasciculare* was reported in the literature as a rich source of terpenoids and organohalogens natural products, including fascicularones and anisaldehyde metabolites, their pharmaceutical properties and biological synthesis remains unsolved (all chemical produced by *H. fasciculare* from 1967 to 2019 are listed in figure S1in the supplementary information). In order to gain a greater understanding of the biology and the chemistry of *H. fasciculare* metabolome, we carried out a series of genetic manipulations, bioactivity assays and chemical analyses of *H. fasciculare* crude and pure extracts to detect their antimicrobial activities and to further investigate the metabolic potential of this fungus in an attempt to bridge the gap between natural product innovation and antibacterial resistance.

## Material and methods

### *H. fasciculare* genome mining

Our previous investigation of *Hypholoma* species genomes suggested that those fungi are mainly produce terpenoid compounds with a range of cyclization patterns (Al-Salihi et al., 2019). However, subsequent in-depth BLAST search of functionally characterized core enzymes (see supplementary information Table S1 for enzymes details) selected from different fungi, resulted in additional BGCs in both *Hypholoma* species (see supplementary information Table S2). Introns and exons of selected scaffolds were predicted using a combination of Softberry and Local BLAST search, allowing the subsequent functional analysis of predicted biosynthetic gene clusters. Annotation of the predicted open reading frames of each of the submitted contig was then carried out using Artemis. Protein BLAST search on NCBI was then done for each gene to predict their function. At least 10 genes of either direction of the predicted SM core enzyme were annotated. We then manually curated our *Hypholoma* BGCs by blast search them against *H. sublateritium* genome on JGI.

### Antisense plasmid construction and Agrobacterium transformation *H. fasciculare* terpene synthase silencing

#### RT-PCR

To link each predicted terpene synthase to at least one of the previously reported natural molecules from *H. fasciculare*, and to further confirm the accuracy of our *in silico* prediction, we performed RNAi-mediated gene silencing. During our *in silico* analysis it was appeared that terpene synthases are the most abundance core enzymes of *H. fasciculare* SM biosynthetic genes. Enzymes were selected according to their carbon cyclization pattern, including five representatives (HfasTerp-255, HfasTerp-94A, HfasTerp-94B, HfasTerp-105) 1,11 carbon cyclization (as half of the predicted terpene synthase of *H. fasciculare* were of 1,11 cyclization patterns. The remaining genes were as follow: HfasTerp-147 for 1,10 3RNNP, HfasTerp-804 for 1,6 3R/S-NPP, HfasTerp-342 and HfasTerp-179 for 1, 10, E, E-FPP. The separately clustered HfasTerp-85b was also included in this investigation.

Prior to antisense plasmid construction, RT-PCR was carried out for the genes selected, to confirm their predicted splicing pattern. introns of all nine selected terpene synthases plus two housekeeping genes (*gpd* and *β-tubulin*) were predicted using a combination of SoftBerry and Artemis. Amplification of 150-250 bp DNA fragment spanning at least one intron was carried out for each gene. RNA was extracted from growing mycelial cultures which their antimicrobial activity was already confirmed. cDNAs were then synthesised with Oligo(dT)18 primers, and their integrities were examined prior their use for RT-PCR.

#### Construction of antisense vector targeting argininosuccinate gene

Due to the lack of efficient genome alteration techniques in basidiomycetes, downregulating of biosynthetic genes expression in such species has been neglected in the past and the consequences of such approaches remain unclear. Argininosuccinate synthetase is an essential protein for fungus growth and successful silencing of its biosynthesis, would display irregular growth pattern and allowing for quick and simple assessment of any potential silencing. Argininosuccinate synthetase silencing experiments were therefore carried out alongside the selected terpene synthases to detect any potential role of selected terpene synthases in *H. fasciculare* fitness via phenotypic comparison with Argininosuccinate synthetase silenced lines. For efficient analysis, we selected arginine a visualizable essential element for protein synthesis.

The published sequences of *H. sublateritium* argininosuccinate synthetase was blast search against *H. fasciculare* genome. A gene with 93% identity was identified in *H. fasciculare* contig-63. Argininosuccinate antisense plasmid consisted of pCAMBIA0380YA backbone, 500 bp of *H. fasciculare* argininosuccinate gene, which was inserted (in the antisense orientation) between *H. sublateritium* gpd promoter and TrpC terminator and the hygromycin cassette (*hph* gene under *A. bisporus gpd*II promoter and CaMV35S terminator). The verified argininosuccinate and terpene synthases plasmids, were transformed into strain LBA4404 of *A. tumefaciens* individually and used in *H. fasciculare* silencing experiments.

#### Chemical profiling of *H. fasciculare* silenced lines

Fungal plugs of silenced transformants were individually inoculated into 100 ml of MEB (15 g/L malt extract broth) in a 250 ml flask and incubated at 25°C and 200 rpm for 21 day. The ethyl acetate chemical extraction (9) was used for metabolites extractions. 20 µl (final concentration of 5 mg/ml) of each crude extract was then subjected to HPLC for direct comparison between the wild type and silenced lines in terms of their chemical constituents.

#### Generating *A. oryzae* transformants of Key enzymes

To avoid the potential issue associated with intron miss-splicing, the cDNA templates for the selected genes (HfasTerp-94A, HfasTerp94B, HfasTerp179 and HfasTerp344), were synthesized. The cDNA version of the sesquiterpene synthases (Cop-1, Cop-2, Cop-3, Cop-4, Omph-6 and Omph-7) were kindly provided by Schmidt’s group (10 and 11). *A. oryzae* transformants were generated for the ten selected enzymes and chemically analysed using the protocol described in (12).

## Results

### Bioassay

We bio-assayed nine basidiomycetes to investigate their ability to produce bioactive SM over a range of solid media (see supplementary information for method details), from which the two Strophariaceae members -*H. fasciculare* and *H. sublateritium*-displayed noticeable antimicrobial activity against the three challenged microbes (Fig.1 A & B). In contrast, *Paxillus involutus* showed no activity against any of the tested microbes. Variable inhibition zones were produced by the remained basidiomycetes (see supplementary information Figs. S2 to S7 for media, test microbe and clearing zones diameter description).

**Figure 1:**
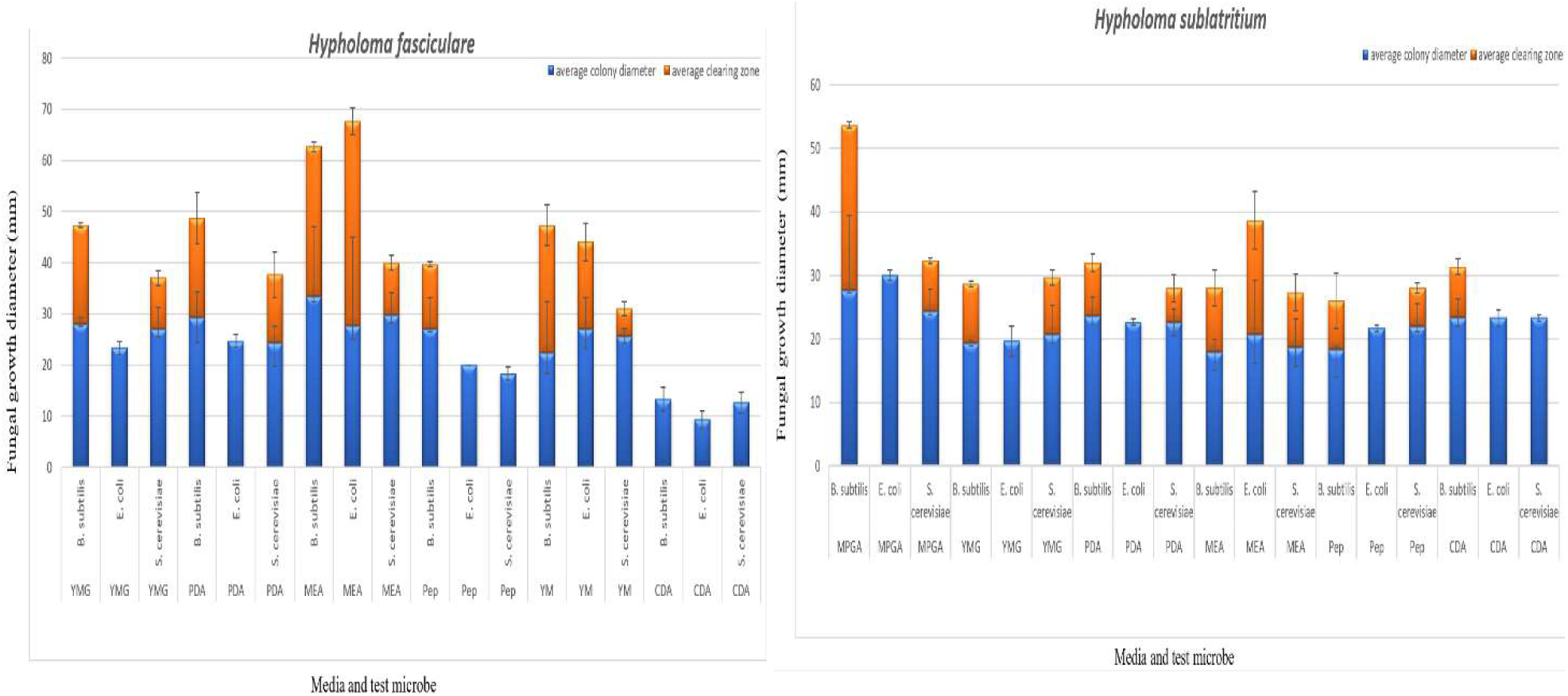
Bioassay test to evaluate the antimicrobial activity of *H. fasciculare* growing on different media against *B. subtilis, E. coli* and *S. cerevisiae*. YMG = yeast extract malt, PDA = potato dextrose agar, MEA = malt extract agar, Pep = peptone agar, YM = yeast malt extract, CDA = Czapek Dox agar. Error bars indicates the standard deviation of three technical replicates measurement for both fungal colony diameter (column in blue) and inhibition zone diameter (column in red). B- Bioassay test to evaluate the antimicrobial activity of *H. sublateritium* growing on different media against *B. subtilis, E. coli* and *S. cerevisiae*. MPGA = malt peptone glucose agar, YMG = yeast extract malt, PDA = potato dextrose agar, MEA = malt extract agar, Pep = peptone agar, CDA = Czapek Dox agar. Error bars indicates the standard deviation of three technical replicates measurement for both fungal colony diameter (column in blue) and inhibition zone diameter (column in red).

### Bioautography

Due to the broad similarity between the chemical profile of the crude extracts of both mycelia and supernatant of the five tested cultures for both *Hypholoma* (data not shown), it was decided to use homogenized cultures for metabolites extraction, where the whole culture was homogenised prior to the chemical extraction. The chemical extracts of CSO, YMG, PDB, CGC and MEB for both *Hypholoma*, were then spotted on the TLC plates and developed in different systems of solvents; non-polar, polar and semi-polar, which were then tested against *B. subtilis* (see supplementary information Figs. S8-S11 for TLC plate examples).

The antibacterial activity of the crude extracts loaded on TLC plates in a polar system showed one big clearing zone (size ranged from 20-26 mm), and the highest antibacterial activity (26 mm) was observed in YMG crude extracts amongst the other extracts of *H. fasciculare* and in PDB (25 mm) crude extract of *H. sublateritium*. Figs. 12-14 in the supplementary information show some examples of bioautography plates of *H. fasciculare*.

**Figure 2:**
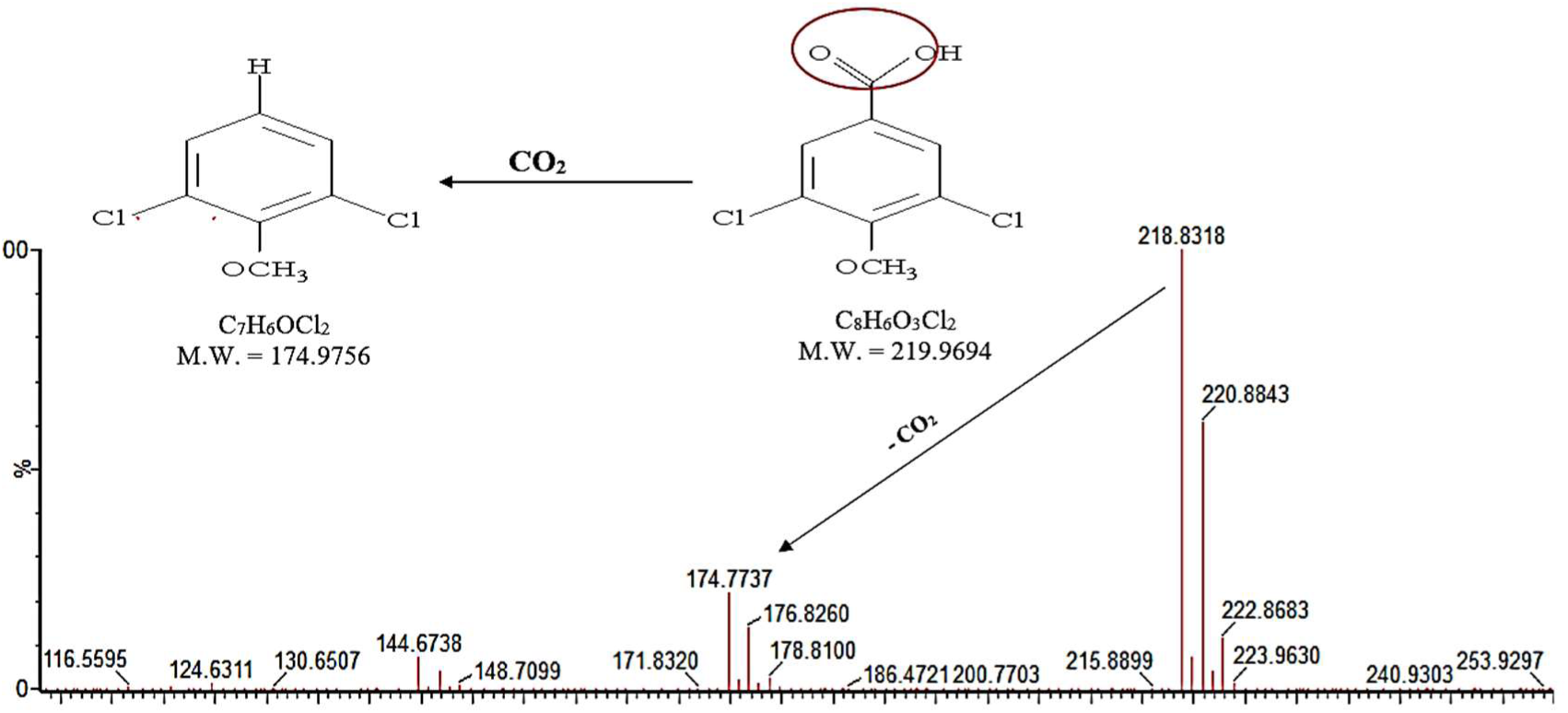
High resolution mass spectrum (-ESI) of fraction C, shows the fragmentation pattern of putative 3,5 dichloro-4-methoxy benzoic acid and, a proposed loss of carboxylic group.

**Figure 3:**
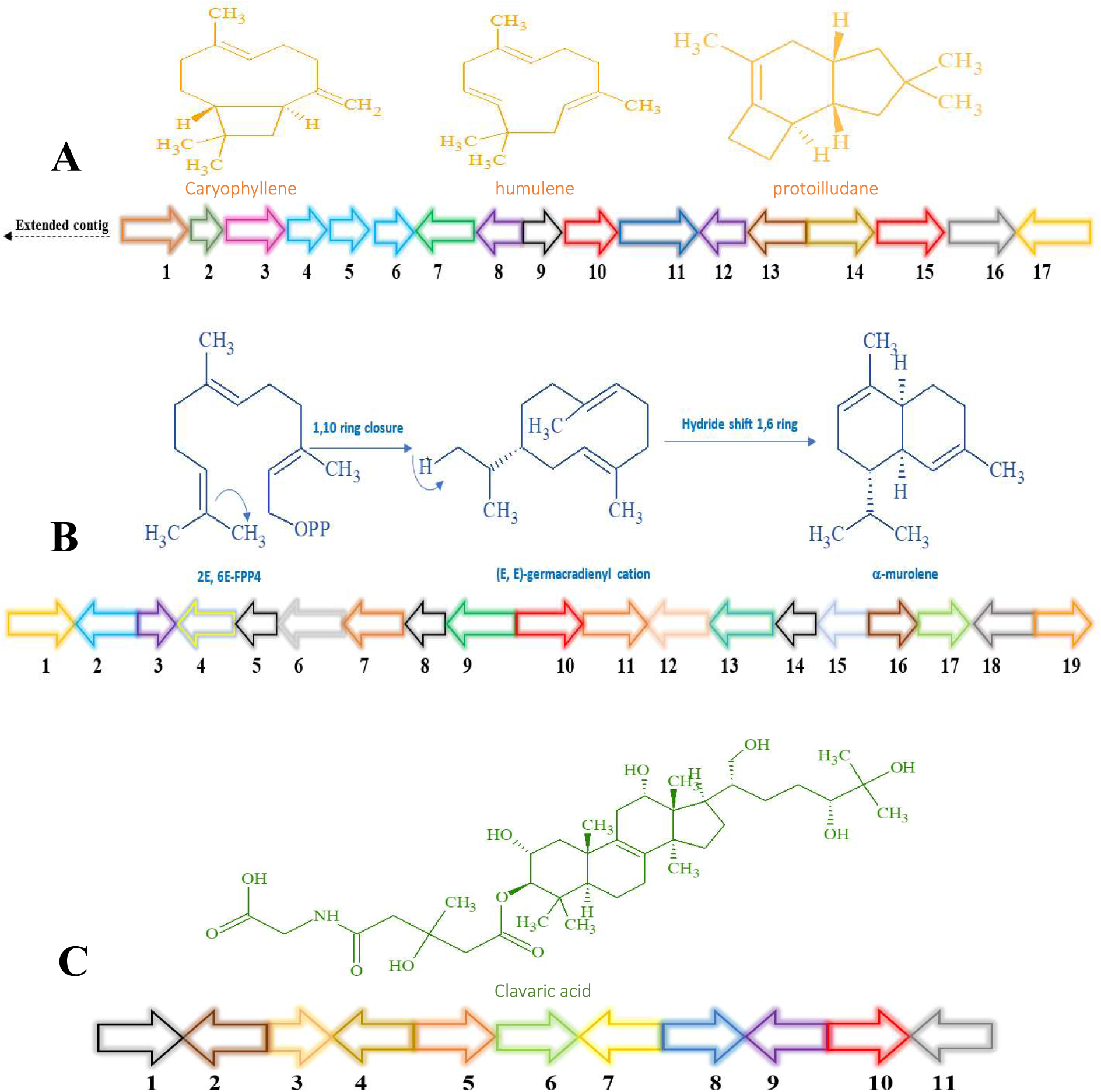
**A- HfasTerp-94a & b:** 1- Serine/threonine kinase, 2- Zinc carboxypeptidase, 3- Hypothetical protein, 4- Short chain dehydrogenase 1, 5- Short chain dehydrogenase 2, 6- Short chain dehydrogenase 3, 7- Tyrosinase, 8- F box domain, 9- Anchor signaling protein, 10- Terpene synthase-B, 11- Splicing co-activator, 12- F box domain, 13- Glucose transporter, 14- Peptide transporter, 15- Terpene synthase-A, 16- Aromatic ring hydroxylase, 17- Glycoside hydrolase. **B- HfasTerp-179:** 1- Alpha/beta-hydrolases, 2- Aspartyl protease, 3- E3 ubiquitin-protein ligase, 4- NAD- dependent histone deacetylase, 5- Hypothetical protein, 6- Transcription factor, 7- GMC oxidoreductase, 8- Hypothetical protein, 9- Cytochrome P450, 10- Terpene synthase, 11- GMC oxidoreductase, 12- Fructose 2,6 bisphosphatase, 13- Kinase like protein, 14- Hypothetical protein, 15**-** DNA repair endonuclease, 16- Telomerase transcriptase, 17- Monooxygenase FAD, 18- Glycoside hydrolase, 19- Alpha ketogluturate. **C- HfasTerp-804:** 1- Hypothetical protein, 2- Flavin-containing amine oxidase, 3- Pyruvate decarboxylase, 4- Cytoplasmic protein, 5- Phosphomannomutase, 6- Delta DNA polymerase, 7- Putative transcription factor, 8- 60S ribosomal protein translation, 9- Actin-related protein, 10- Oxidosqualene clavarinone cyclase, 11- Nuclear condensin complex.

**Figure 4:**
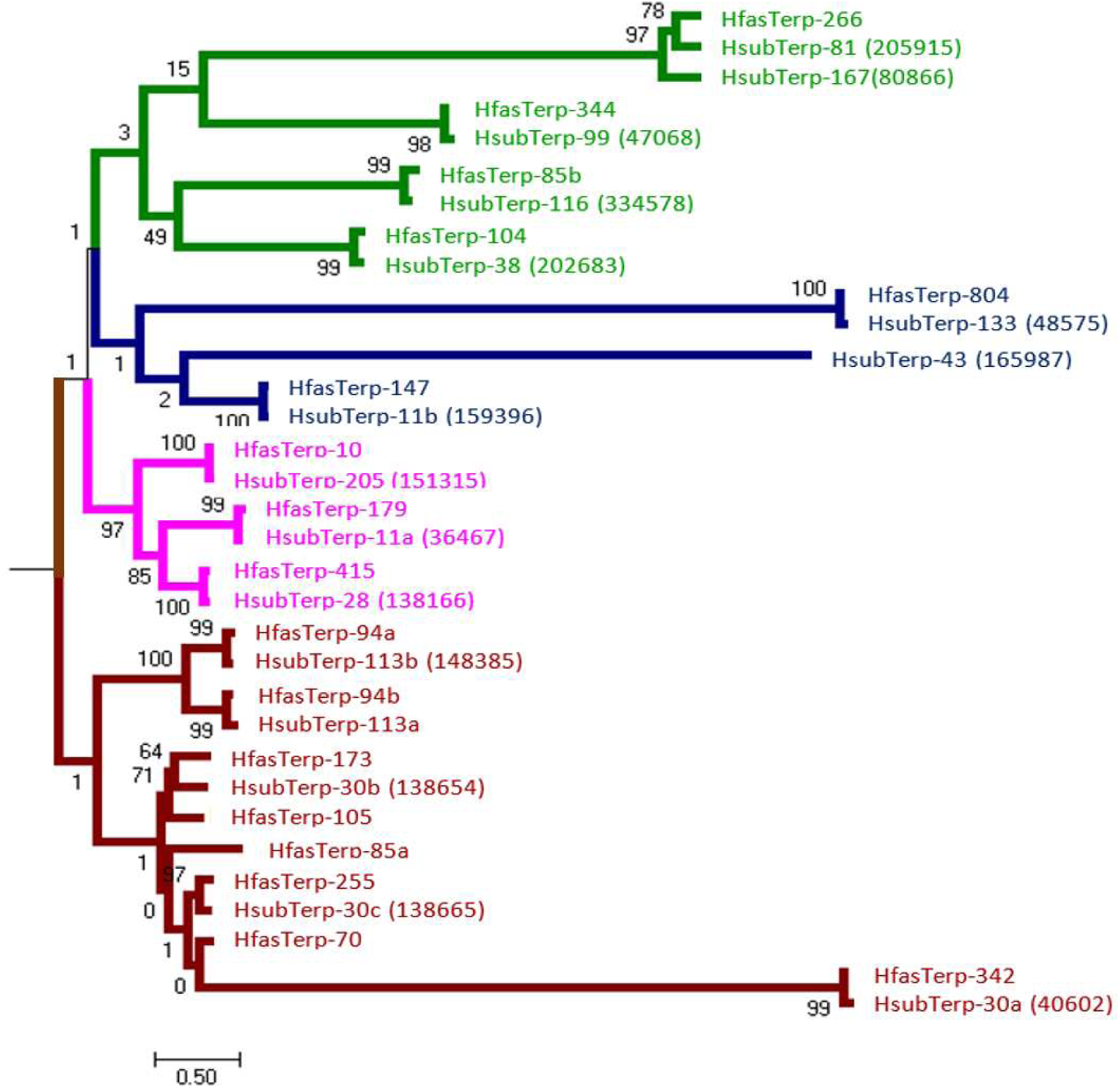
Maximum likelihood tree of both *Hypholoma* putative terpene synthases. The clades compromising Hfas putative terpene synthases based on sequences homologs and their initial cyclisation reaction. The *H. sublateritium* gene sequences were obtained from JGI and curated using Artemis. Contigs or scaffold and accession numbers are shown adjacent to species abbreviations.

**Figure 5:**
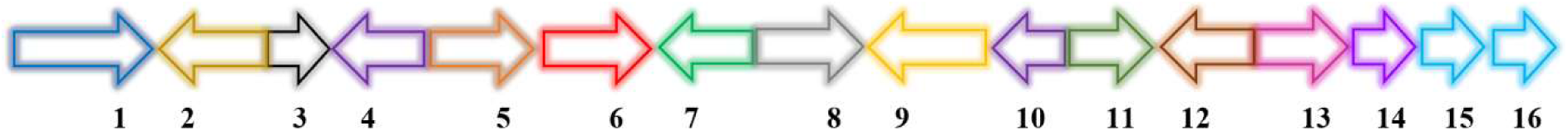
**HfasTerp-147;** 1- Complex I intermediate- associated protein. 2- Protein binding. 3- F-box. 4- Histon methyltransferase. 5- RNA polymerase. 6- Terpene synthase- 11. 7- Cystathionine beta-lyase. 8- Pyruvate carboxyltransferase. 9- Dolichol kinase. 10- Zinc finger. 11- Splicing factor. 12- Proliferation- associated protein. 13- Chaperone regulator. 14- Enoyl-Co A hydratase. 15- Aspartic peptidase. 16- Aspartic peptidase.

**Figure 6:**
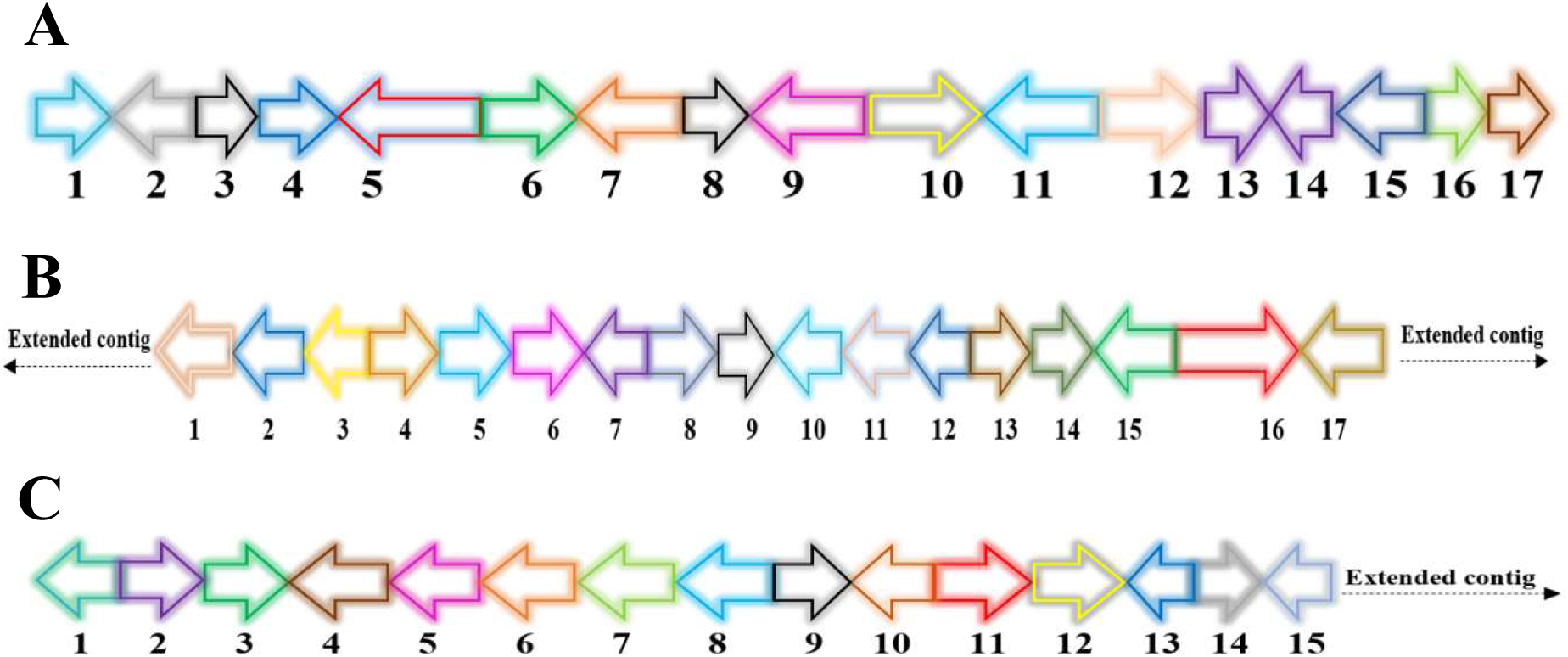
**A- HfasPKS-221 (74 Kb): 1-** Triphosphate hydrolase, 2- Multidrug resistance, 3- Hypothetical protein, 4- Alpha-beta hydrolase, 5- PKS1, 6- Zinc finger H2C2, 7- Kinesin-like protein, 8- Hypothetical protein, 9- Actin binding protein, 10- DNA binding protein, 11- Acetylcholinesterase, 12- Actin depolymerizing factor, 13- Metalloproteases, 14**-** Metalloproteases, 15- Metalloproteases, 16- N- acetylglucosaminyltransferase, 17- Transcription factor. **B- HfasNRPS-29 (76 Kb):** 1- mRNA-decapping enzyme, 2- Peptidase aspartic, 3- Glycoside hydrolase, 4- Polyadenylate-binding protein, 5- Mitochondrial carrier protein, 6- Ubiquitin-protease, 7- Acid phosphatase, 8- AMP-dependent synthetase and ligase, 9- Exocyst complex, 10- DUF-domain protein, 11- Carbohydrate transporter, 12- Adenine DNA glycosylase, 13- Cargo transport protein, 14- WD-repeat containing nucleolar rRNA, 15- Oxidoreductase, 16- NRPS, 17- Acyl-CoA N-acyltransferase. A homologous biosynthetic gene cluster was found in scaffold 100 of H. sublateritium genome. **C- HfasSidA-14 (98Kb):** 1- Transcription factor, 2- Sterol desaturase, 3- Cytochrome P450, 4- Triphosphate hydrolase, 5- Calcium transporting, 6- Glycosyltransferase, 7- GMC oxidoreductase, 8- Amino acid transporter, 9- Zinc binding protein, 10- ABC transporter, 11- Sid A, 12- Glycoside hydrolase, 13- Transporter, 14- RNA associated protein, 15- F-box.

**Figure 7:**
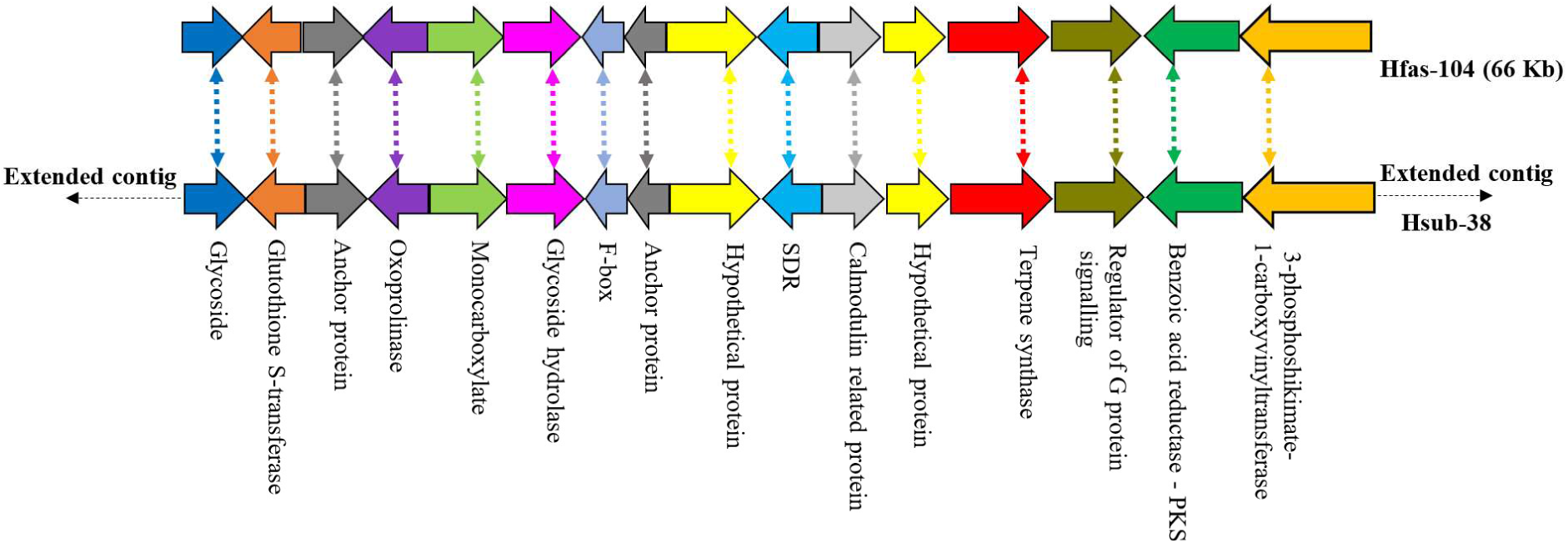
Predicted gene cluster of *H. fasciculare* and the homologous cluster of *H. sublateritium*.

**Figure 8:**
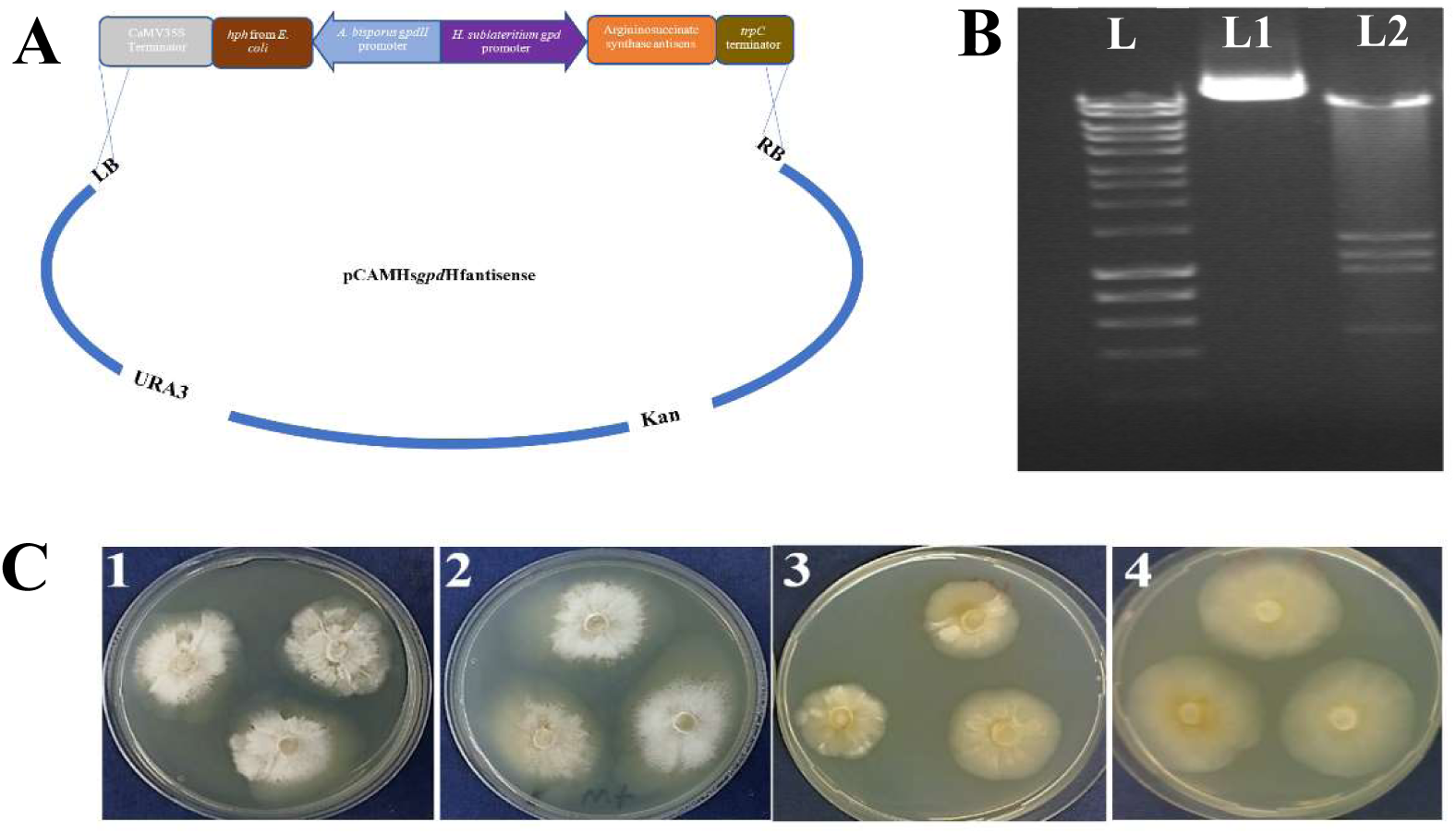
**A-** A schematic representing the antisense vector pCAMHs*gpd*Hfass used for targeting argininosuccinate gene in *H. fasciculare*. pCAMBIA0380YA was linearized by *Bam*HI and the hygromycin and antisense cassettes were inserted by yeast homologous recombination. **B-** Restriction enzyme (EcoRI) analysis of the silencing plasmids showing the correct construction. L = 5 *μ*L of hyperladder I, Lane 1 and 2 contain 5 *μ*L of pCAMHs*gpd*Hf-ass miniprep and digestion reaction respectively. **C-** *H. fasciculare* wild type and antisense transformant-14, showing differences in colony growth rate on PDA with and without arginine supplementation. *1- H. fasciculare* wild type on PDA medium. 2- *H. fasciculare* wild type on PDA medium supplemented with 5mM of arginine. 3- *H. fasciculare* antisense transformant-49 on PDA medium. 4- *H. fasciculare* antisense transformant-49 on PDA medium supplemented with 5mM of arginine.

**Figure 9:**
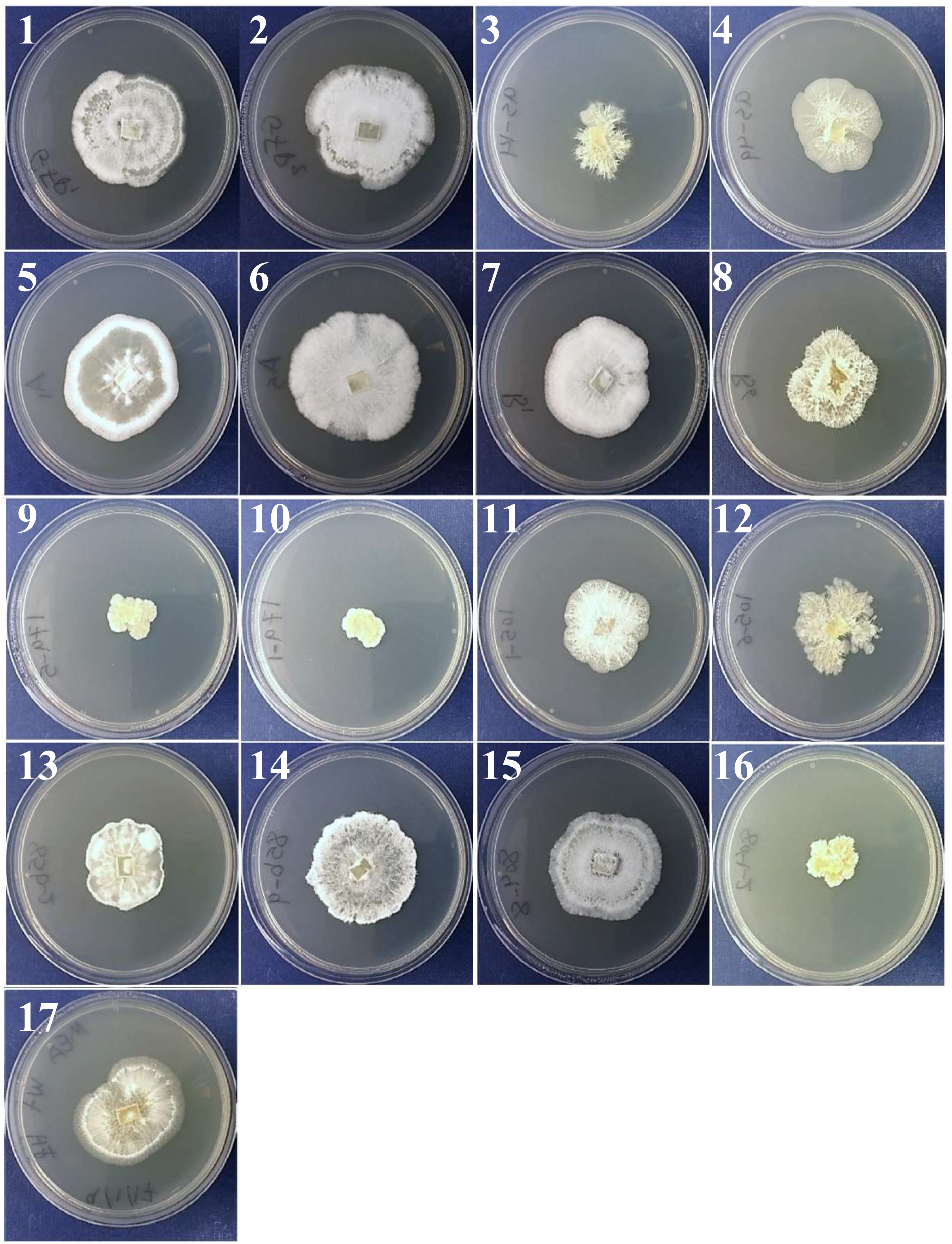
Shows the growth pattern of *H. fasciculare* silenced lines alongside the WT. Varied morphogenesis included rigid, fluffy, wrinkle, condense and lossy form of mycelia were observed among silenced lines. 1. HfasGFP-TR1, 2. HfasGFP-TR2, 3. Hfas-assTR14, 4. Hfas-assTR49, 5. HfasTerp94aTR1, 6. HfasTerp94aTR5, 7. HfasTerp94bTR1, 8. HfasTerp94bTR6, 9. HfasTerp179TR5, 10. HfasTerp179TR1, 11. HfasTerp105TR1, 12. HfasTerp105TR6, 13. HfasTerp85bTR2, 14. HfasTerp85bTR2, 15. HfasTerp804TR8, 16. HfasTerp804TR2, 17. HfasWT.

**Figure 10:**
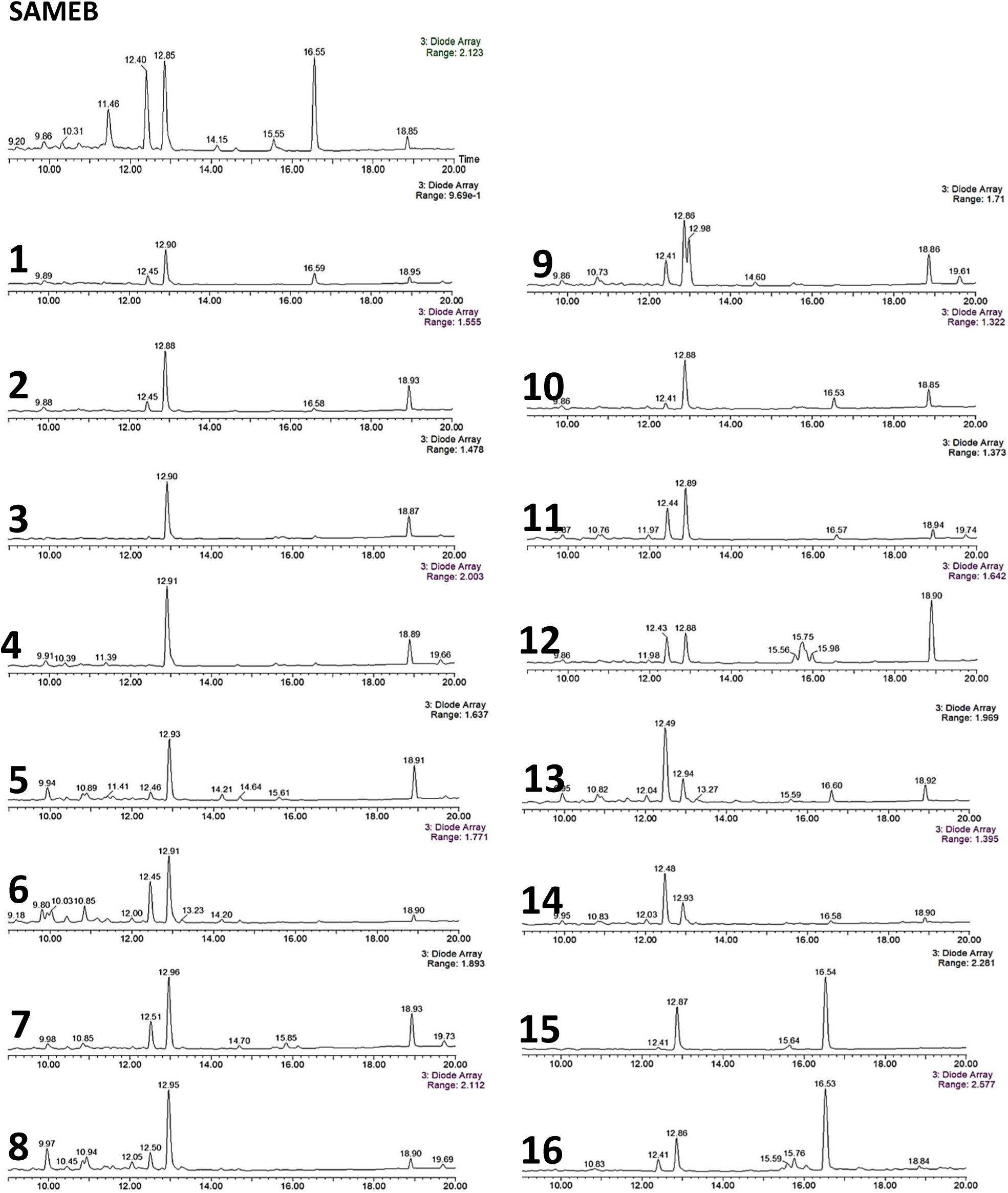
Diode array chromatograms for *H. fasciculare* WT and putative silenced transformants crude extracts. The production of main secondary metabolites was compared in the WT with two transformants of each silenced line. The genes investigated in this experiment were, argininosuccinate synthase and seven terpene synthases from different biosynthetic clusters. SAMEB = Hfas WT. 1 = Hfas-assTR14, 2- Hfas-assTR49, 3- HfasTerp85bTR2, 4- HfasTerp85bTR9, 5- HfasTerp94aTR1, 6- HfasTerp94aTR5, 7- HfasTerp94bTR1, 8- HfasTerp94bTR6, 9- HfasTerp105TR1, 10- HfasTerp105TR6, 11- HfasTerp179TR1, 12- HfasTerp179TR5, 13- HfasTerp342TR18, 14- HfasTerp342TR6, 15- HfasTerp804TR2, 16- HfasTerp804TR8.

**Figure 11:**
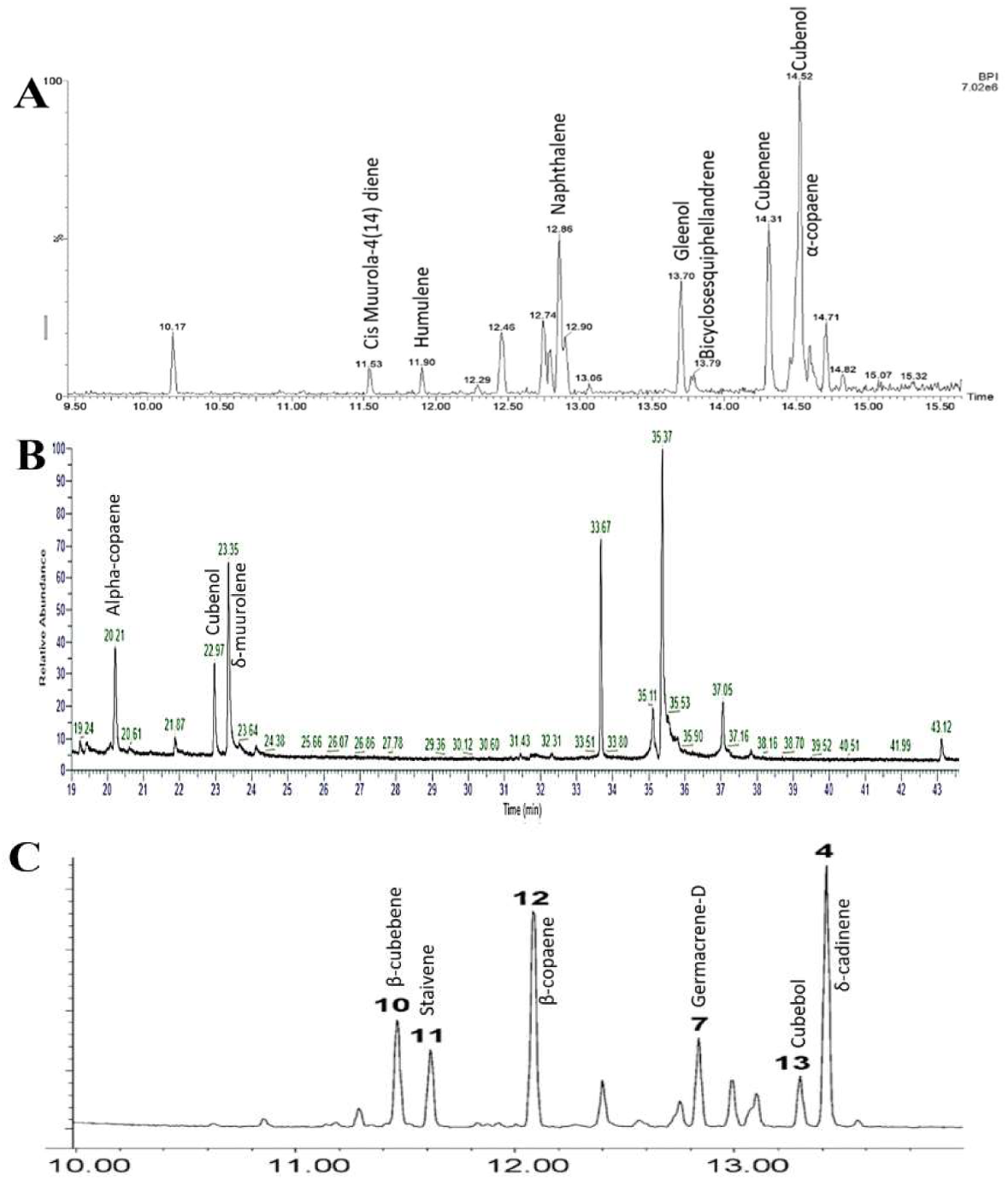
GC-MS comparison of the sesquiterpene synthase Cop4. A- GC-MS analysis of NSAR1 harbouring Cop4 using HP-5 MS quartz capillary column (30 m x 0.25 mm, 0.25 µm film thickness). B- GC-MS analysis of NSAR1 harbouring Cop4. C. GC-MS analysis of E. coli harbouring Cop4 using HP-1 MS capillary column (30 m ¥ 0.25 mm inner diameter ¥ 1.5 mm) adapted from (Agger et al., 2009).

**Figure 12:**
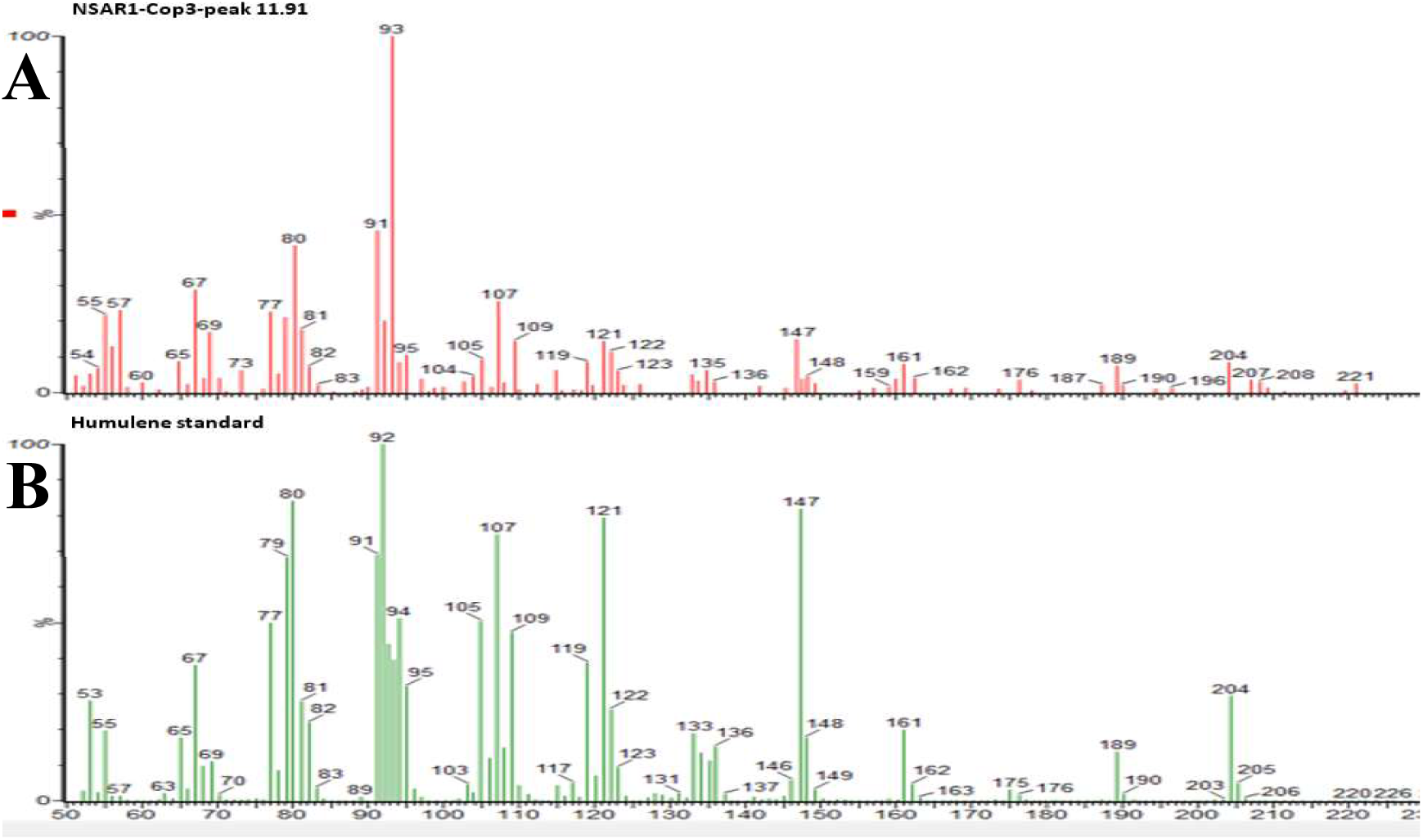
GC-MS spectrum comparison of A. Sesquiterpene synthase Cop3 and B. Humulene standard.

### High performance liquid chromatography (HPLC)

#### Chemical profiling of 3,5 dichloro-4-methoxy benzoic acid (3,5 D)

Our observation on bioautography assays, suggested similar chemical profiles for both *Hypholoma* species, at this point, we decided to narrow down the chemical purification of metabolites to *H. fasciculare* species. The variation in the RF of the observed active compounds in the bioautography plates, proposed different types of metabolites. We therefore selected CSO-1A, YMG, CGC and MEB for compounds purification. An initial analytical experiment of those crude extracts was run on HPLC, to select the most potential extract for metabolite purification. This has led to the detection of five main products; A, B, C, D and E (Fig. S15 supplementary information). Subsequently, isolation experiments were carried out to attempt the purification of the detected five major peaks.

Despite attempting different programme sets of fractionations, only two fractions (A and D) could be purified from *H. fasciculare* crude extracts. A comparison was then carried out between the mass spectra of the purified metabolites (A and D) and the masses of the so far reported compounds from *Hypholoma* genus. Leading to the characterization of metabolite A as fascicularon G (known sesquiterpene from *Hypholoma*). However, no match could be found between any previously characterised metabolites from *Hypholoma* and our purified metabolite (fraction D). We therefore subjected fraction D to high resolution mass (HRMS) to confirm its mass precisely and predict its molecular formula as well as. This run has returned with an aromatic molecules of chemical formula (C_8_H_6_O_3_Cl_2_) known as 3,5 dichloro-4-methoxy benzoic acid (3, 5 D). Looking at the ionization pattern of the negative mode of this run, two chlorines and a carboxyl group typical isotope could be observed, further confirming the acidic nature of metabolite D (Fig. 2).

To further confirm the chemical structure of compound D, we run NMR analysis for it, where the presences of a symmetrical benzoic acid molecule and two chloro substituent groups plus one methoxy group were confirmed via the 1H NMR spectrum. The correlation between the aromatic protons at 7.9 and 3.8 and carbons signals were also investigated and a molecule composite of 3,5-dichloro-4-methoxybenzoic acid was structurally confirmed via HMBC experiments (see supplementary information Figs. S16-S20).

#### Antimicrobial activity of 3,5 dichloro-4-methoxy benzoic acid (3,5 D)

The antibacterial activity of 3,5 D was assessed following the determination of its accurate mass and formula. The MIC method was performed through which, a gradient concentration of 100, 300 and 500 µg/disc of 3,5 D, was independently pipetted on 6 mm sterile filter disks and positioned on agar plates containing an overlay of the examined bacteria. Disks containing methanol and kanamycin, were used as negative and positive control respectively. As showing in (Table 1) 3, 5 D antibacterial activity against the examined bacteria; *Staphylococcus aureus* and *B. subtilis* was increasing with the increase of its concentration, where a maximum activity could be achieved with a 500 µg/disc.

**Table 1:** Antimicrobial activity of 3-5 D tested against *B. subtilis* and *S. aureus*.

| Microorganism | Clearing zone (mm) disc diameter 5mm |  |  |  |  |
| --- | --- | --- | --- | --- | --- |
|  | 100 µg/disc<br>Kanamycin<br>(+ve control) | 100 µl of 98%<br>Methanol<br>(-ve control) | 100 µg/disc of<br>3-5 D | 300 µg/disc of<br>3-5 D | 500 µg/disc of<br>3-5 D |
| *B. subtilis* | 12 | - | 12 | 16 | 20 |
| *S. aureus* | 11 | - | 10 | 14 | 20 |

#### Genomic investigations

To provide a comprehensive insight into *H. fasciculare* metabolomic, and potentially link them to their related biosynthetic gene clusters, we used Artemis tool to further analyse both *Hypholomas’* genomes sequences surrounding the predicted core enzymes of both *Hypholoma* (Figure 1 phylogenetic tree). 15 twined biosynthetic gene clusters were annotated for *H. fasciculare* and *H. sublateritium*, of which a representative of each family (categorised according to their core enzyme function) will be presented in the following sections (detailed description of all predicted BGCs can be found in the supplementary information Figs. S21-S29).

##### 1 HfasTerp-94 biosynthetic gene cluster

located in contig 94 of *H. fasciculare* genome, its terpene synthase was predicted as a member of 1, 11 E, E-FPP carbon cyclization enzymes group. This group of enzymes are responsible for the production of *trans*-humulyl cation sesquiterpenes such as humulene, caryophyllene and protoilludane (7).When the sequences of these genes were blast searched against *H. sublateritium* genome, an analogous biosynthetic gene cluster was found with one exception; SDR3 which could not be found (Fig. 3 A, B and C).

##### 2 HfasTerp-179 biosynthetic gene cluster

identified in contig 179 of *H. fasciculare*. HfasTerp-179 signature gene identified at the left end of *H. fasciculare* contig 179. The clade that this terpene synthase belongs to, is consist of genes that are identified as germacradienyl derivatives producers. Manual blast search of the core enzyme and its adjacent biosynthetic genes against *H. sublateritium* genome unveiled a match cluster in *H. sublateritium* scaffold 11. To further confirm HfasTerp-179 cluster boundary, we blast search the genes located to the left side of *H. sublateritium* terpene synthase. This revealed the presence of their homologous which located in contig 14 of *H. fasciculare*. The other two biosynthetic genes of Hfasterp-179 gene cluster were found in contig 615 of *H. fasciculare*, indicative of genome misassembling.

##### 3 HfasTerp-804 BGC

this cluster consisted of the terpene synthase which demonstrated high sequence similarity with the experimentally characterized oxidosequaline synthase located in scaffold 133 of *H. sublateritium* (key enzyme for the antitumor clavaric acid production in *H. sublateritium* (13).

##### 4 HfasTerp-255

deep sequences annotation of *H. sublateritium* scaffold 30 predicted three terpene synthases HsubTerp-30a, HsubTerp-30b and HsubTerp-30c. Sequences blast on NCBI and literature search, revealed HsubTerp-30a function as squalene synthase and HsubTerp-30c as potential protoilludane synthase. Blasting search those terpenes against *H. fasciculare* genome revealed high sequence similarity with several terpene synthases; the highest similarity (88%) observed between HfasTerp-255 and HsubTerp-30c (Fig. 4).

##### 5 HfasTerp-147 BGC

HfasTerp-147 represents the only predicted terpene enzyme of *H. fasciculare* that follows the 1, 10 3R NPP cyclisation pattern. Sequence matching of its sequences with *H. sublateritium* genome, resulted in significant homologous with a terpene synthase located in scaffold 11 of *H. sublateritium* genome. Subsequent analysis of the tailoring genes of *H. sublateritium* biosynthetic gene cluster suggested that HfasTerp-147 gene cluster was assembled into two different contigs; HfasTerp-458 and HfasTerp-147 (Fig. 5).

##### 6 HfasPKS-221 biosynthetic gene cluster

positioned in contig 221 of *H. fasciculare*. Subsequent comparison with *H. sublateritium* genome, revealed identical BGC suited in scaffold 53 (Fig. 6 A).

##### 7 HfasNRPS-29 biosynthetic gene cluster

identified in contig 29 of *H. fasciculare* and its twin cluster was found in scaffold 100 of *H. sublateritium* genome (Fig. 6 B).

##### 8 HfasSid-14 biosynthetic gene cluster

predicted in contig 14 of *H. fasciculare*, blasting sequences of its genes against *H. sublateritium* genome, resulted in an identical cluster located in scaffold 11 (Fig. 6 C).

##### 9 The 3,5 dichloro-4-methoxy benzoic acid (3,5 D) putative biosynthetic gene cluster

Among the shared hybrid biosynthetic gene clusters of *Hypholoma* spp., HfasTerp-104 and HsubTerp-38, appeared to have the necessary enzymes for the synthesis of compound 3, 5-D, such as 3-phosphoshikimate-1-carboxyvinyltransferase, benzoic acid reductase-PKS, SDR, glycoside hydrolase, highlighting their likely role in halogenated natural products synthesis (Fig. 7).

#### Silencing experiments

Correctly designed plasmids (Fig. 8A) were transferred into *H. fasciculare* using Agrobacterium mediated transformation following the protocol described in (14). Silencing efficiency were assessed by comparing hyphal growth pattern, gene expression, inhibition zone diameter and chemical profiles of transformants with the wild type.

For arginosuccinate synthetase silencing, randomly selected putative silenced colonies alongside the WT, were sub-cultured on PDA plate to investigate whether silencing arginosuccinate synthetase has an impact on their growth development. Several transformants showed reduction in their colony size, among which as-14 displayed the lowest growth rate (20 mm) compared to the wild type (30 mm), suggesting potential silencing in those lines. To further analyse this observed phenotype, replicates of transformant as-14 and the WT were subcultured on PDA plates with and without 5 mM of the supplement arginine. Following fortnight of incubation at 25°C, colony diameters were measured, and results indicated potential successful gene silencing in transformant Hfas-as-14, whereby silenced colonies demonstrated slower rate of growth and different form of mycelia compared to the wild type (Fig. 8C).

Following argininosuccinate silencing experiments, terpene synthase silencing transformation was carried out in similar manner. Classical selection on supplemented PDA plates was performed and two selected silenced transformants for each line including two argininosuccinate synthetase transformants, were further investigated for their biological activity using plate-based bioassay. Despite standardising the size of inoculum and culture conditions for all transformants, silenced lines displayed variation in terms of colony diameter and inhibition zone. The observed morphogenesis of silenced colonies was rigid, fluffy, wrinkle, condense and lossy form of mycelia (Fig. 9).

Of nine silenced lines, HfasTerp-94bTR6 and HfasTerp-85bTR9, displayed the highest reduction in their clearing zone; 38% and 45% respectively, while, transformant HfasTerp- 804TR8 and the argininosuccinate antisense Hfas-asTR49, produced larger inhibition zones (25%) compared to the wild type. Reduction from 12 to 32% was demonstrated by the remaining transformants (Table 2).

**Table 2:** Represent the average colony and clearing zone diameter of two selected putative antisense transformants alongside the wild type.

| **Colony on MEA plates** | **Average colony diameter (mm) of three technical replicates** | **St. deviation of three technical replicates** | **Average clearing zone (mm) of three technical replicates** | **St. deviation of three technical replicates** |
| --- | --- | --- | --- | --- |
| **HfasWT** | 27 | $\pm 0.7$ | 32 | $\pm 0.7$ |
| **HfasTerp94aTR1** | 30 | $\pm 1$ | 26 | $\pm 0.5$ |
| **HfasTerp94aTR5** | 33 | $\pm 0.7$ | 24 | $\pm 0.7$ |
| **HfasTerp94bTR1** | 29 | $\pm 0.5$ | 24 | $\pm 0$ |
| **HfasTerp94bTR6** | 28 | $\pm 0$ | 20 | $\pm 0.7$ |
| **HfasTerp85bTR2** | 30 | $\pm 1$ | 22 | $\pm 0$ |
| **HfasTerp85bTR9** | 26 | $\pm 1.4$ | 18 | $\pm 0.7$ |
| **HfasTerp105TR1** | 24 | $\pm 1$ | 24 | $\pm 1.4$ |
| **HfasTerp105TR6** | 27 | $\pm 0.5$ | 22 | $\pm 1.5$ |
| **HfasTerp179TR1** | 25 | $\pm 0.7$ | 28 | $\pm 0.7$ |
| **HfasTerp179TR5** | 19 | $\pm 0.7$ | 26 | $\pm 0.7$ |
| **HfasTerp342TR6** | 32 | $\pm 0.7$ | 26 | $\pm 1.5$ |
| **HfasTerp342TR18** | 29 | $\pm 1.4$ | 28 | $\pm 0.7$ |
| **HfasTerp804TR2** | 21 | $\pm 1$ | 32 | $\pm 0.7$ |
| **HfasTerp804TR8** | 20 | $\pm 0.5$ | 40 | $\pm 0.7$ |
| **Hfas-assTR14** | 27 | $\pm 0.7$ | 34 | $\pm 0.7$ |
| **Hfas-assTR49** | 27 | $\pm 0.7$ | 40 | $\pm 0.7$ |

#### Gene expression analysis of H. fasciculare silenced lines

Genes HfasTerp-94a, HfasTerp-94b and HfasTerp-105, *gpd* and *β- tubulin*, were used to detect their expression level in selected silenced transformants alongside the wild type. All transformants displayed reduction in their expression level attributed to the successful downregulating of their corresponding genes. Table 3 shows fold differences of selected genes over the conserved ones.

**Table 3:** Represent the RT-qPCR outcome of silenced lines. Each gene was examined against two references genes GAPDH and *β-tubulin*.

| Sample | Ct Target gene | Ct Reference gene (β-*tubulin*) | ΔCt (Target-Reference) |
| --- | --- | --- | --- |
| HfasTerp94aTR1 | 22.25 | 21.53 | 0.72 |
| HfasTerp94aTR5 | 23.8 | 23.24 | 0.56 |
| Hfas-WT94a | 20.74 | 20.11 | 0.63 |
| HfasTerp94bTR1 | 25.51 | 19.73 | 5.78 |
| HfasTerp94bTR6 | 25.81 | 21.11 | 4.7 |
| Hfas-WT94b | 20.68 | 21.11 | -0.43 |
| HfasTerp105TR1 | 26.26 | 21.94 | 4.32 |
| HfasTerp105TR6 | 28.93 | 20.76 | 8.82 |
| HfasWT 105 | 23.23 | 20.11 | 3.12 |
| Sample | Ct Target gene | Ct Reference gene (GAPDH) | ΔCt (Target-Reference) |
| HfasTerp94aTR1 | 22.25 | 17.22 | 5.03 |
| HfasTerp94aTR5 | 23.8 | 15.61 | 8.19 |
| HfasWT94a | 20.74 | 11.64 | 9.1 |
| HfasTerp94bTR1 | 25.51 | 16.20 | 9.31 |
| HfasTerp94bTR6 | 25.81 | 15.95 | 9.86 |
| HfasWT94b | 20.68 | 11.64 | 9.04 |
| HfasTerp105TR1 | 26.26 | 15.43 | 10.83 |
| HfasTerp105TR6 | 28.93 | 16.09 | 12.84 |
| HfasWT 105 | 23.23 | 11.64 | 11.59 |

#### Chemical analysis of silenced lines

Different levels of SM production were observed among silenced lines. Particularly, transformants; Hfas-assTR49, HfasTerp85bTR2, HfasTerp85bTR9, where the production of most of the molecules was reduced. However, the production of the newly characterized (in *H. fasciculare*) 3,5 D, showed no reduction in all transformants, indicating the involvement of different type of key enzyme in its biological synthesis (Fig. 10).

#### Heterologous expression of selected terpene synthase

Although silencing constructs have proven successful for functional studies in *H. fasciculare*, their role in linking sesquiterpene metabolites to their specific biogenetic genes was limited. We therefore adapted the vector pTYAGS-arg to express selected sesquiterpene synthases in *A. oryzae* and, to further assess whether using *A. oryzae* as expression host, as well as using different GCMS machine, would influence the expression outcome of some selected genes, we used our *A. oryzae* transformants from previous work (12), as well as six previously characterised fungal sesquiterpene synthases from two different basidiomycetes; *Omphalotus olearius* and *Coprinopsis cinerea* -cop-1, cop-2, cop-3, cop-4, omp-6 and omp-7- (10, 11) as controls. Seven verified constructs, pTYGS-argHfasTerp179, pTYG-argCop-1, pTYG- argCop-2, pTYG-argCop-3, pTYG-argCop-4, pTYG-argOmp6 and pTYG-argOmp7; were independently transferred to *A. oryzae* strain NSAR1 protoplasts via PEG mediated transformation reactions. Subsequent GC-MS analysis on previous (pTYGS-argHfasTerp94a, pTYGS-argHfasTerp94b and pTYGS-argHfasTerp344) and current transformants, confirmed our previous results of HfasTerp94a, HfasTerp94b and HfasTerp344, through the replication of humulene and caryophyllene production, although with lower level. The production of most of the major products of two controls -pTYG-argCop-3, pTYG-argCop-4- were confirmed as well previously uncharacterised humulene in Cop-3 extract (Figs. 11 & 12). However, no product could be observed for HfasTerp179 *A. oryzae* transformant.

#### Co-expression and chemical analysis of selected biosynthetic genes of HfasTerp94 gene cluster

Adjacent genes (SDR1, SDR2, SDR3 and tyrosinase) of the HfasTerp94 were selected for co-expression with NSAR1-humulene synthase. Due to the unsuccessful attempts of full-length cDNA amplification of selected genes, an alternative approaches of fragments amplification were chosen. Our *in silico* analysis predicted two exons for each SDR gene. Accordingly, four pairs of primers with 60 bp were used to amplify the two exons of each SDR from *H. fasciculare* gDNA. pTYGS-arg backbone was used for fragments recombination (Fig. 13). However, due to the prediction of several introns within tyrosinase gene, a synthetic version was used.

**Figure 13:**
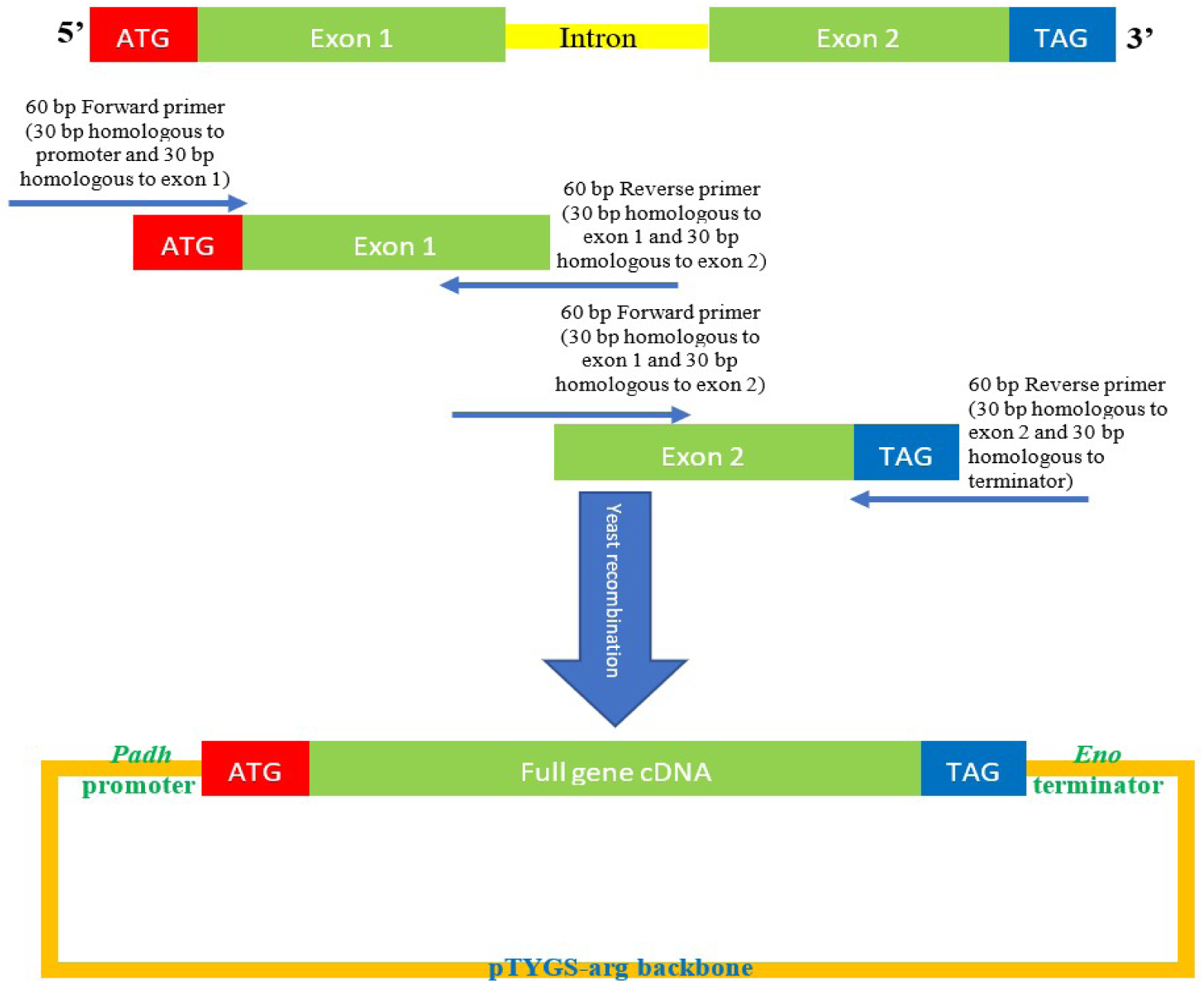
Schematic representation of the principle of constructing the pTYGS-arg-SDR plamid. Each SDR gene consisted of two exons and one intron. Exons shown in green. Intron shown in yellow. Overlapping primers shown as blue arrows. pTYGS-arg backbone was digested with *Asc*I and the region between *Padh* and *Teno* was replaced by the created full cDNA of one of the SDR genes.

Following successful integration of the selected genes within *A. oryzae* genome, mass spectrum comparison between the five produced transgenics (NSAR1-humulene synthase- SDR1, NSAR1-humulene synthase-SDR2, NSAR1-humulene synthase-SDR3, NSAR1- humulene synthase-SDR1-SDR2, NSAR1-humulene synthase-SDR1-SDR2-Tyrosinase) and NSAR1-humulene synthase, NSAR1-humulene synthase-SDR, crude extracts was carried out, and at least one new metabolite was detected in each transformant crude extract (Figs. 14 and 15).

**Figure 14:**
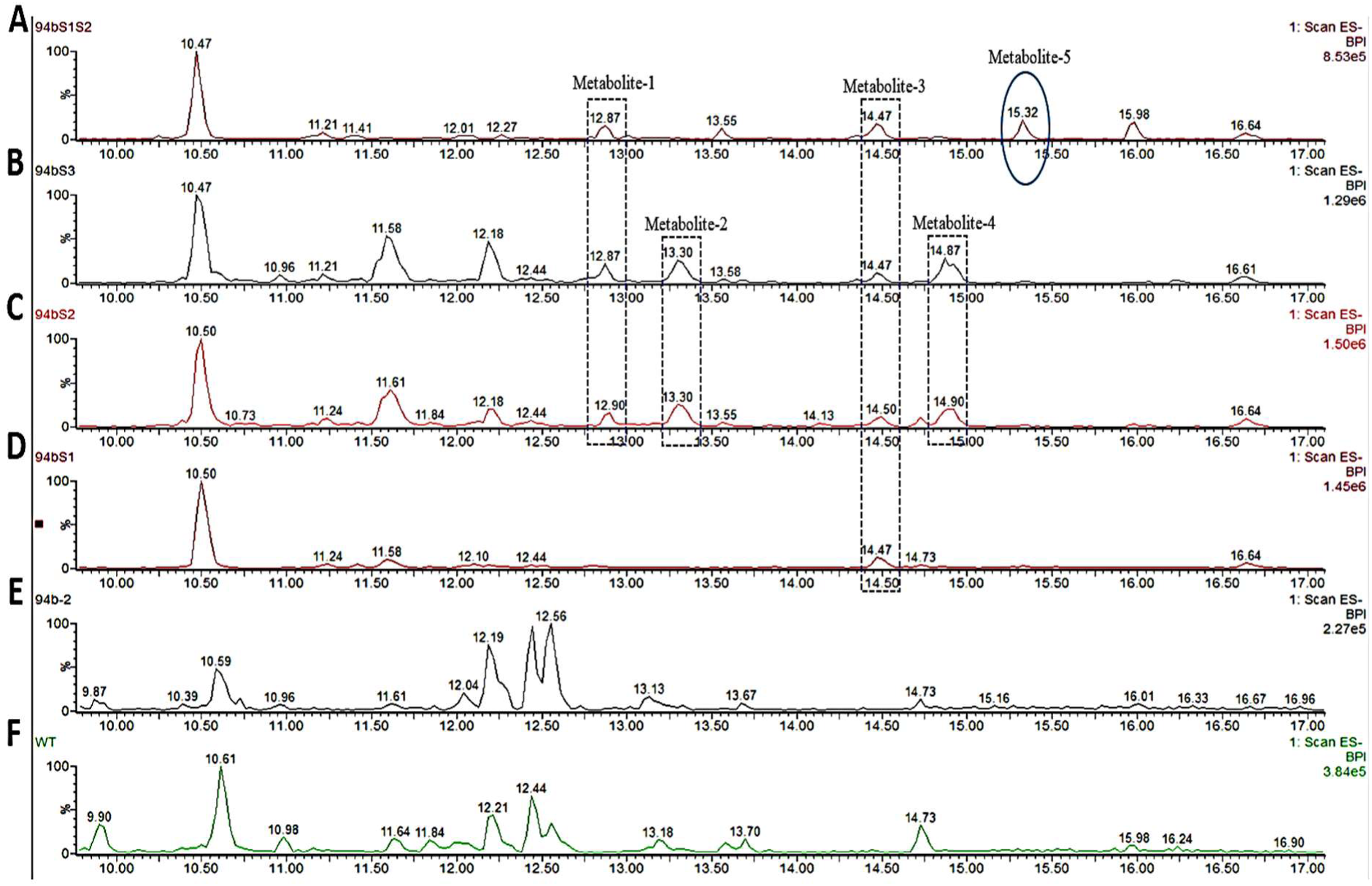
LC-MS comparison of NSAR1 transgenics. **A-** NSAR1-humulene synthase-SDR1-SDR2. **B-** NSAR1-humulene synthase-SDR3. **C-** NSAR1-humulene synthase-SDR2. **D-** NSAR1-humulene synthase-SDR1. **E-** NSAR1-humulene synthase.The mass spectrum was selected from minute 10 to minute 17 to avoid traces overlapping. In total five novel metabolites were detected within this comparison. NSAR1-humulene synthase-SDR1showed the lowest number of novel metabolites compared to NSAR1-humulene synthase-SDR2 and NSAR1-humulene synthase-SDR3.

**Figure 15:**
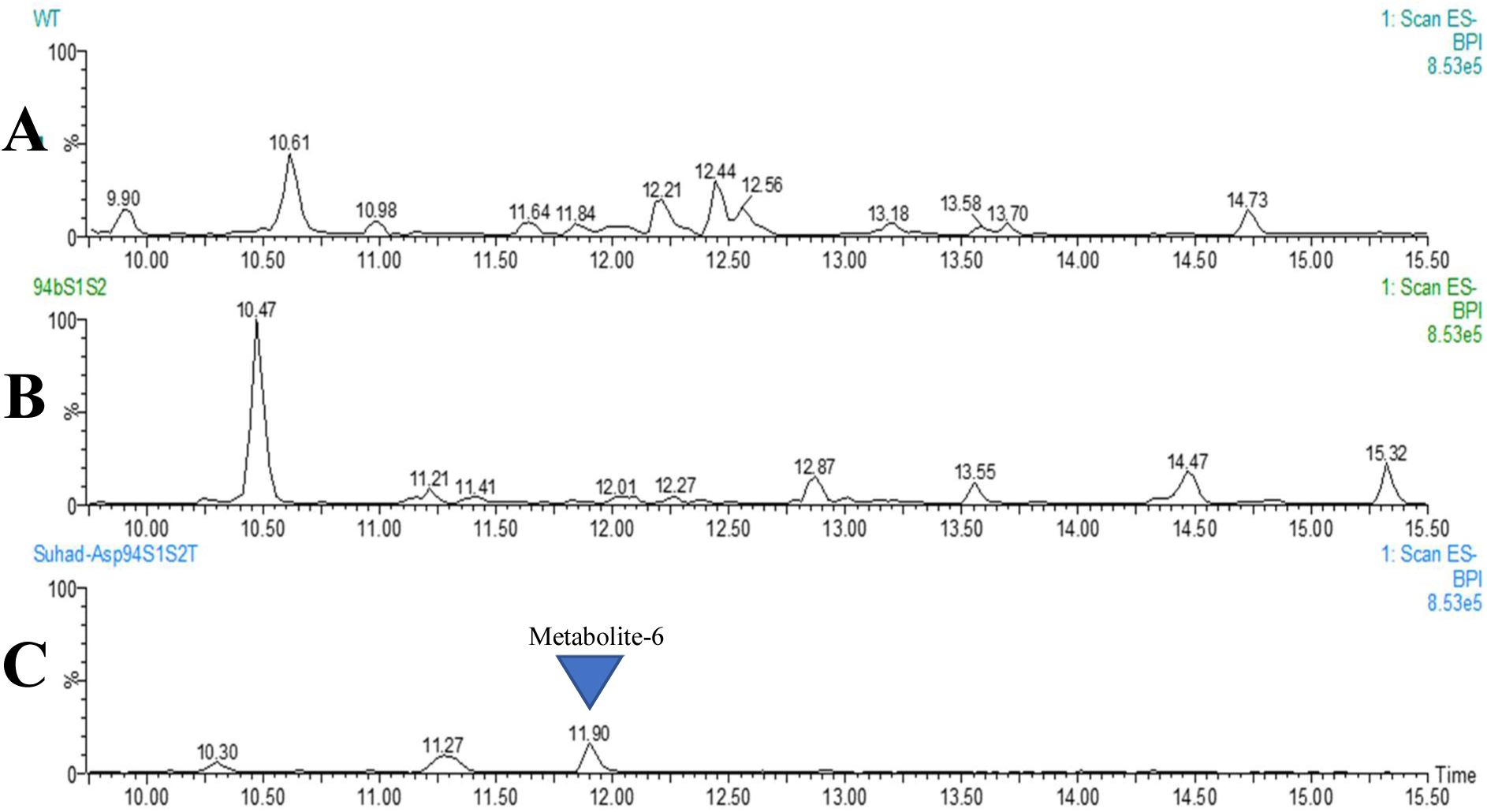
LC-MS comparison of NSAR1 transgenics. **A-** NSAR1. **B-** NSAR1-humulene synthase-SDR1-SDR2. **C-** NSAR1-humulene synthase-SDR1-SDR2- Tyrosinase.

## Discussion

The constant rise in antibiotic resistance has prompted scientists to look for new ways of drug discovery. While mushroom forming fungi reported as a rich repository of structurally diverse natural products, the limited access to their metabolomic and genomic data pose bottleneck to the development of such potential source of antibiotics. In our study we applied and bespoke approaches to investigate the potential of selected fungi as a source of novel antibiotics. Our antagonistic assays verified eight of the nine selected species as potentially antibiotic species, of which the two *Hypholoma* species displayed a wide-range bioactivity, demonstrating their flexibility in their response to different niche conditions and their ability to protect themselves from potential predation (15), which could be potentially important in defeating resistance microbes and worth further investigations.

The direct bioautography bioassay and subsequent chemical purification and spectroscopic analysis, led to the identification of a biologically active aromatic compound 3,5 D. The antibacterial activity test against *B. subtilis* and *S. aureus*, distinguished this metabolite as a potent antibacterial agent. Although, 3, 5 D first isolation was from *Stropharia sp*. by (16), their spectroscopic characterization was poor where only two aromatic chemical shifts were observed in their 1D NMR experiments. Here in we further confirm the presence of the substituent groups (Cl, CH_3_ and COOH) through a full 1D and 2D NMR spectroscopic characterization including 1H, 13C, HMBC, HMQC and NOESY, allowing the first elucidation of 3, 5 D in *Hypholoma* genus and second within fungi kingdom. 3, 5 D is a dichlorinated like natural product, although its biosynthetic pathway is not reported thus far, there is evidence that this compound may be synthesised from benzoate backbones via the shikimate pathway (17-19). Previous literatures reported that such compounds synthesised through shikimate pathway involving the conversion of phenylalanine precursor to benzoic derivatives via sequential condensation, hydroxylation and chlorination reactions. Interestingly, the predicted HfasTerp104 gene cluster includes enzymes that are likely responsible for the synthesis of compound 3, 5 D including benzoic acid reductase-PKS, SDR, glycoside hydrolase and the multifunctional 3-phosphoshikimate-1-carboxyvinyltransferase. 3-phosphoshikimate-1-carboxyvinyltransferase is a multidomain enzyme its main role is to catalyse the conversion of phenylalanine like compounds to cinnamic acid derivatives. Subsequent hydroxylation and reduction on the resulted acid leads to the production of anisaldehyde isomers such as 3, 5 D (20). The biological activity of chlorinated natural products is well documented (21-23) having their putative BGC is a step forward towards the biochemical manipulation of such potent antimicrobials.

Unlike *H. sublateritium, H. fasciculare* genome was assembled into greater number of short contigs. It is likely that our assembling method assembled the resulted gene clusters of *H. fasciculare* into several short contigs rather than one long scaffold as the case of its related *H. sublateritium* genome assembled by JGI. Nevertheless, phylogenetic comparisons and in-depth manual curation with *H. sublateritium* genome sequences guided the annotation of a set of unique biosynthetic gene clusters in both species that awaiting investigation. Most of the 15 putative gene clusters contain key genes encoding terpene synthase and predicted cytochrome P450, short chain dehydrogenase, GMC oxidoreductase, transcription factor, glycoside hydrolase, aromatic ring hydroxylase, pyruvate decarboxylase which represent the main features of fungal SM biosynthetic pathways. Our *in silico* analysis suggest that these clusters are responsible for the production of terpene like compounds, particularly sesquiterpenes and triterpenes (previously reported compounds in *Hypholoma* spp.). A multimodule NRPS core enzyme was also found in contig 29 of *H. fasciculare* genome, subsequent manual curation led to the identification of a putative biosynthetic cluster in both *Hypholoma* (contig 29 of Hfas and scaffold 100 of Hsub), including glycoside hydrolase, polyadenylate-binding protein, AMP-dependent synthetase and ligase, oxidoreductase, acyl-CoA N-acyltransferase. Most often multidomains NRPS are involved in the synthesis of SM with medicinal properties (24) indicative that *Hypholoma* species may produce NRPS based metabolites that yet to be characterised.

We also identified several hybrid clusters that contain two structurally different key enzymes (e.g. a predicted PKS and a terpene synthase). Such biosynthetic clusters are often coordinate the production of meroterpenoid molecules such as melleolide (25). As well as our newly characterised 3,5 D which is a phenolic molecule synthesize through a specific biosynthesis rout involves seven sequential steps. Further annotation of our clusters suggested that the PKS of Hfasterp104 cluster is actually a benzoic acid reductase which apparently produce benzoate precursor. This enzyme along with the multifunctional gene; 3- phosphoshikimate-1-carboxyvinyltransferase (catalyses five sequential steps in shikimate pathway) are the key providers of phenolic compounds biosynthesis in plants (26).

Additionally, our genome investigation identified a biosynthetic cluster (Hfas-Sid14) in contig 14 with putative role in siderophore biosynthesis. Typical biosynthetic enzymes including GMC oxidoreductase, cytochrome P450, glycoside hydrolase and a transcription factor were predicted in this 98 kb cluster (see Figure S4 in supplementary information), indicative that our *Hypholoma* produces siderophore derived metabolites that awaiting exploration.

To support our *in silico* prediction, we attempted gene silencing to down regulate the expression of some selected terpene key enzymes alongside argininosuccinate synthetase gene, where we demonstrated that RNA silencing is an effective tool in knocking down transcript of argininosuccinate gene. Silenced transformants showed variability in their phenotypes represented by slow rate of growth or production of unusual form of mycelia, demonstrating successful triggering of RNAi silencing in *H. fasciculare*. The observed phenotypic variation is congruent with silencing work in basidiomycetes (27, 28). Extended silencing work on sesquiterpene synthases revealed a crossed role of such enzymes in the synthesis of *H. fasciculare* secondary metabolites, where production of several SM was abolished as a result of silencing one enzyme. For example, silencing HfasTerp84b resulted in the absence of most metabolites that observed in the WT. This promiscuous feature of sesquiterpene synthase of basidiomycetes, was previously reported by (10 and 11), where independent heterologous expression of a group of sesquiterpene synthases from two different basidiomycetes; *O. olearius* and *C. cinerea* in *E. coli*, resulted in several chemically distinguished sesquiterpene molecules for each enzyme. As well as, we expressed our sesquiterpene synthase alongside above mentioned enzymes in *A. oryzae*. Interestingly the GC-MS analysis of *A. oryzae* transformants harbouring some of (10 and 11) enzymes confirmed the fact that *A. oryzae* is a feasible tool in linking SM to their specific enzymes and that its metabolomic system has no effect on the final product of sesquiterpene enzymes, as most major products of (10 and 11) were replicated here. The miss observation of the remaining compounds in the other *Aspergillus* transgenics, is likely due to the use of different isolation method, as compounds detection is largely influenced by the isolation method and fiber used during the characterisation process (29).

The detection of humulene in the GC-MS analysis of HfasTerp94 motivated us to further express the remaining biosynthetic genes within this cluster in *A. oryzae*. In this cluster three SDR genes were predicted alongside other distinctive genes including zinc carboxypeptidase, tyrosinase. SDR genes belong to a large family of oxidoreductases that have a key role in oxido-reductase reactions through changing the oxidation number of molecules (30). Bearing this in mind we hypothesised that co-expressing of SDR gene with humulene synthase would cause the production of an oxidized humulene derivatives.

Different combinations of genes (dual and triple gene vector) were therefore transformed into NSAR1 to potentially reveal the function of each enzyme. Positive *Aspergillus* transformants were chemically investigated leading to the detection of several new peaks, suggestive that SDR and tyrosinase protein sequences were correctly predicted and they can further modify humulene backbone.

Although most of detected metabolites showed typical mass fragmentation of sesquiterpene like compounds (11), further characterization of compounds structure (e.g. NMR) is needed to precisely link produced molecules to their corresponding genes.

## Conclusion

Mushroom forming fungi meet the demand in antibiotic industry to produce molecules with novel antimicrobial mechanism. However, a small number of these fungi have been fully explored at the genetic and chemical level. To address problems associated with *H. fasciculare* SM production -low yield and growth difficulties- in depth sequences analysis of *H. fasciculare* genome carried out, leading to the characterization of many putative biosynthetic gene clusters that are likely involve in the production of previously reported compounds in this fungus. When we experimentally tested our *in silico* genome prediction via RNAi mediated and heterologous expression in *A. oryzae*, we further demonstrated that RNAi silencing is still an efficient tool in triggering the downregulation of gene expression and could facilitate other genetic manipulation in *H. fasciculare*, and that sesquiterpene synthase are promiscuous enzymes. The study also permitted the chemical characterization of a biologically active metabolite (3,5 D), as well as the *in silico* prediction of its associated biosynthetic gene cluster, offering the opportunity to further manipulate its production in *Hypholoma* species and other rot decaying fungi, either homologously or heterologously, and demonstrates that *Hypholoma’s* secondary metabolites warrant further investigations, providing a remarkable source for the pharmaceutical industry.

## Acknowledgments

SA-S was financially supported by Ministry of Higher Education and Scientific research-IRAQ. Research conducted at the University of Bristol, UK. The authors thank Dr Fabrizio Alberti, Dr Claudio Greco, and Dr Colin Lazarus for their assistance throughout the study.

## Notes

### Competing Interest Statement

The authors have declared no competing interest.

