## supplementary information for "Further biochemical profiling of *Hypholoma fasciculare* metabolome reveals its chemogenetic diversity" (from another version of this preprint: version 1, 2020-05-31)

Figure S1: Represent the so far characterized compounds from *H. fasciculare*.

Table S1: Selected experimentally characterised terpene cyclase, polyketide synthase, non-ribosomal peptides and their sequences details used in Hypholoma genome comparison.

Table S2: Predicted SM of Hypholoma spp. using antiSMASH and Local BLAST

Figure S2 Bioassay test to evaluate the antimicrobial activity of *Paxillus involutus*.

Figure S3: Bioassay test to evaluate the antimicrobial activity of *Coniophora puteana.*

Figure S4: Bioassay test to evaluate the antimicrobial activity of *Fomitiporia mediterranea.*

Figure S5: Bioassay test to evaluate the antimicrobial activity of *Postia placenta*.

Figure S6: Bioassay test to evaluate the antimicrobial activity of *Gloeophyllum trabeum*.

Figure S7: Bioassay test to evaluate the antimicrobial activity of *Laccaria bicolor.*

Figure S8: Bioassay test to evaluate the antimicrobial activity of *Suillus luteus.*

Figure S9: TLC plate developed in semi polar system and visualized under UV 320nm showing separated compounds of *H. fasciculare*.

Figure S10: TLC plate developed in non-polar system and visualized under UV 320nm showing separated compounds of *H. fasciculare*.

Figure S11: TLC plate developed in semi polar system and visualized under UV 320nm showing separated compounds of *H. sublateritium*.

Figure S12: TLC plate developed in non-polar system and visualized under UV 320nm showing separated compounds of *H. sublateritium*.

Figure S13: Direct bioautography of *H. fasciculare* crude extracts showing the antimicrobial activity of different crude extracts developed in polar solvents.

Figure S14: Direct bioautography of *H. fasciculare* crude extracts showing the antimicrobial activity of different crude extracts developed in semi polar solvents.

Figure S15: Direct bioautography of *H. fasciculare* crude extracts showing the antimicrobial activity of different crude extracts developed in non-polar solvents.

Figure S16: LC-MS chromatograms electrospray mass negative mode (ES-).

Figure S17: ^1^H NMR (125 MHz) spectrum of 3, 5-dichloro-4-methozybenzoic acid.

Figure S18: ^13^C NMR (500 MHz) spectrum of 3, 5-dichloro-4-methozybenzoic acid.

Figure S19: COSY (500 MHz) spectrum of 3, 5-dichloro-4-methozybenzoic acid.

Figure S20: HSQC (500 MHz) spectrum of 3, 5-dichloro-4-methozybenzoic acid.

Figure S21: HMBC (500 MHz) spectrum of 3, 5-dichloro-4-methozybenzoic acid.

Figure S22: Predicted gene cluster of Hfas-terp105 and the homologous cluster (Hsub-terp83) in *H. sublateritium*

Figure S23: Predicted gene cluster of *H. sublateritium* terpene synthases (30, 113, 38) and their homologous in *H. fasciculare*.

Figure S24: Predicted gene cluster of *H. sublateritium* terpene synthases (11A, 205, 28) and their homologous in *H. fasciculare.*

Figure S25: Predicted gene cluster of *H. sublateritium* terpene synthases (133, 167) and their homologous in *H. fasciculare*.

Figure S26: Predicted gene cluster of *H. sublateritium* terpene synthase 11B and the homologous cluster in *H. fasciculare*

Figure S27: Predicted gene cluster of *H. sublateritium* terpene synthase 99 and the homologous cluster in *H. fasciculare*.

Figure S28: Predicted gene cluster of *H. fasciculare* and *H. sublateritium* PKS.

Figure S29: Predicted gene cluster of *H. fasciculare* and *H. sublateritium* NRPS.

Figure S30: Predicted gene cluster of *H. sublateritium* SidA and the homologous cluster in *H. fasciculare*.

Bioassay preparation

References

**Figure S1: Represent previously characterized compounds from *H. fasciculare* between 1967-2019.**

Molecular weight and chemical formula were illustrated in this table.

A- Fascicularone isomers, B- Fasciculic acid isomers, C- Fasciculol isomers, D- p**- anisaldehyde isomers,** E- **Naematolin isomers and** F- **Ergosterol isomers**

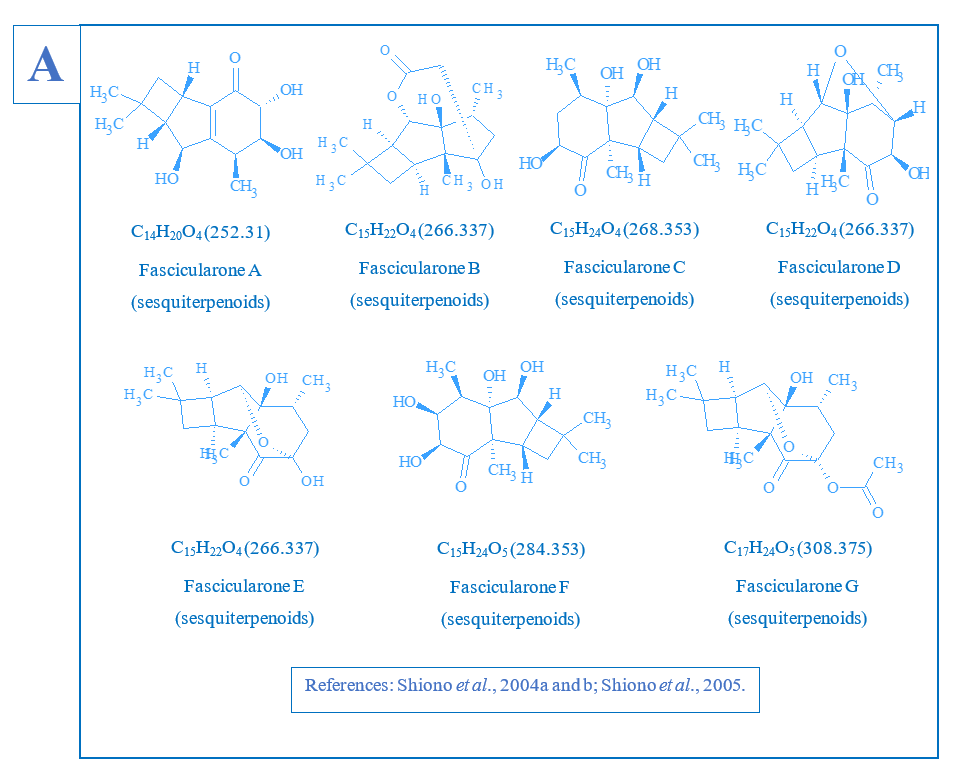

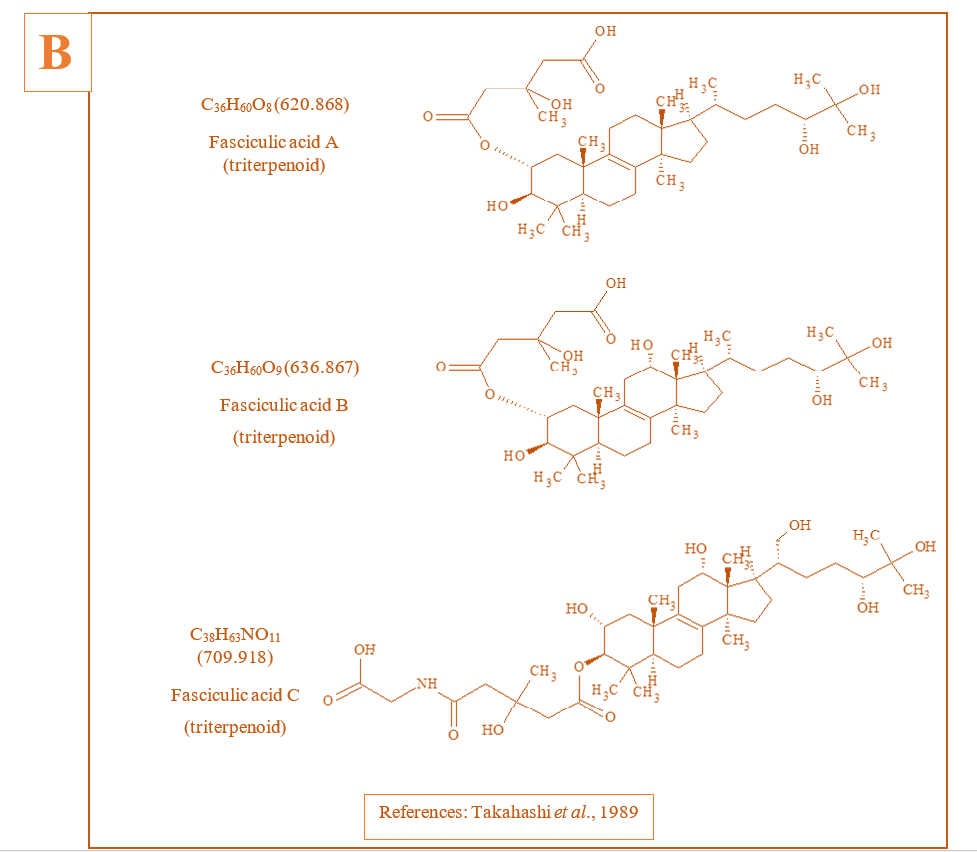

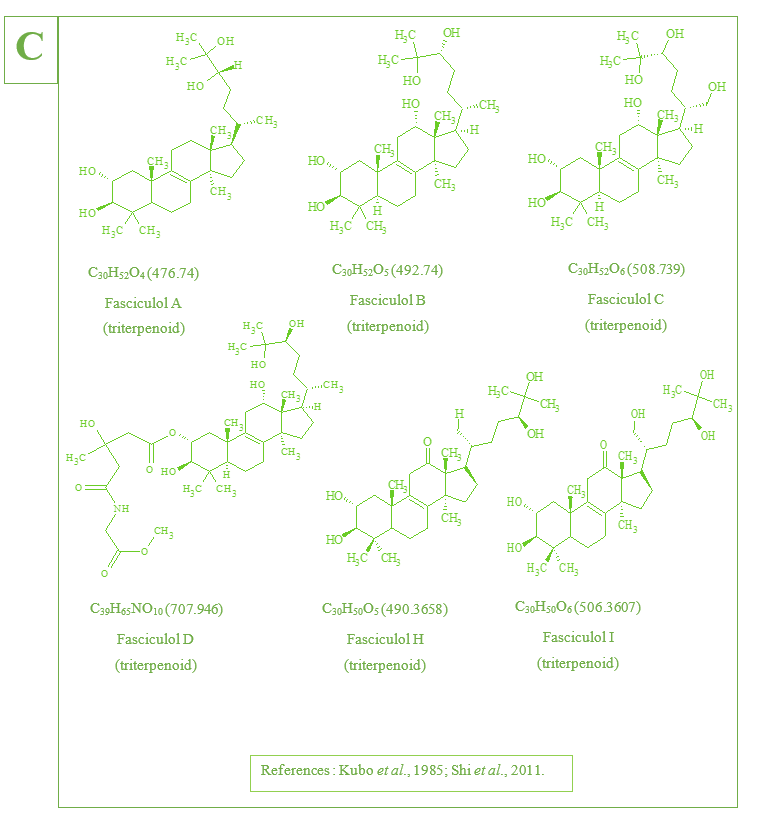

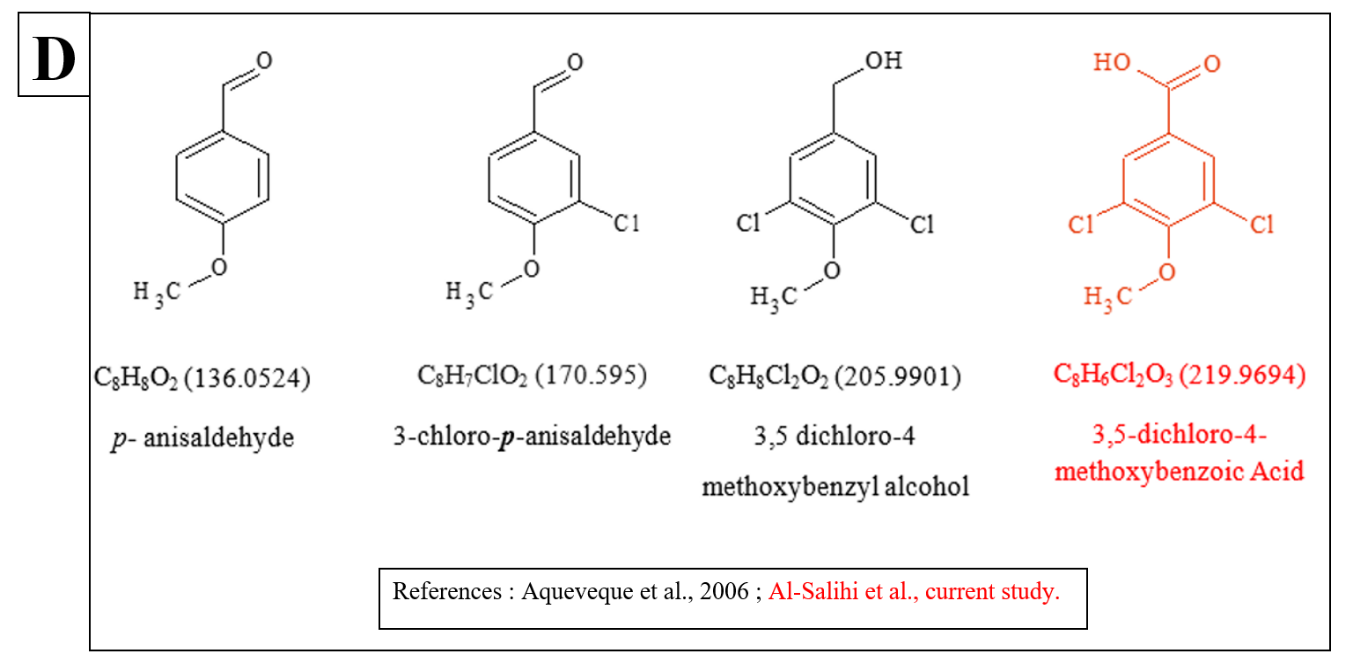

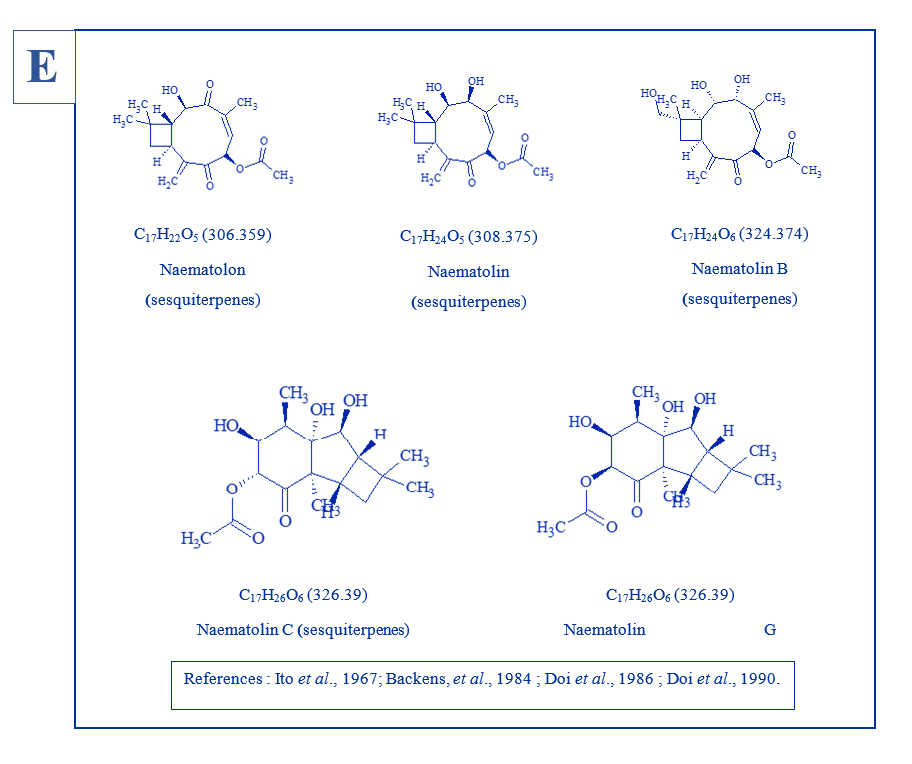

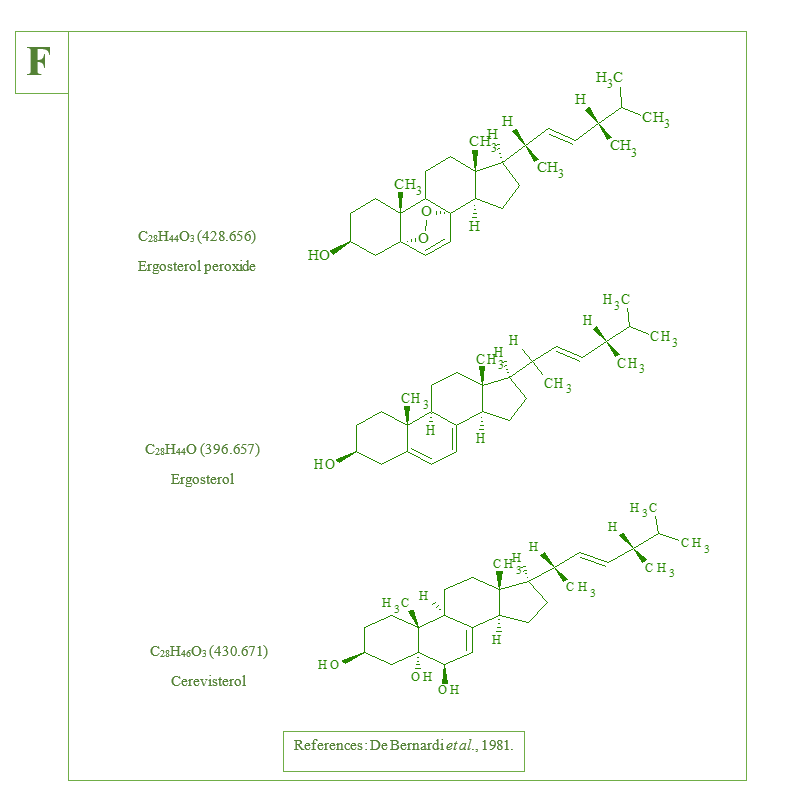

Table S1: Selected experimentally characterised terpene cyclase, polyketide synthase, non-ribosomal peptides and their sequences details used in Hypholoma genome comparison.

| SM type | | SM cyclase or synthetase | Enzyme feature | NCBI or JGI protein ID | Organism | References |
| --- | --- | --- | --- | --- | --- | --- |
| 1 | TC | Ompol1\| (1) | 1,10 cyclization of E, E, F-PP | 1311 | Omphalotus olearius | Wawrzyn et al., 2012 |
|  |  | Copci1\| (3) |  | 14587 | Coprinus cinereus | Agger et al., 2009 |
|  |  | Stehi1 | 1,10 cyclization of 3R-NPP | 128017 | Stereum hirsutum | Quin et al., 2013 |
|  |  | Ompol1\| (4) |  | 1447 | Omphalotus olearius | Wawrzyn et al., 2012 |
|  |  | Stehi1\| | 1,11 cyclization of E,E,FPP | 64702 | Stereum hirsutum | Quin et al., 2013 |
|  |  | Ompol1\| (7) |  | 2271 | Omphalotus olearius | Wawrzyn et al., 2012 |
|  |  | Ompol1\| (9) | 1,6 cyclization of 3R/S-NPP | 3258 | Omphalotus olearius | Wawrzyn et al., 2012 |
|  |  | Copci1\| (6) |  | 20268 | Coprinus cinereus | Quin et al., 2013 |
| 2 | PKS | LDKS | Lovastatin diketide synthase | Q9Y7D5 | *Aspergillus terreus* | Meehan et al., 2010 |
|  |  | PKSA | Sterigmatocystin polyketide synthase | Q12053.1 | *Aspergillus parasiticus* | Feng and Leonard, 1995 |
|  |  | hybrid PKS-NRPs | Beta-Keto acyl synthase | Q8J0F7 | *Penicillium citrinum* | Abe et al., 2002 |
|  |  | MSAS | 6-Methylsalicyclic acid synthase | AAC49814.1 | *Aspergillus terreus* | Fujii et al., 1996 |
|  |  | ArmB |  | JQ801748 | Armillaria mellea |  |
|  |  | PKS1 |  | 169950 | Coniophora puteana |  |
|  |  | PKS1 |  | 116317 | Gloeophyllum trabeum |  |
| 3 | NRPS | AMT | AM-toxin synthetase | AAF01762.1 | *Alternaria alternata* | Johnson et al., 2000 |
|  |  | SidD | SidD | 748662.1 | *Aspergillus fumigatus* | Nierman et al., 2005 |

Table S2: Predicted SM of Hypholoma spp. using antiSMASH and Local BLAST

| Cluster type | Predicted SM gene clusters using antiSMASH | |
| --- | --- | --- |
|  | ***H. fasciculare*** | ***H. sublateritium*** |
| Terpene | Contigs: 70, 85, 94, 105, 173, 179 | Scaffolds: 11a, 30, 43, 133 |
| Siderophore | Contig 14 | Scaffold 11 |
| NRPS | Contig 43 | Scaffold 7 |
| T1pks | Contig 221 | Scaffold: 12, 53 |
| Predicted SM gene clusters using Local BLAST | | |
| Terpene | Contigs: 10, 147, 266, 339, 378, 28 | Scaffolds: 11b, 28, 38, 81, 113, 116, 167, 205 |
| PKS | Hfas85 | Hsub116 |
| NRPS | Hfas29 | Hsub99 |

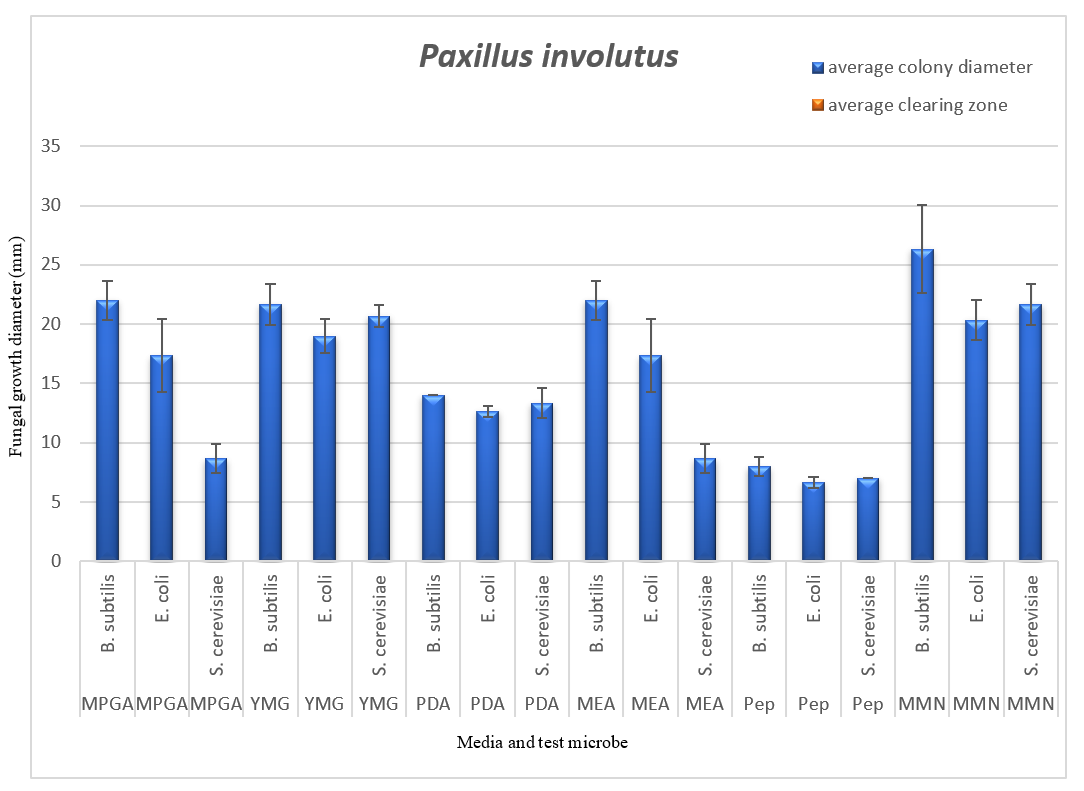

Figure S2: Bioassay test to evaluate the antimicrobial activity of *P. involutus* growing on different media against *B. subtilis*, *E. coli* and *S. cerevisiae*. MPGA = malt peptone glucose agar, YMG = yeast extract malt, PDA = potato dextrose agar, MEA = malt extract agar, Pep = peptone agar, MMN = minimum melin norkans agar. Error bars indicates the standard deviation of three technical replicates measurement for both fungal colony diameter (column in blue) and inhibition zone diameter (column in red).

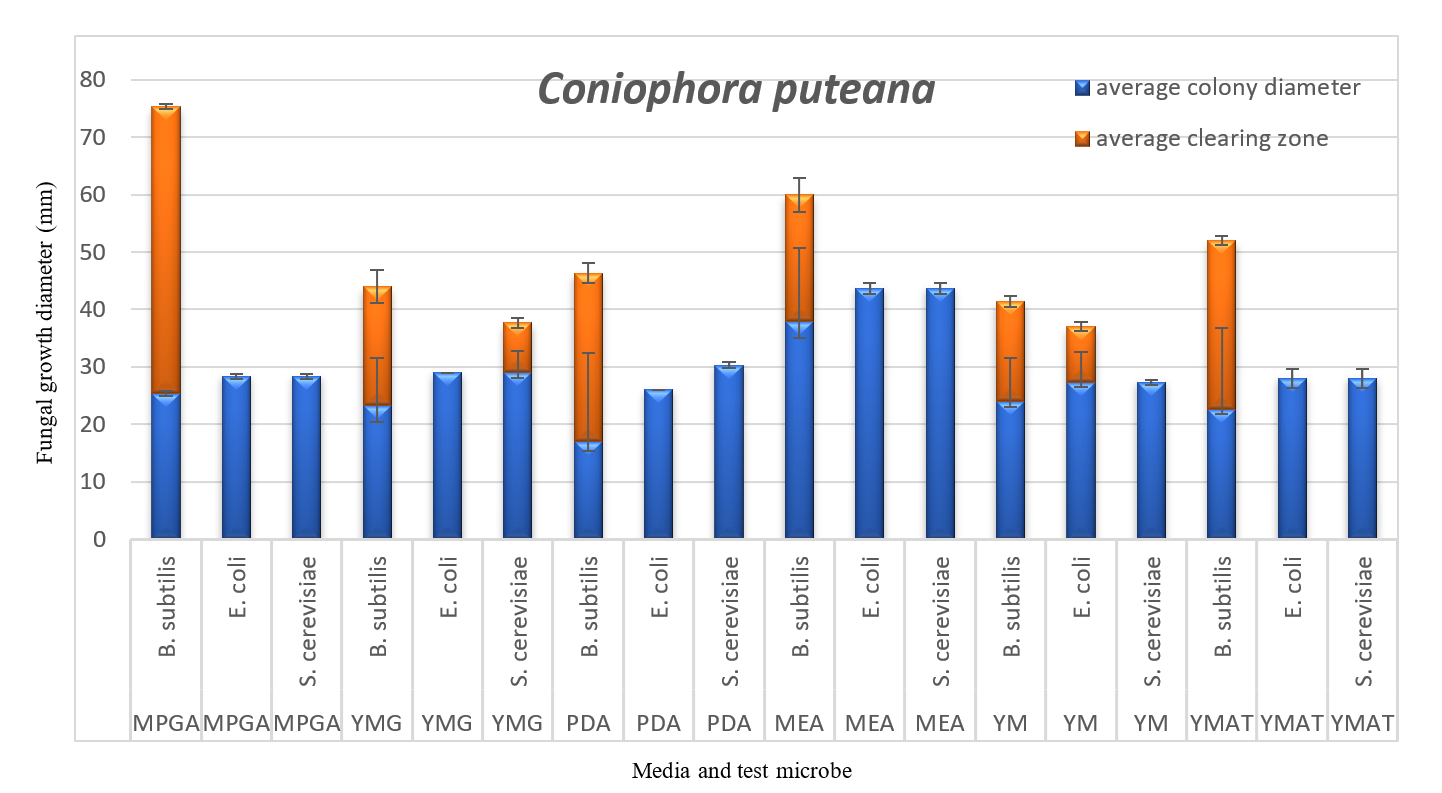

Figure S3: Bioassay test to evaluate the antimicrobial activity of *C. puteana* growing on different media against *B. subtilis*, *E. coli* and *S. cerevisiae*. MPGA = malt peptone glucose agar, YMG = yeast extract malt, PDA = potato dextrose agar, MEA = malt extract agar, Pep = peptone agar, YM = yeast malt extract, YMAT = yeast malt dextrose peptone agar. Error bars indicates the standard deviation of three technical replicates measurement for both fungal colony diameter (column in blue) and inhibition zone diameter (column in red).

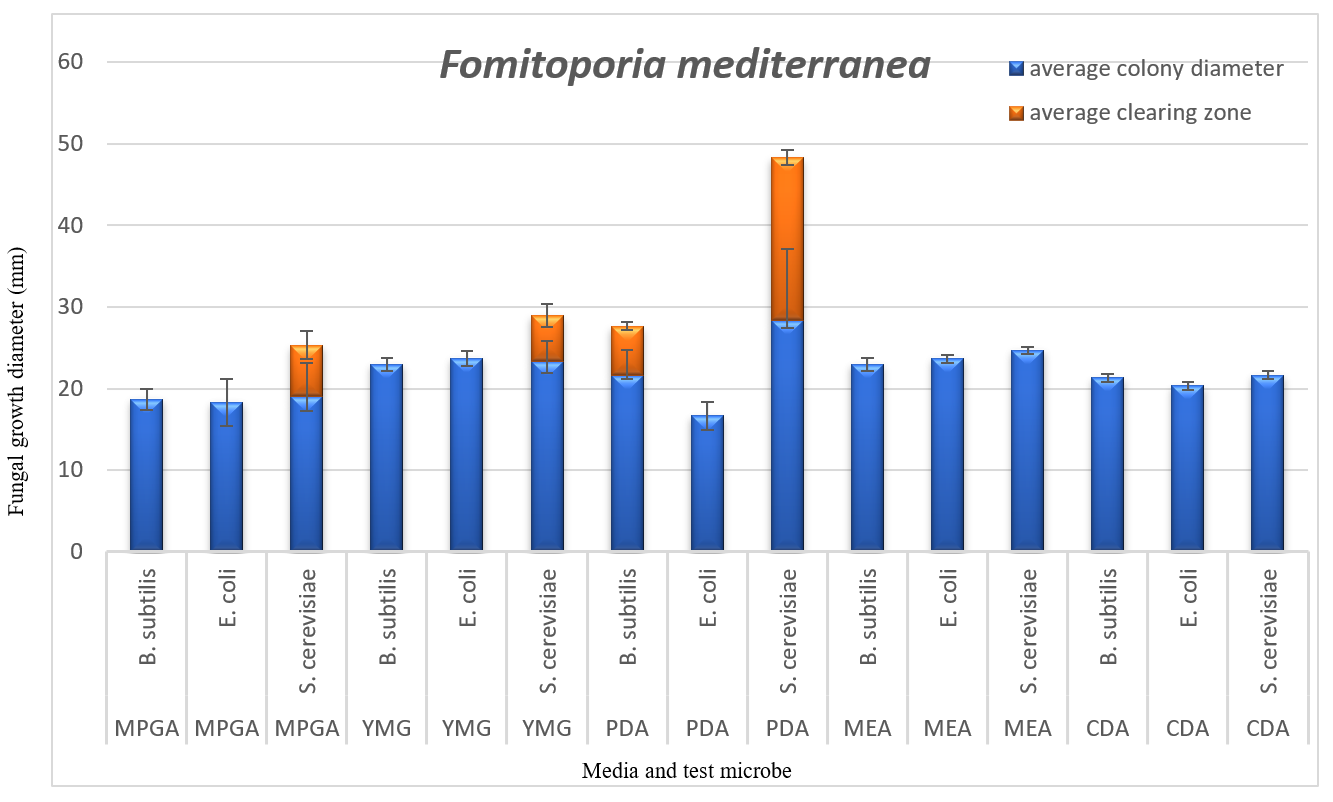

Figure S4: Bioassay test to evaluate the antimicrobial activity of *F. mediterranea* growing on different media against *B. subtilis*, *E. coli* and *S. cerevisiae*. MPGA = malt peptone glucose agar, YMG = yeast extract malt, PDA = potato dextrose agar, MEA = malt extract agar, CDA = Czapek Dox agar. Error bars indicates the standard deviation of three technical replicates measurement for both fungal colony diameter (column in blue) and inhibition zone diameter (column in red).

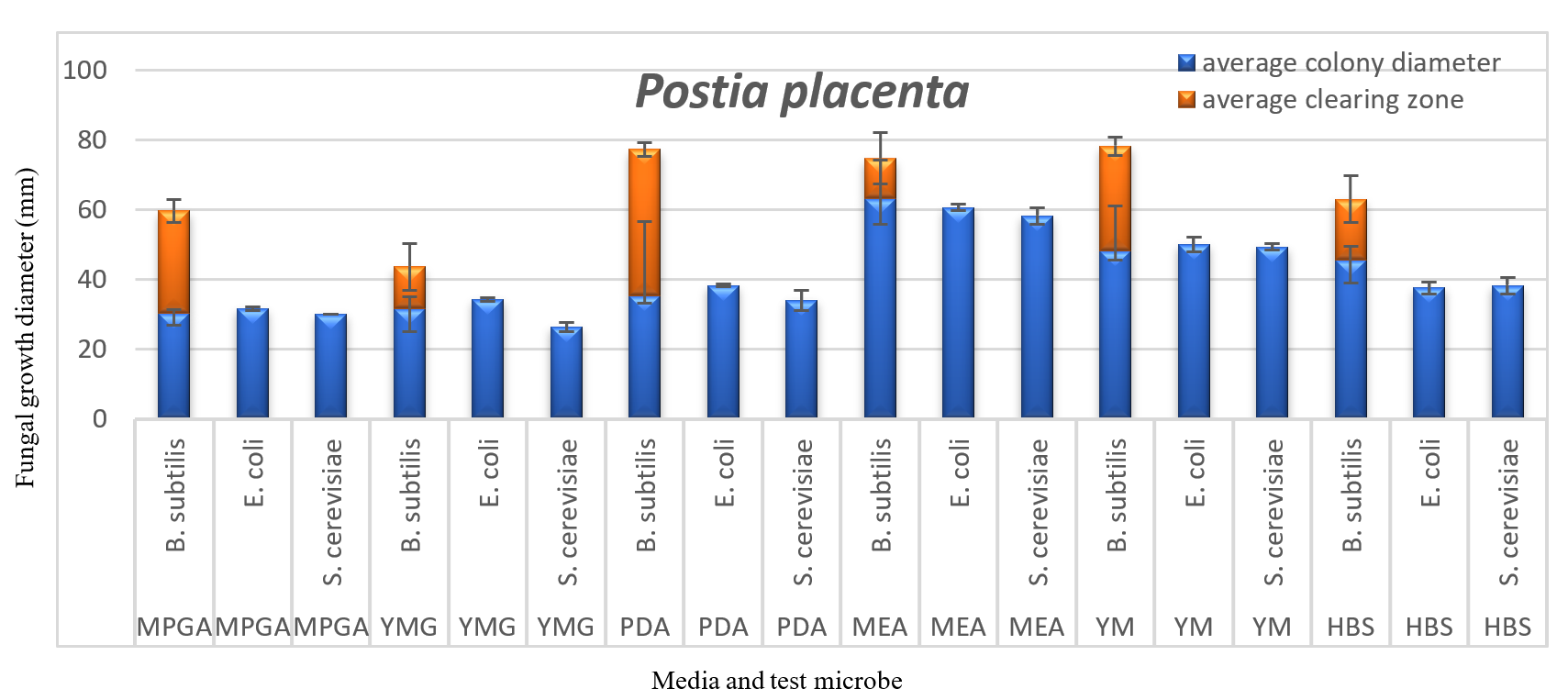

Figure S5: Bioassay test to evaluate the antimicrobial activity of *P. placenta* growing on different media against *B. subtilis*, *E. coli* and *S. cerevisiae*. MPGA = malt peptone glucose agar, YMG = yeast extract malt, PDA = potato dextrose agar, MEA = malt extract agar, YM = yeast malt agar, HBS = highly’s basal salt agar. Error bars indicates the standard deviation of three technical replicates measurement for both fungal colony diameter (column in blue) and inhibition zone diameter (column in red).

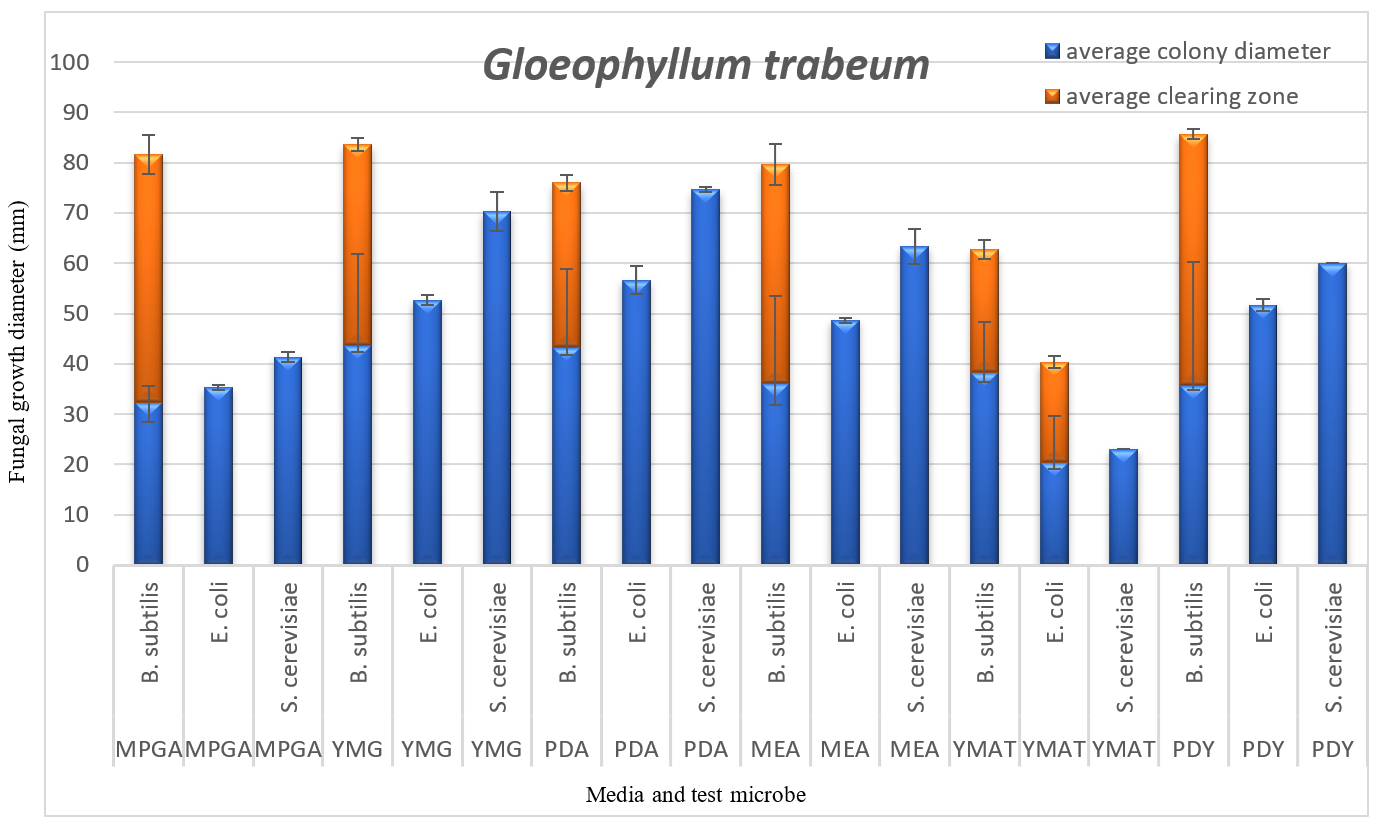

Figure S6: Bioassay test to evaluate the antimicrobial activity of *G. trabeum* growing on different media against *B. subtilis*, *E. coli* and *S. cerevisiae*. MPGA = malt peptone glucose agar, YMG = yeast extract malt, PDA = potato dextrose agar, MEA = malt extract agar, YMAT = yeast malt dextrose peptone agar, PDY = potato dextrose yeast extract. Error bars indicates the standard deviation of three technical replicates measurement for both fungal colony diameter (column in blue) and inhibition zone diameter (column in red).

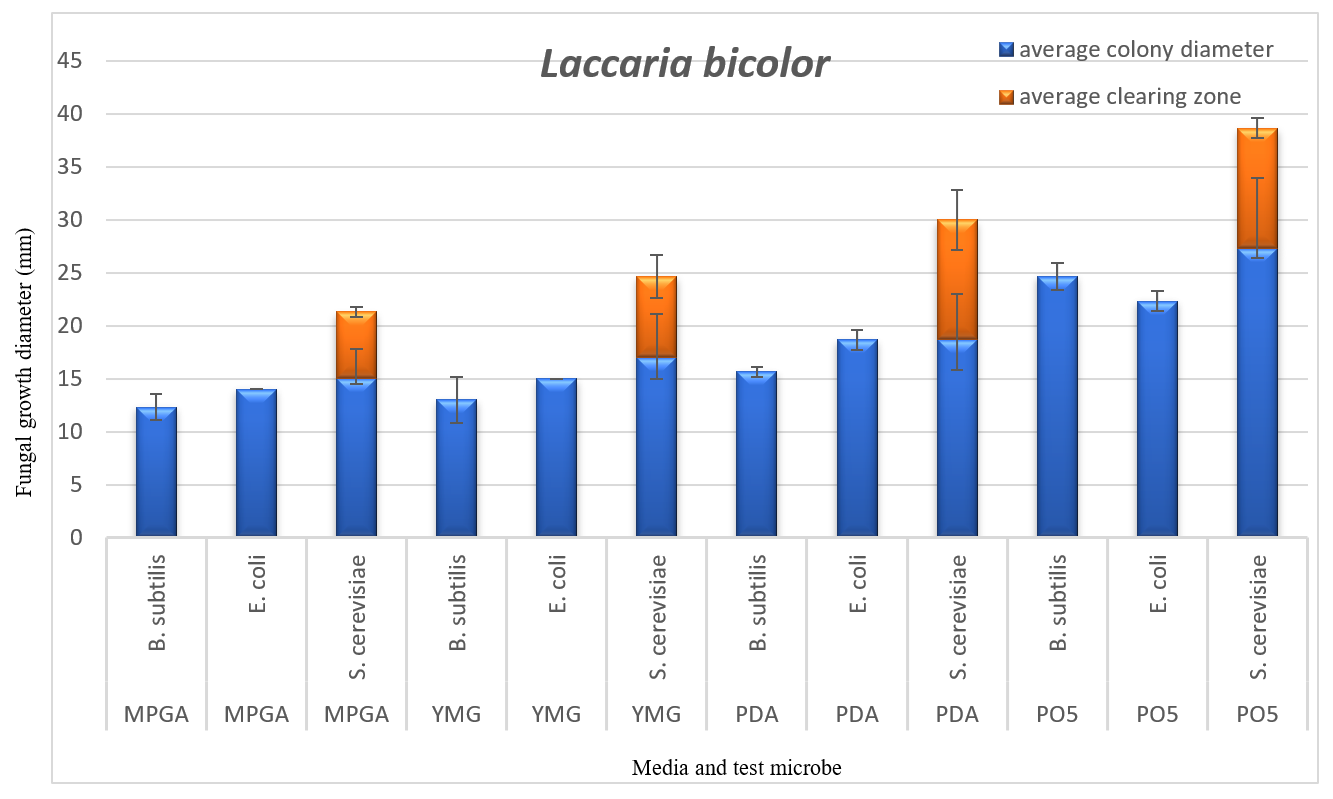

Figure S7: Bioassay test to evaluate the antimicrobial activity of *L. bicolor* growing on different media against *B. subtilis*, *E. coli* and *S. cerevisiae*. MPGA = malt peptone glucose agar, YMG = yeast extract malt, PDA = potato dextrose agar, PO5 = Pachlewski modified agar. Error bars indicates the standard deviation of three technical replicates measurement for both fungal colony diameter (column in blue) and inhibition zone diameter (column in red).

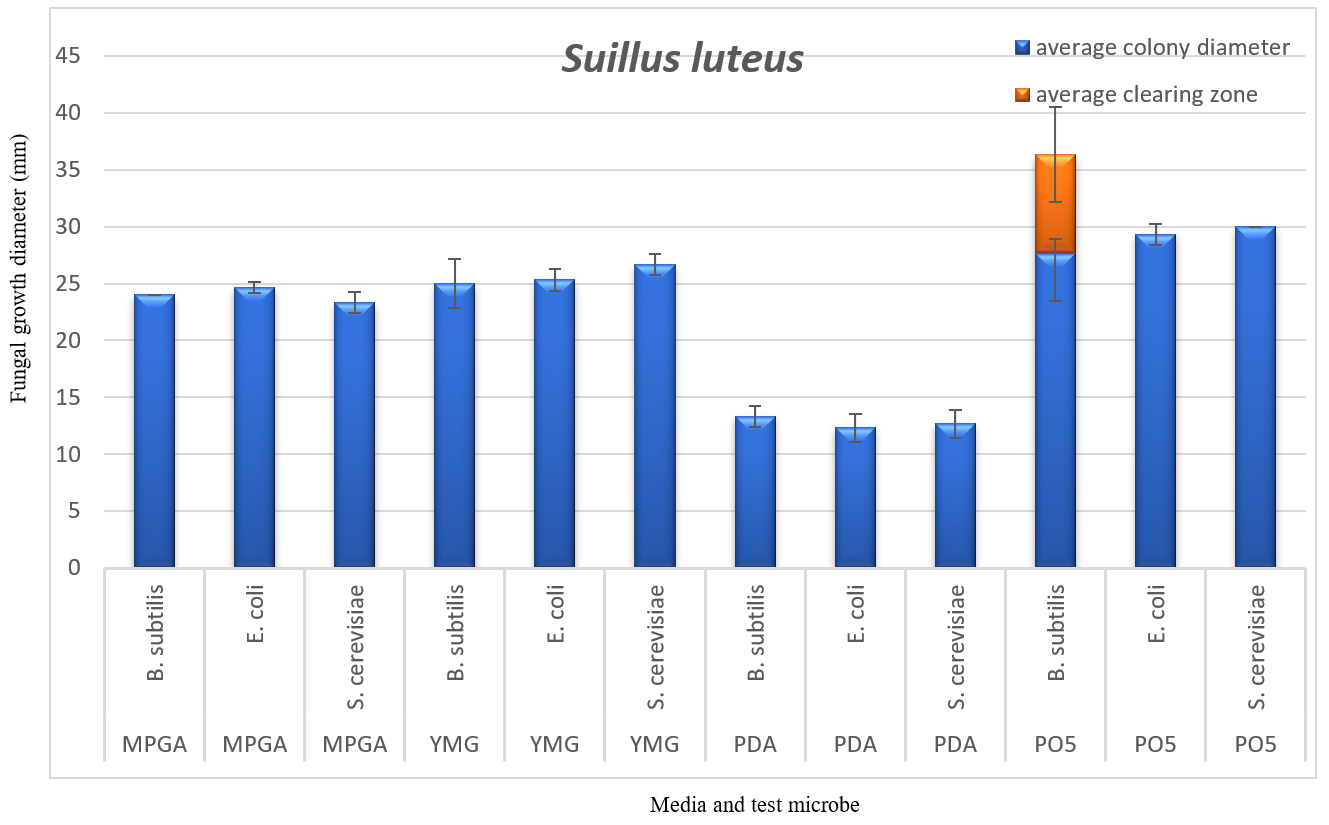

Figure S8: Bioassay test to evaluate the antimicrobial activity of *S. luteus* growing on different media against *B. subtilis*, *E. coli* and *S. cerevisiae*. MPGA = malt peptone glucose agar, YMG = yeast extract malt, PDA = potato dextrose agar, PO5 = Pachlewski modified agar. Error bars indicates the standard deviation of three technical replicates measurement for both fungal colony diameter (column in blue) and inhibition zone diameter (column in red).

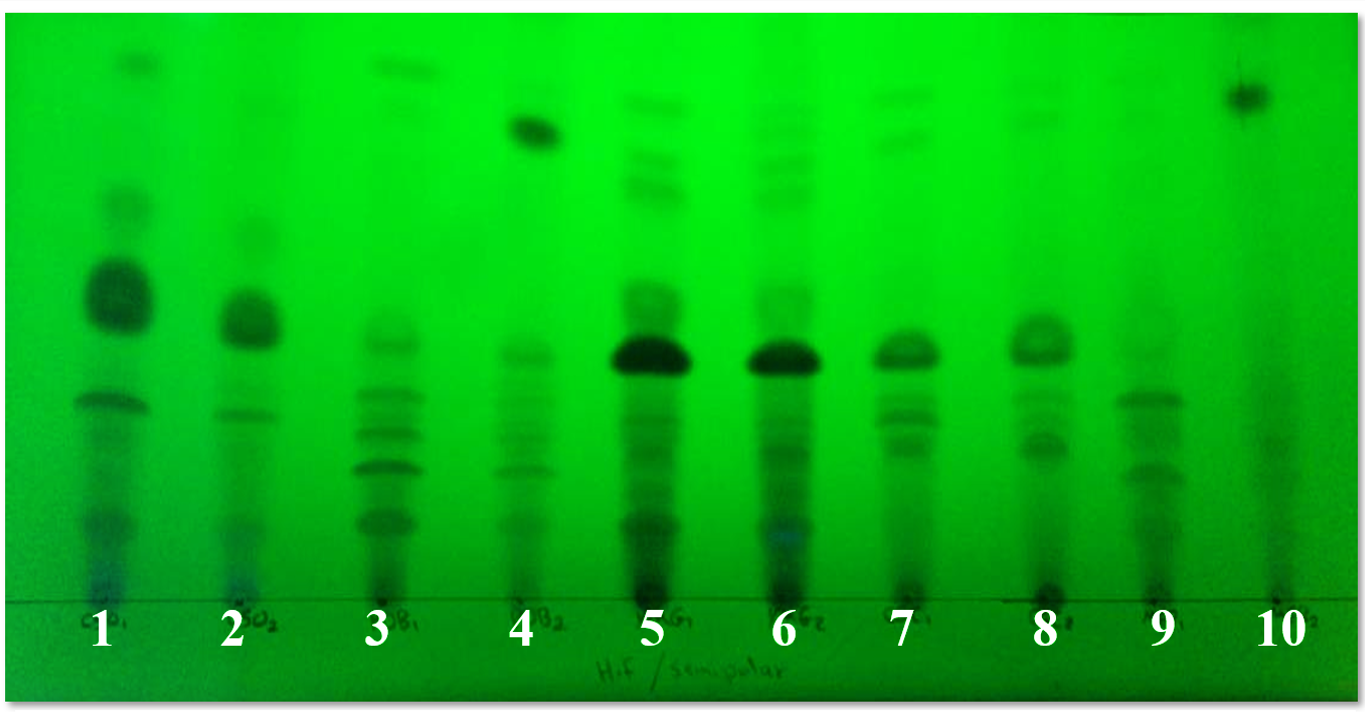

Figure S9: TLC plates developed in semi-polar system and visualized under UV 320 nm showing separated compounds of *H. fasciculare*. 25 µl of 10 mg/ml^-1^ of crude extract of *H. fasciculare* from five different media were spotted.

1 = CSO supernatant extract, 2 = CSO mycelial extract, 3 = PDB supernatant extract, 4 = PDB mycelial extract, 5 = YMG supernatant extract, 6 = YMG mycelial extract, 7 = CGC supernatant extract, 8 = CGC mycelial extract, 9 = MEB supernatant extract, 10 = MEB mycelial extract.

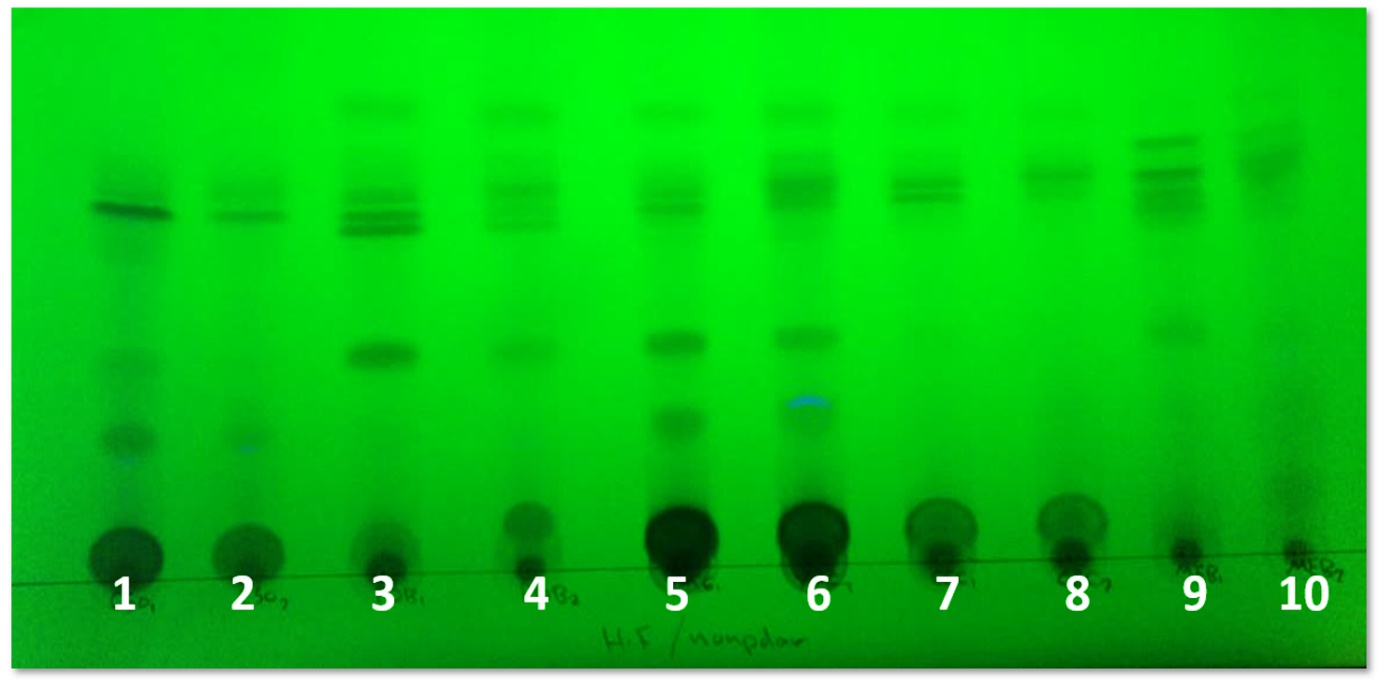

Figure S10: TLC plates developed in non-polar system and visualized under UV 320 nm showing separated compounds of *H. fasciculare*. 25 µl of 10 mg/ml^-1^ of crude extract of *H. fasciculare* from five different media were spotted.

1 = CSO supernatant extract, 2 = CSO mycelial extract, 3 = PDB supernatant extract, 4 = PDB mycelial extract, 5 = YMG supernatant extract, 6 = YMG mycelial extract, 7 = CGC supernatant extract, 8 = CGC mycelial extract, 9 = MEB supernatant extract, 10 = MEB mycelial extract.

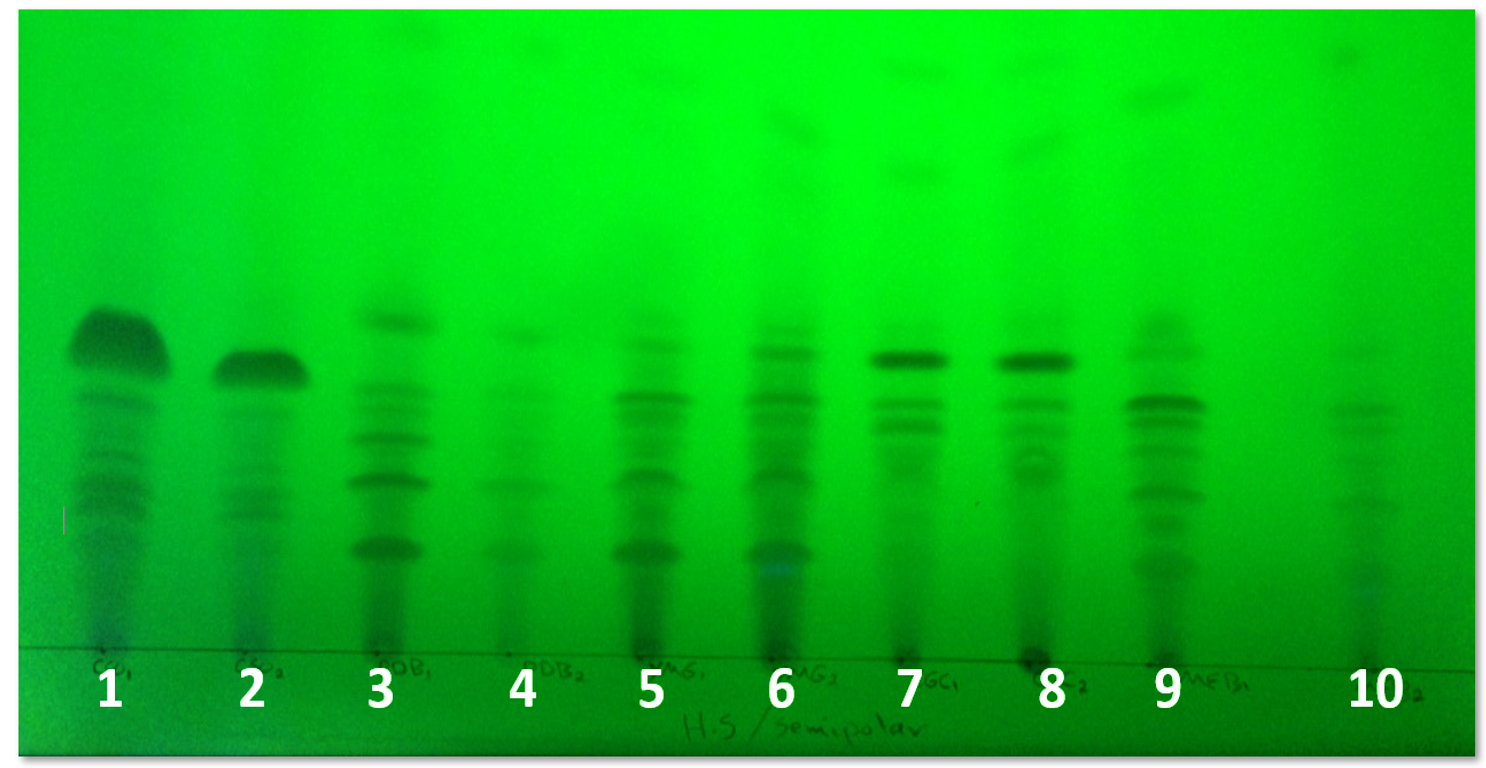

Figure S11: TLC plates developed in semi-polar system and visualized under UV 320 nm showing separated compounds of *H. sublateritium*. 25 µl of 10 mg/ml^-1^ of crude extract of *H. fasciculare* from five different media were spotted.

1 = CSO supernatant extract, 2 = CSO mycelial extract, 3 = PDB supernatant extract, 4 = PDB mycelial extract, 5 = YMG supernatant extract, 6 = YMG mycelial extract, 7 = CGC supernatant extract, 8 = CGC mycelial extract, 9 = MEB supernatant extract, 10 = MEB mycelial extract.

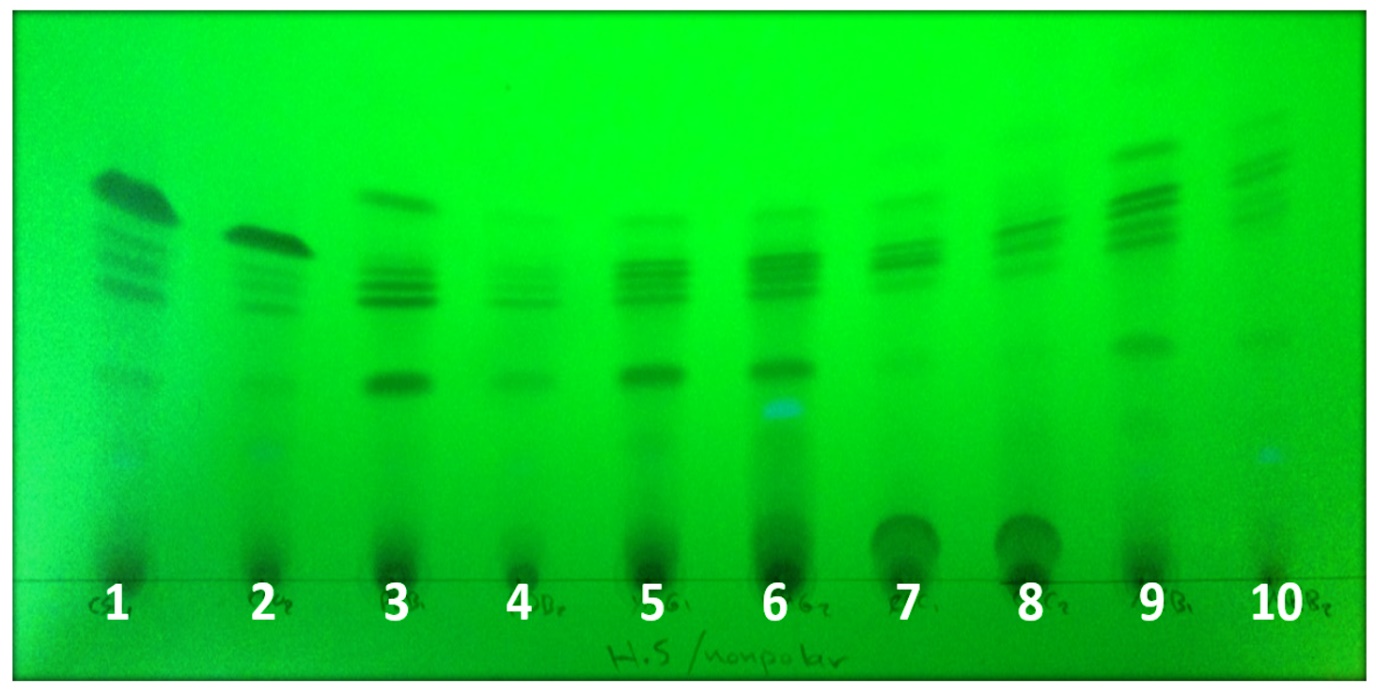

Figure S12: TLC plates developed in non-polar system and visualized under UV 320 nm showing separated compounds of *H. sublateritium*. 25 µl of 10 mg/ml^-1^ of crude extract of *H. fasciculare* from five different media were spotted.

1 = CSO supernatant extract, 2 = CSO mycelial extract, 3 = PDB supernatant extract, 4 = PDB mycelial extract, 5 = YMG supernatant extract, 6 = YMG mycelial extract, 7 = CGC supernatant extract, 8 = CGC mycelial extract, 9 = MEB supernatant extract, 10 = MEB mycelial extract.

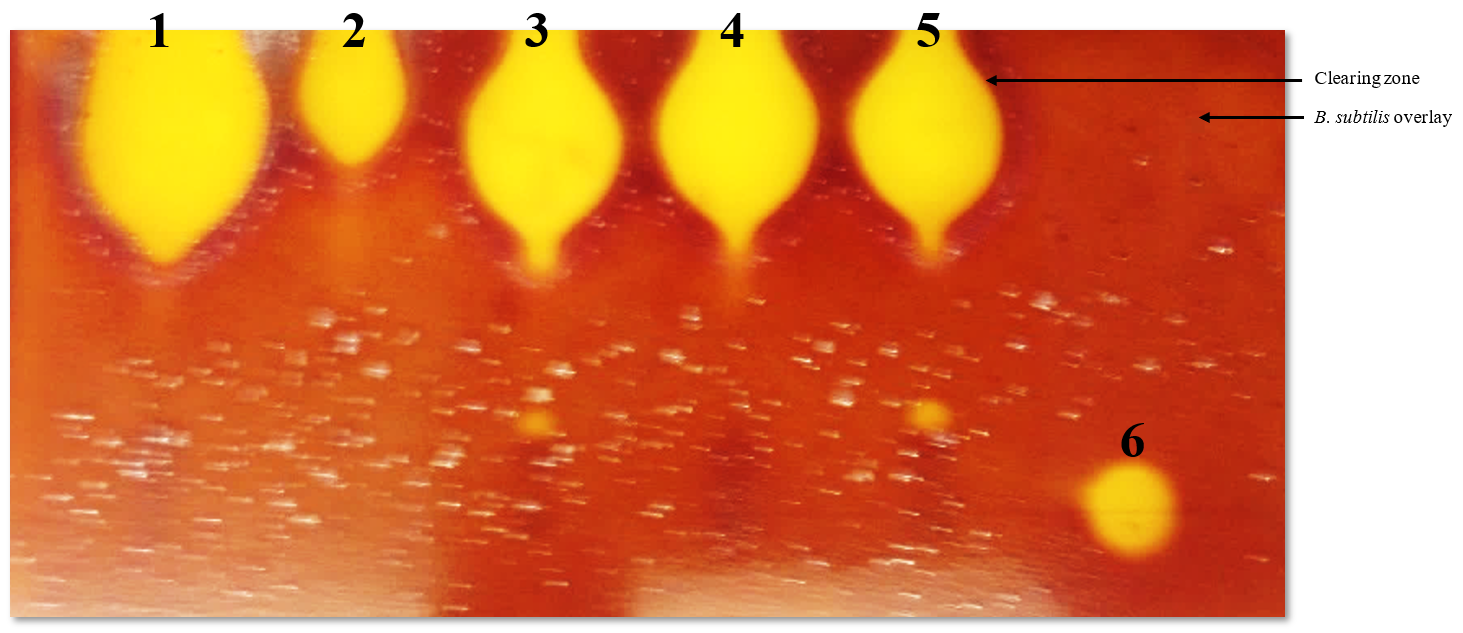

Figure S13: Direct bioautography of *H. fasciculare* crude extracts showing the antimicrobial activity of different culture crude extracts. 3 μl of 100 mg/ml^-1^ of mixed crude extract (supernatant+mycelia) of *H. fasciculare* of 1 = YMG, 2 = CSO, 3 = PDB, 4 = CGC, 5 = MEB media and 6 = 2 μl of 50 mg/ml of Kanamycin (positive control) loaded on TLC plates (developed in polar solvents) against *B. subtilis*.

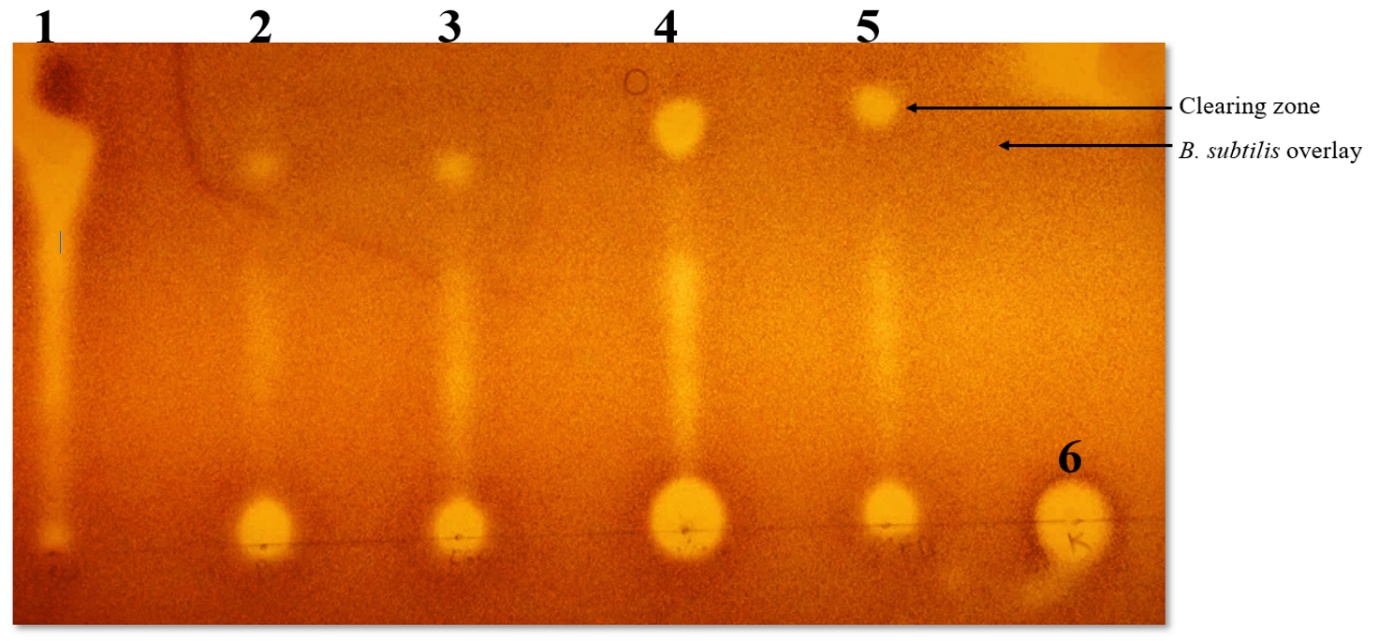

Figure S14: Direct bioautography of *H. fasciculare* crude extracts showing the antimicrobial activity of different culture crude extracts. 3 μl of 100 mg/ml^-1^ of mixed crude extract (supernatant+mycelia) of *H. fasciculare* of 1 = CSO, 2 = PDB, 3 = CGC, 4 =YMG, 5 = MEB media and 6 = 2 μl of 50 mg/ml of Kanamycin (positive control) loaded on TLC plates (developed in semi-polar solvents) against *B. subtilis*.

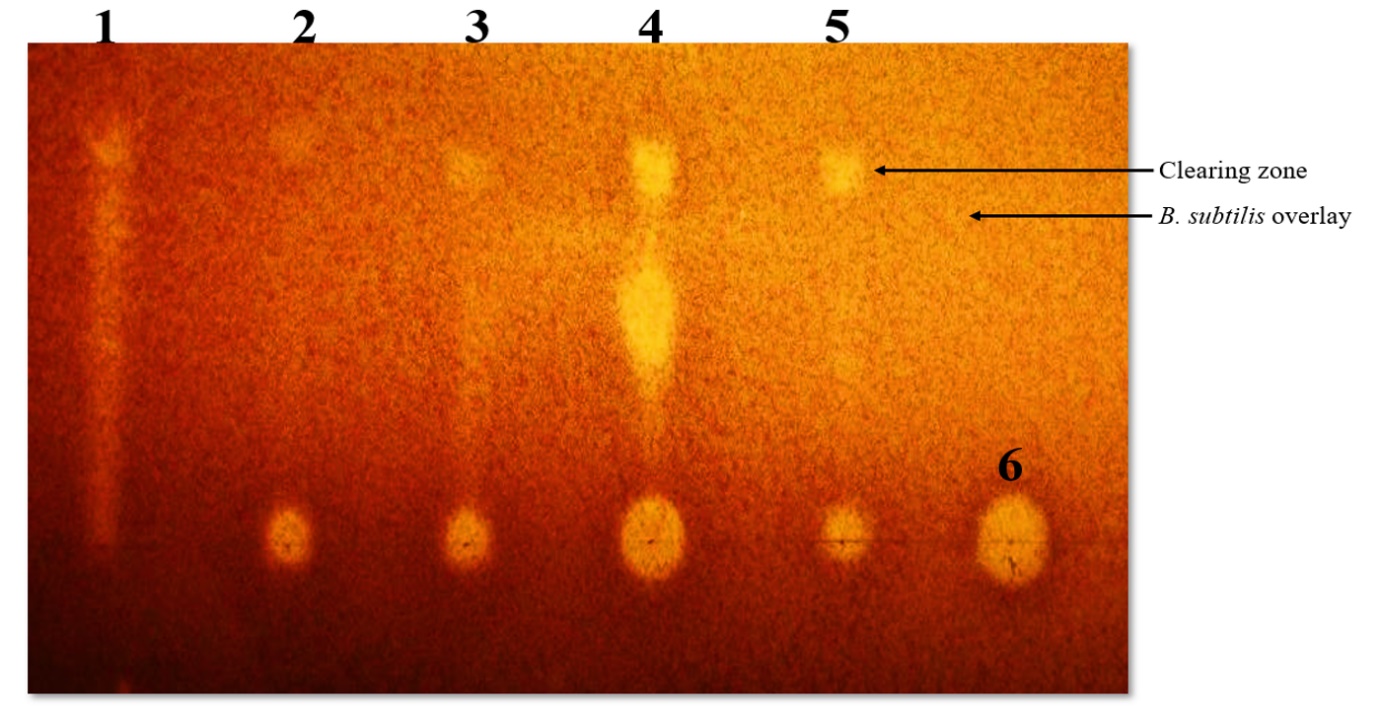

Figure S15: Direct bioautography of *H. fasciculare* crude extracts showing the antimicrobial activity of different culture crude extracts. 3 μl of 100 mg/ml^-1^ of mixed crude extract (supernatant+mycelia) of *H. fasciculare* of 1 = CSO, 2 = PDB, 3 = CGC, 4 =YMG, 5 = MEB media and 6 = 2 μl of 50 mg/ml of Kanamycin (positive control) loaded on TLC plates (developed in non-polar solvents) against *B. subtilis*.

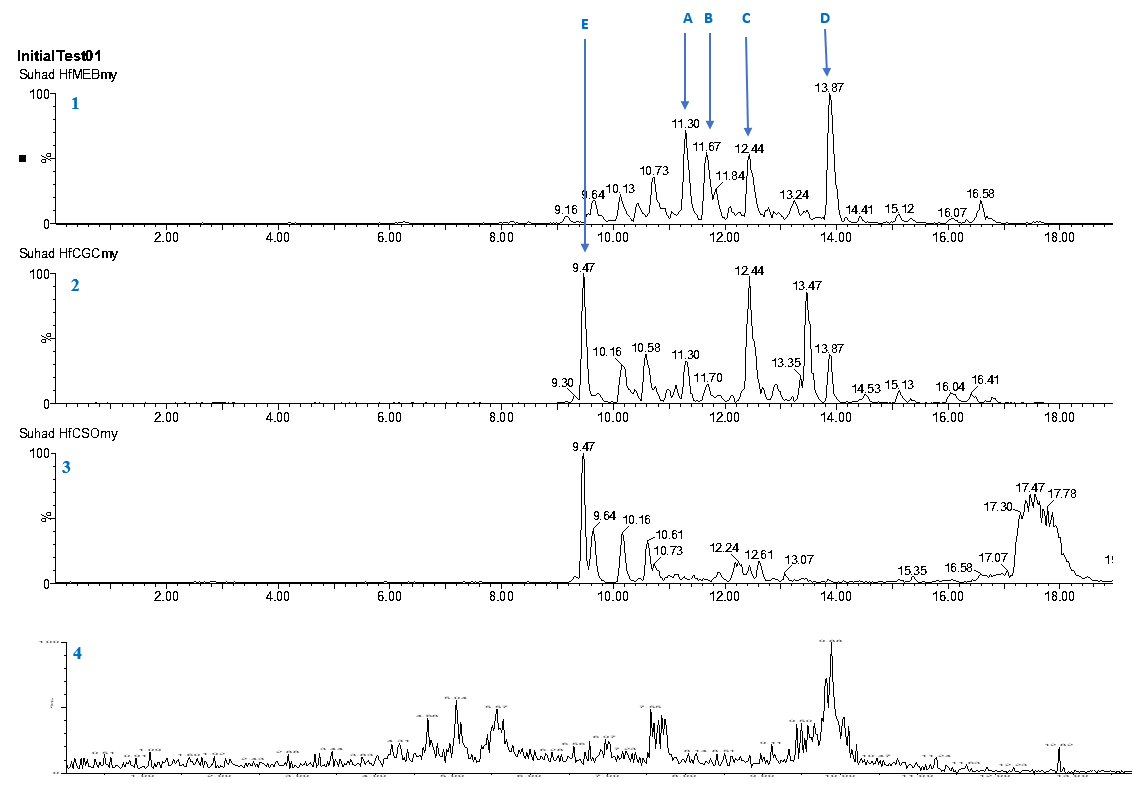

Figure S16: LC-MS chromatograms electrospray mass negative mode (ES-) of 5 mg/ml of:

1- *H. fasciculare* MEB crude extract. 2- *H. fasciculare* CGC crude extract. 3- *H. fasciculare* CSOA-1 crude extract. 4- *H. fasciculare* YMG crude extract

A = Facicularone G. B = naematolin. C = Unknow. D = Unknown. E = Unknow.

Figure S17: ^1^H NMR (125MHZ) spectrum of 3,5-dichloro-4-methoxybenzoic acid (3,5 D).

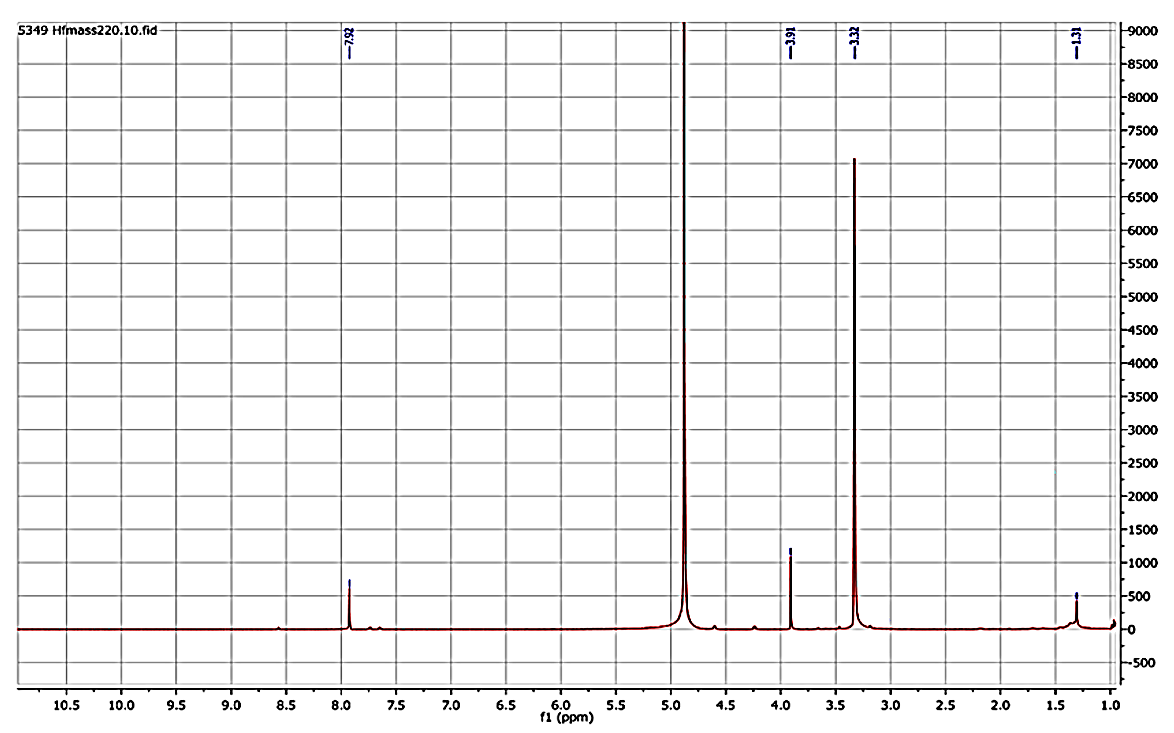

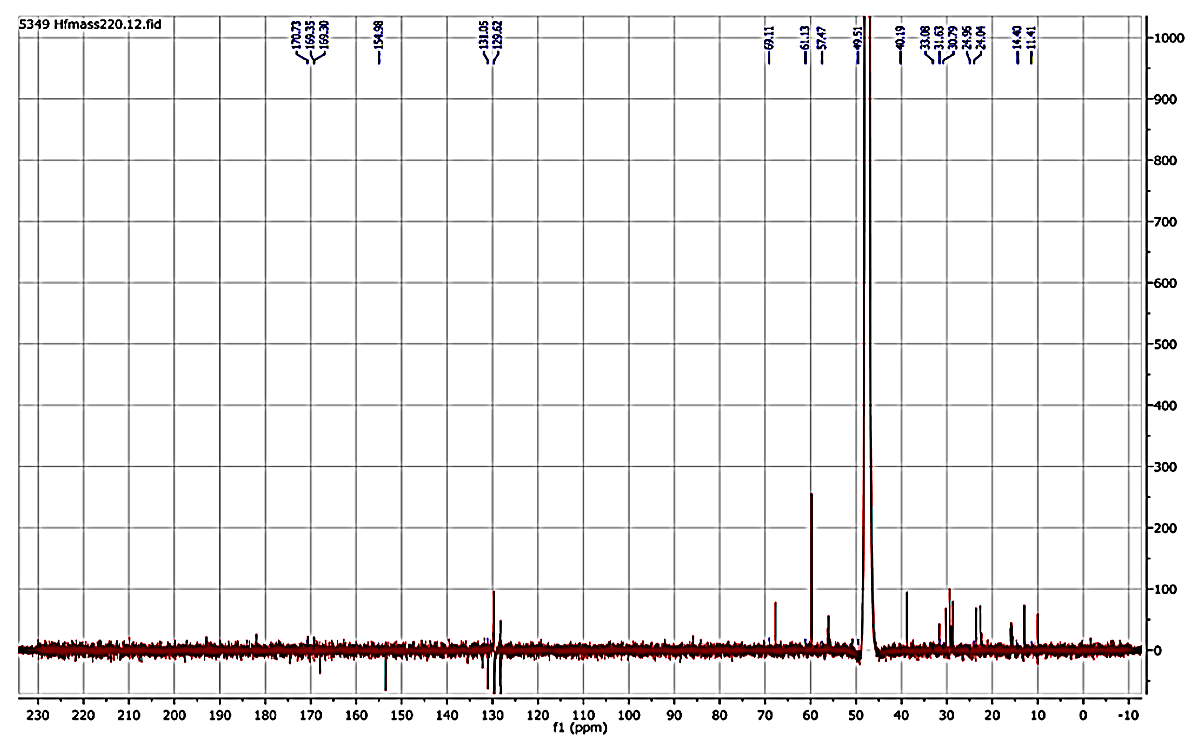

Figure S18: ^13^C NMR (500MHZ) spectrum of 3,5-dichloro-4-methoxybenzoic acid (3,5 D).

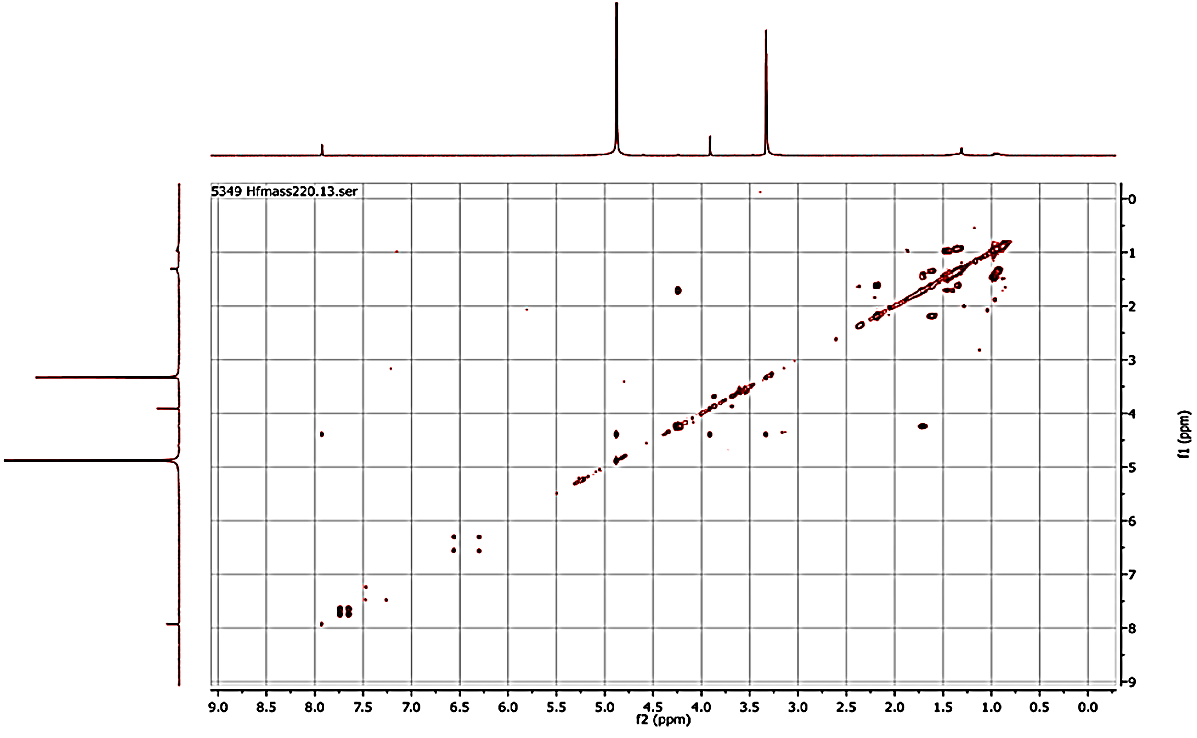

Figure S19: COSY (500MHZ) spectrum of 3,5-dichloro-4-methoxybenzoic acid (3,5 D).

Figure S20: HSQC (500MHZ) spectrum of 3,5-dichloro-4-methoxybenzoic acid (3,5 D).

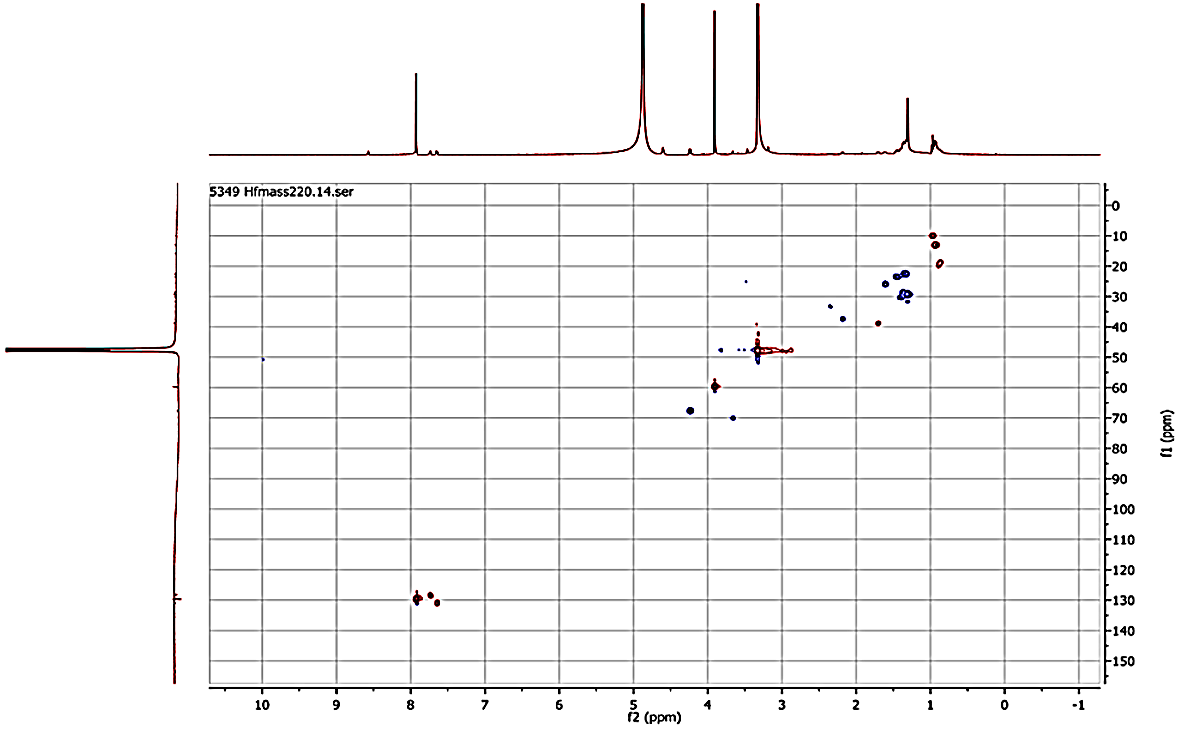

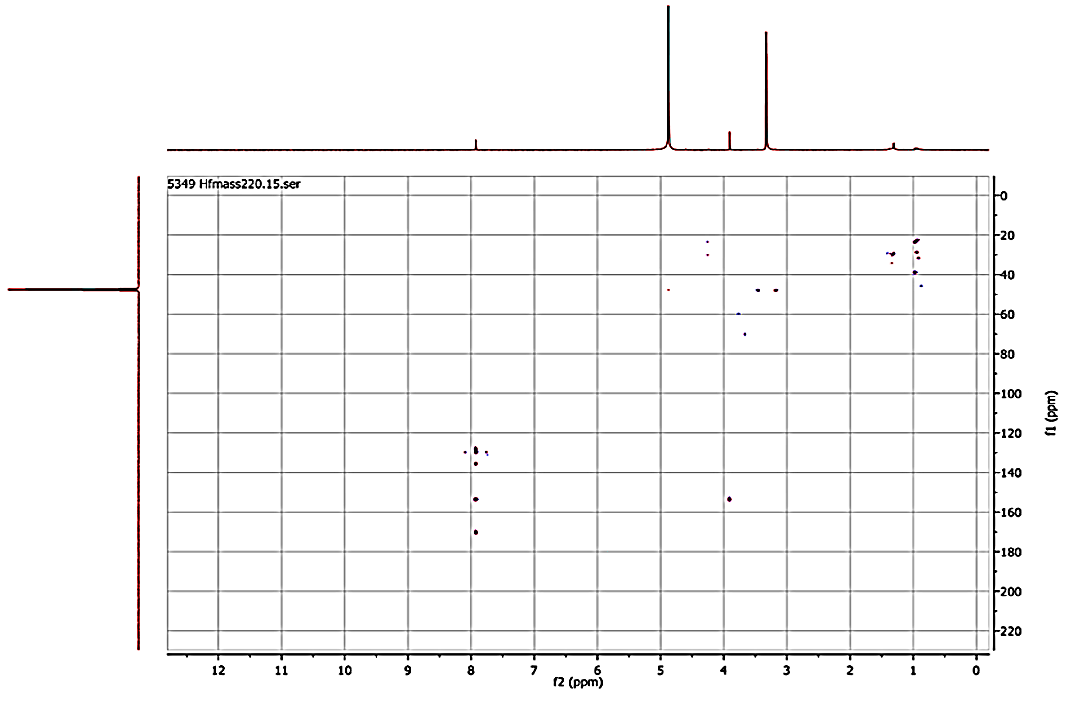

Figure S21: HMBC (500MHZ) spectrum of 3,5-dichloro-4-methoxybenzoic acid (3,5 D).

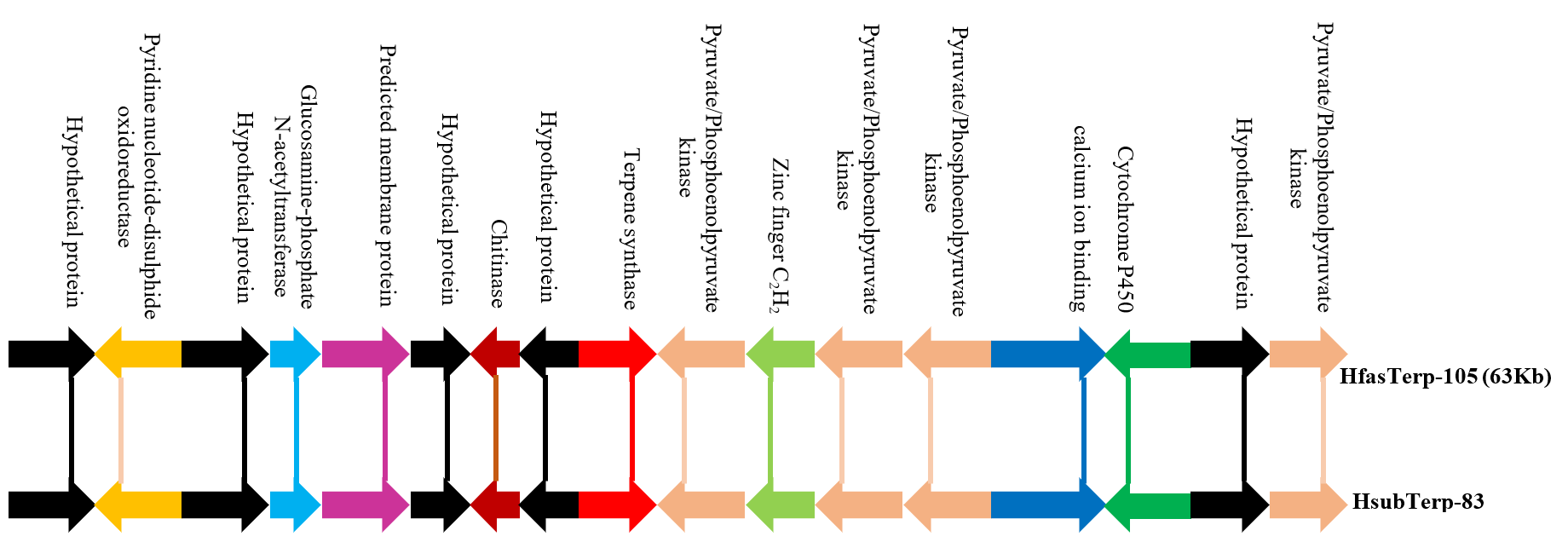

**Figure S22: Predicted gene cluster of Hfas-terp105 and the homologous cluster (Hsub-terp83) in *H. sublateritium***.

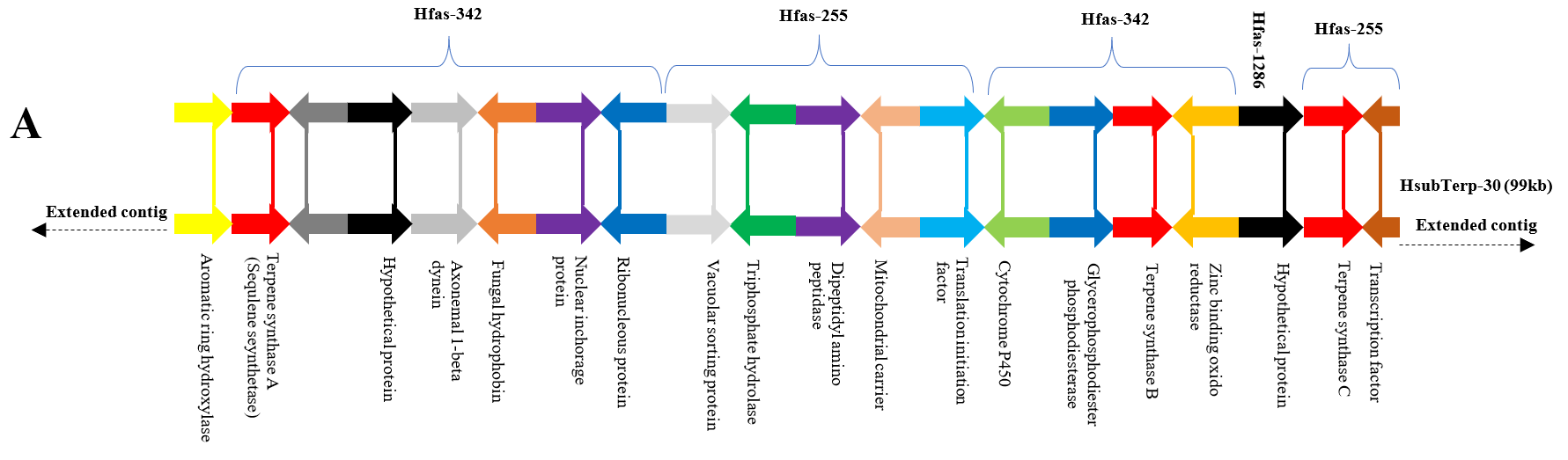

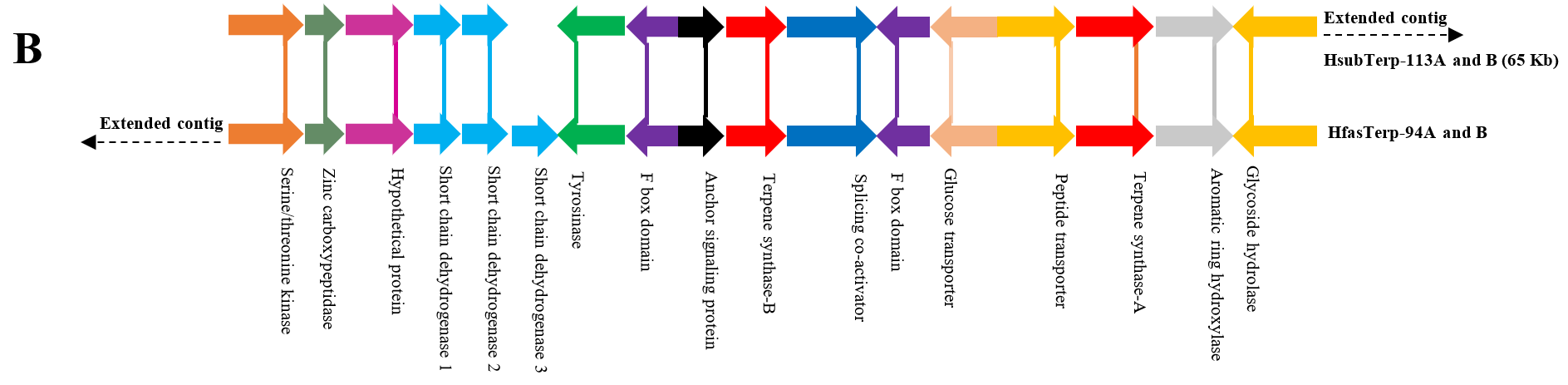

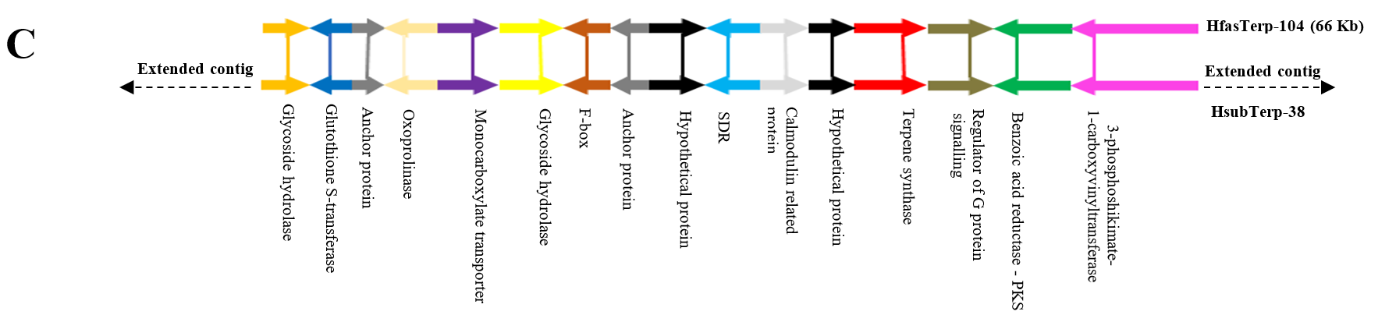

**Figure S23: Predicted gene cluster of *H. sublateritium* terpene synthases and *H. fasciculare*.**

A. *H. sublateritium* scaffold 30 terpene synthase gene clusters and the homologous clusters of *H. fasciculare*.
B. *H. sublateritium* scaffold 113 terpene synthase gene clusters and the homologous clusters of *H. fasciculare*.
C. *H. sublateritium* scaffold 38 terpene synthase gene clusters and the homologous clusters of *H. fasciculare*.

**A**

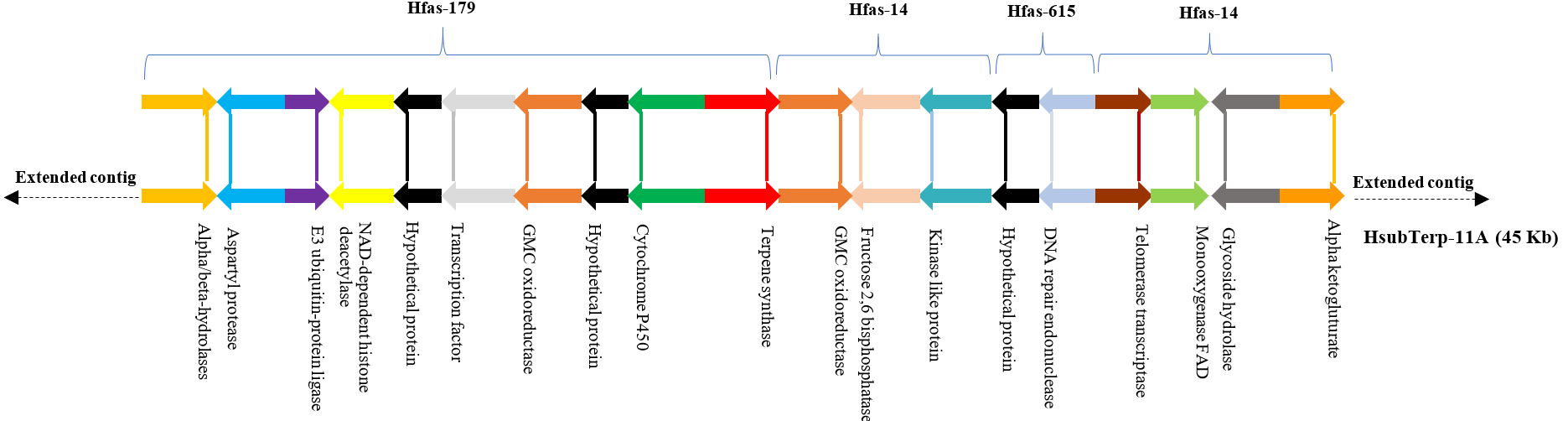

**B**

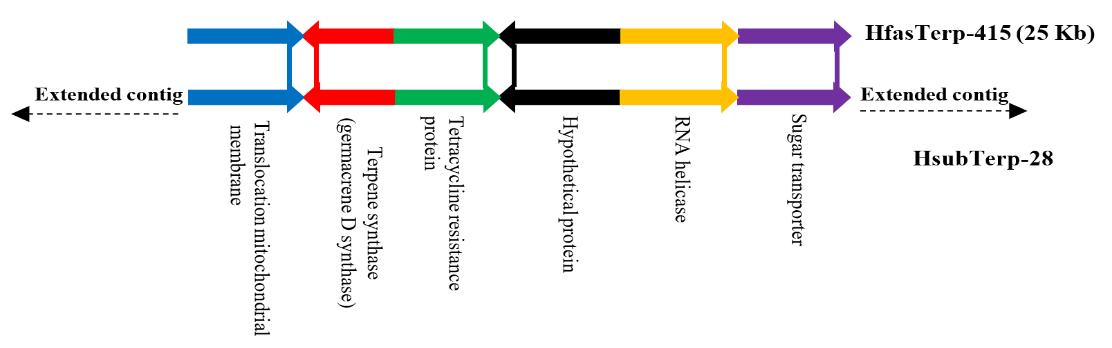

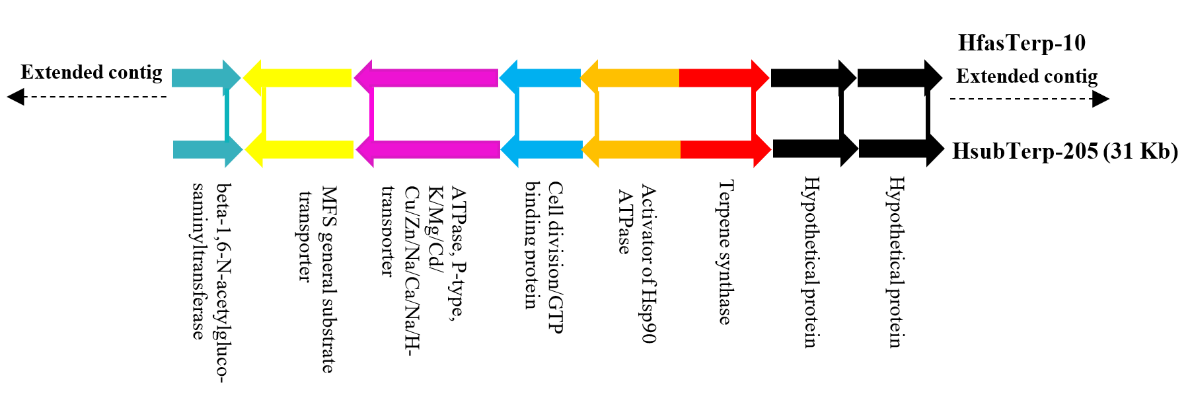

**C**

**Figure S24: Predicted gene cluster of *H. sublateritium* terpene synthases and *H. fasciculare*.**

A. *H.* sublateritium scaffold 11A terpene synthase gene clusters and the homologous clusters of *H. fasciculare*.
B. *H. sublateritium* scaffold 205 terpene synthase gene clusters and the homologous clusters of *H. fasciculare*.
C. *H. sublateritium* scaffold 28 terpene synthase gene clusters and the homologous clusters of *H. fasciculare*.

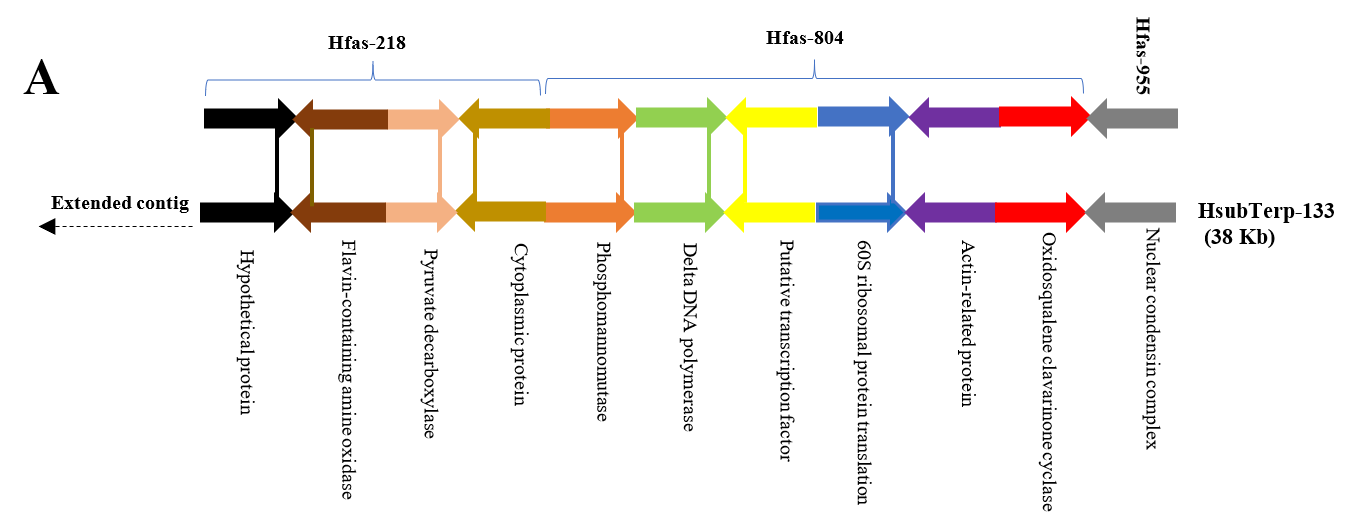

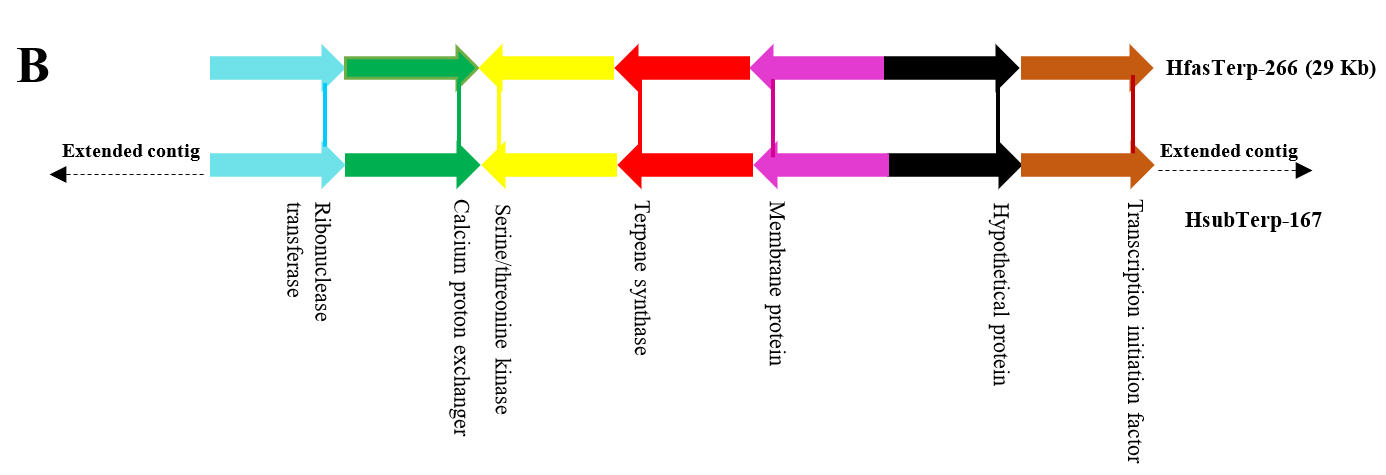

**Figure S25: Predicted gene cluster of *H. sublateritium* terpene synthases and *H. fasciculare*.**

A. *H. sublateritium* scaffold 133 terpene synthase gene clusters and the homologous clusters of *H. fasciculare*.
B. *H. sublateritium* scaffold 167 terpene synthase gene clusters and the homologous clusters of *H. fasciculare*.

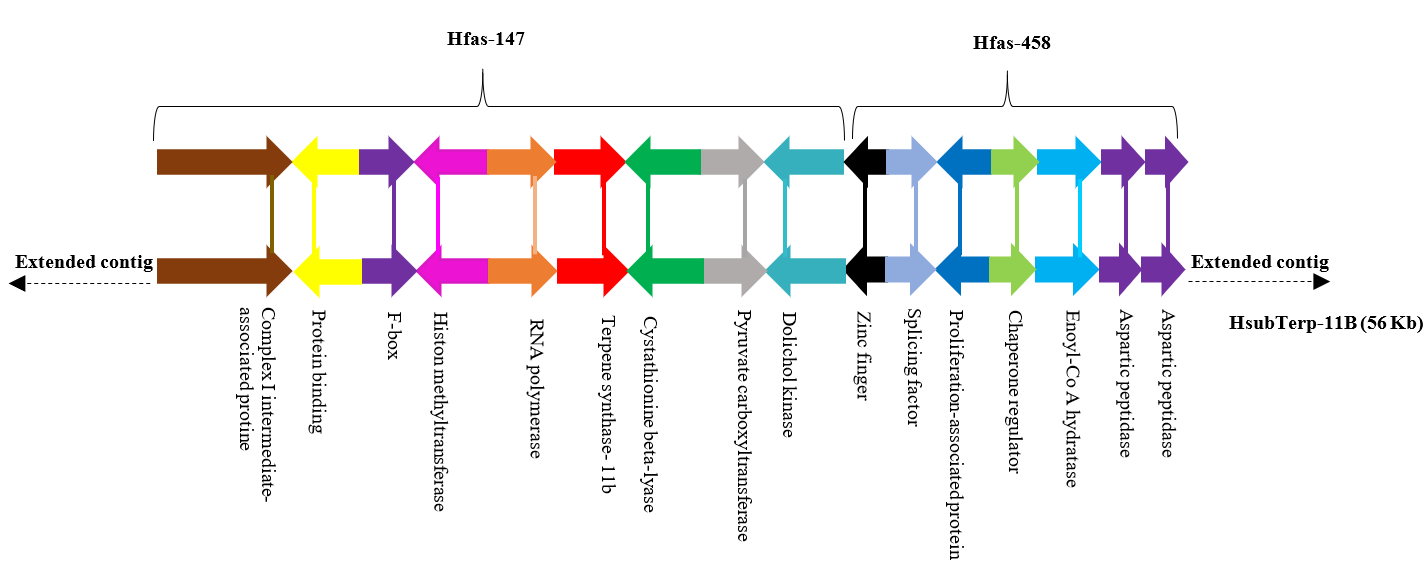

**Figure S26: Predicted gene cluster of *H. sublateritium* terpene synthase 11B and the homologous cluster in *H. fasciculare*.**

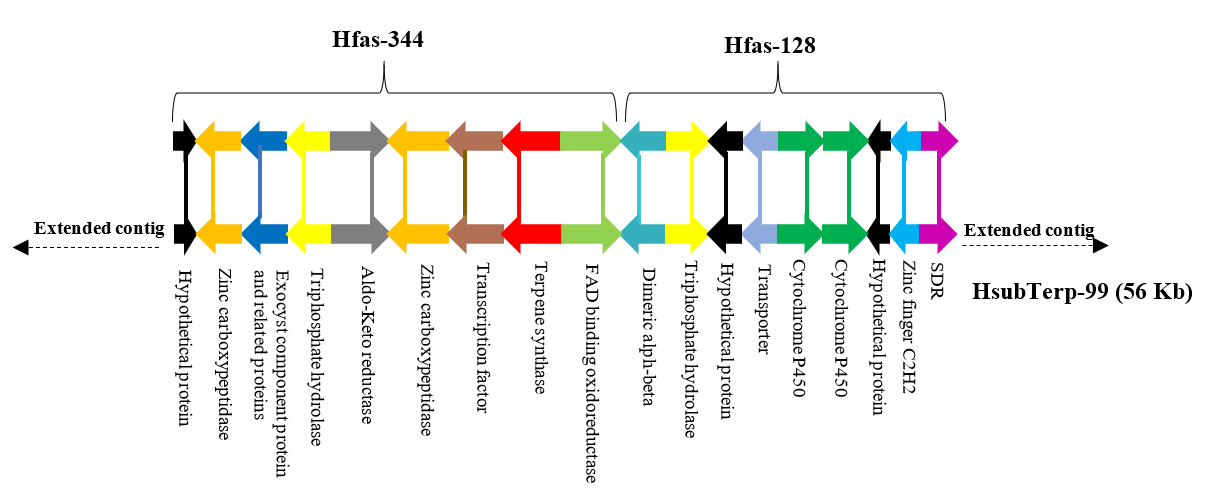

**Figure S27: Predicted gene cluster of *H. sublateritium* terpene synthase 99 and the homologous cluster in *H. fasciculare*.**

**Figure S28: Predicted gene cluster of *H. fasciculare* and *H. sublateritium* PKS.**

A. *H. fasciculare* PKS 102 biosynthetic gene cluster and the homologous cluster of *H. sublateritium.*
B. *H. fasciculare* PKS 221 biosynthetic gene cluster and the homologous cluster of *H. sublateritium.*

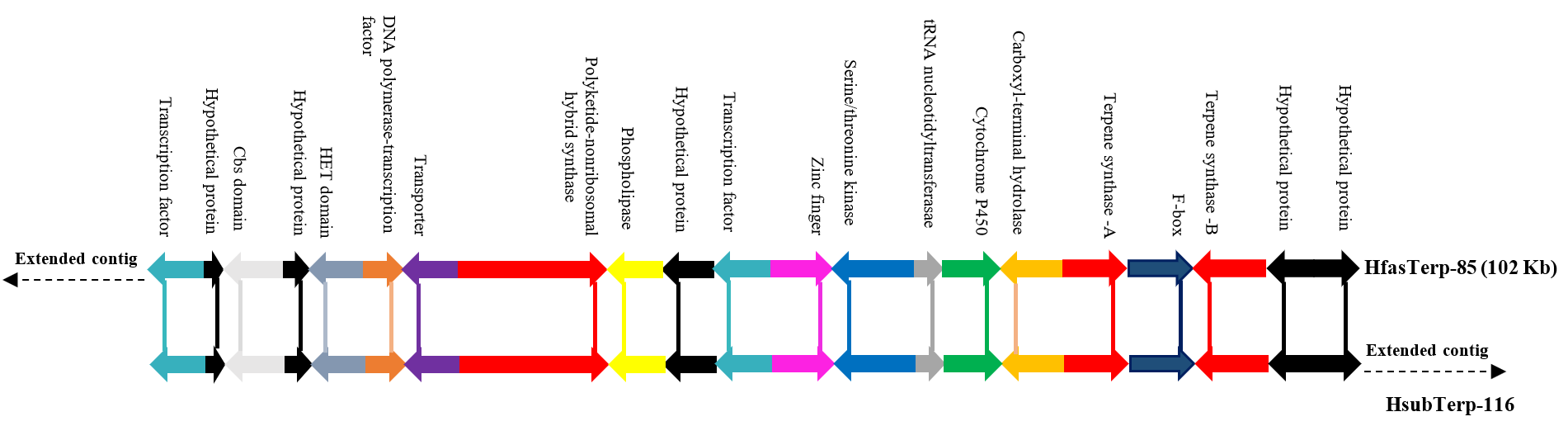

**A**

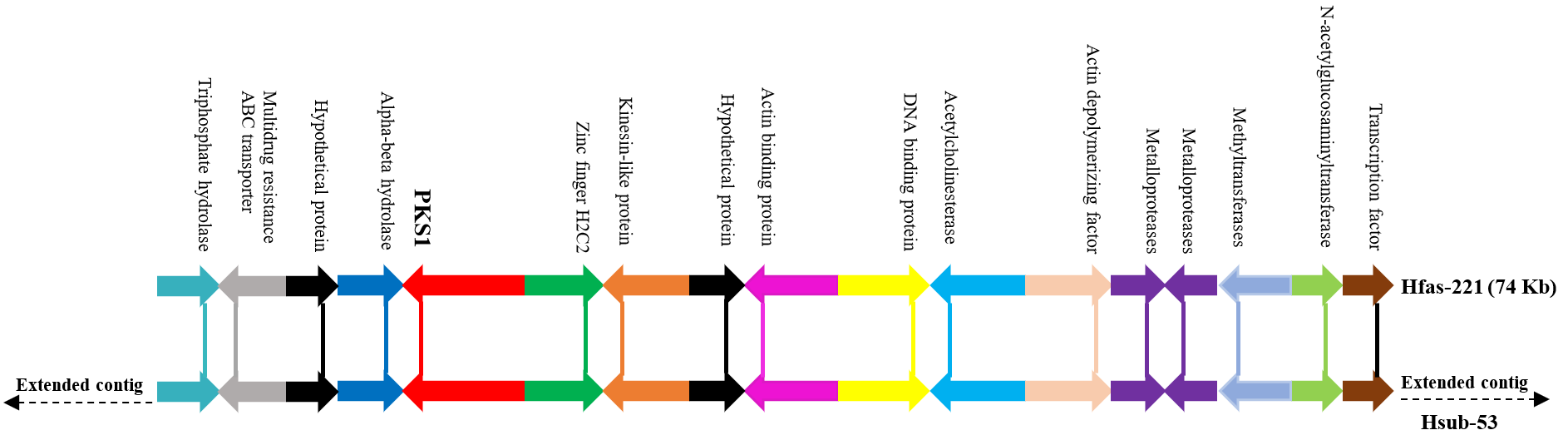

**B**

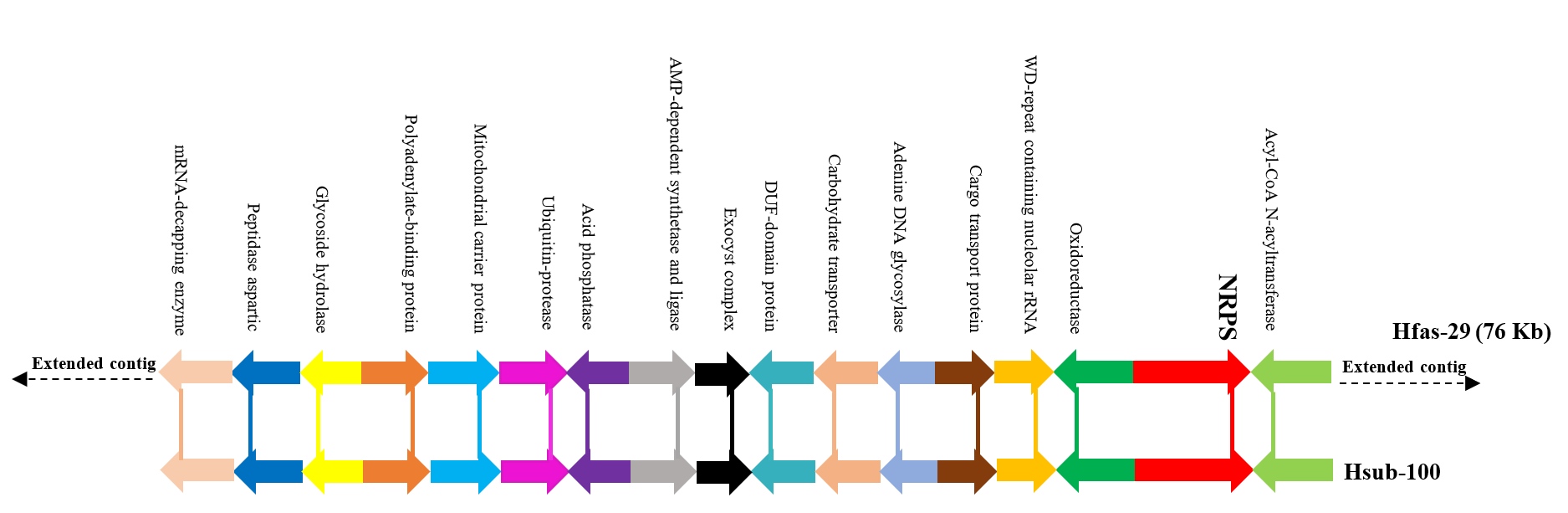

**A**

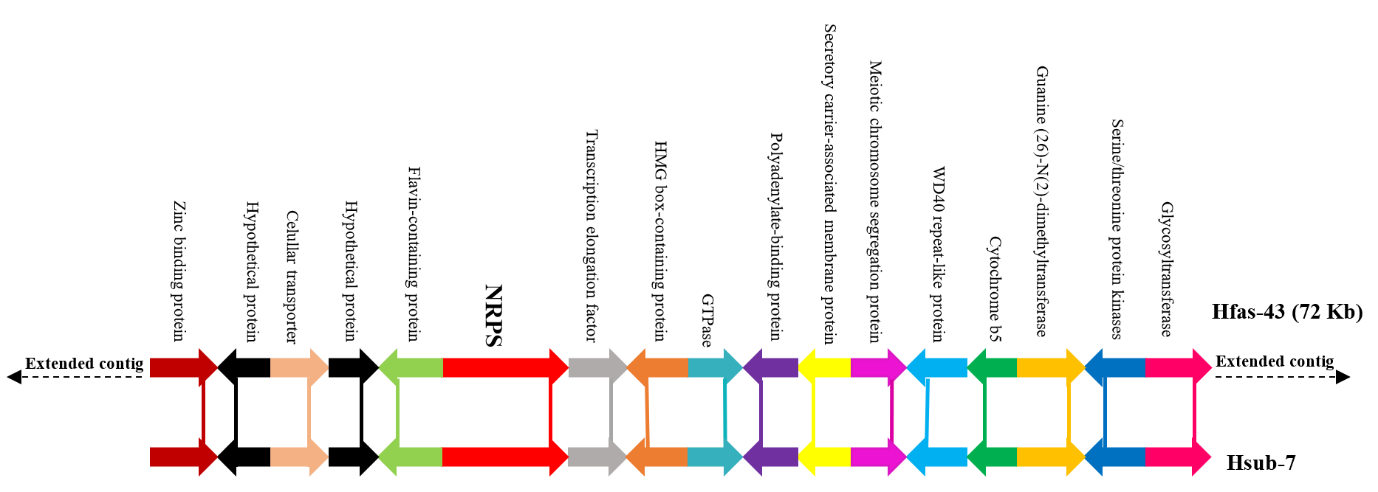

**B**

**Figure S29: Predicted gene cluster of *H. fasciculare* and *H. sublateritium* NRPS.**

A. *H. fasciculare* NRPS 29 biosynthetic gene cluster and the homologous cluster of *H. sublateritium*.
B. *H. fasciculare* NRPS 43 biosynthetic gene cluster and the homologous cluster of *H. sublateritium*.

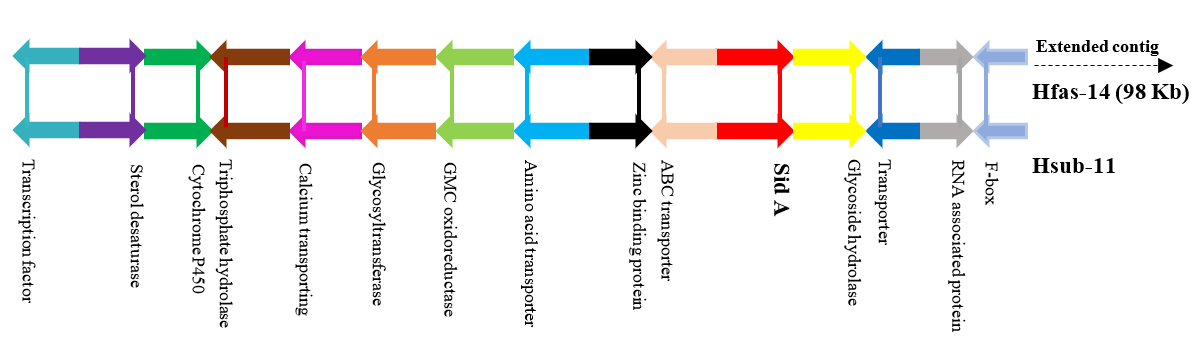

**Figure S30: Predicted gene cluster of *H. sublateritium* SidA and the homologous cluster in *H. fasciculare*.**

**Materials and Methods**

**Bioassays**

**Plate based bioassay**

**Preparing of Bacillus subtilis spores’ suspension**

The method of Hartley et al. (2009) was used to prepare a *B. subtilis* spore suspension, in which glycerol stock of *Bacillus* was used to inoculate 50 ml culture of tryptic soy broth (TSB), and this was incubated for 24 hours at 30°C with shaking at 200 rpm. 1ml of this overnight culture was spread with sterile glass beads on the surface of tryptic soy agar slant (prepared by adding 75 ml of tryptic soy agar (TSAb) to Thompson bottle, and allowed to set horizontally after autoclaving. The cells were harvested after 6 days incubation at 30°C by using again sterile glass beads and 100 ml of sterile deionised water. *Bacillus* suspension was then heated at 70°C in a water bath for 30 minutes to activate the spores and kill the vegetative cells. These spores were subsequently stored at 4°C until required.

**Preparation of overlays**

*B. subtilis* overlay was prepared by adding 800 µl of *B. subtilis* spore suspension to 400 ml molten overlay medium (TSAg 70°C) to heat shock the spores. To visualize the overlain bacteria, 2 ml of 0.4% 2,3,5-triphenyl-2H-tetrazolium chloride (TTC) was added to the cooled agar (approximately 50°C), before pouring the overlay.

For *E. coli* overlay preparation, 5 ml of LB broth was inoculated with a colony from a fresh DH5α plate, before incubating overnight at 200 rpm at 37°C. This culture was then transferred to a sterile Erlenmeyer flask containing 100 ml of fresh LB medium and incubated at 200 rpm, 37°C until the optical density OD600 of the culture reached 1.0. 2 ml of the bacterial suspension and 0.4% of 2,3,5-triphenyl-2H-tetrazolium chloride (TTC) were then added to 1L of the molten (approximately 50°C) overlay medium LBA.

The *S. cerevisiae* culture was prepared by inoculating 5 ml of YPD medium with a yeast colony (Y10000), before being shaken at 200 rpm at 28°C for 24 hours. The overnight culture was then transferred to 100 ml of YPD broth, and shaken as before. Once the optical density OD600 reached 1.0, 2 ml of this culture and 0.4% 2,3,5-triphenyl-2H-tetrazolium chloride (TTC) were added to 1 L of the molten medium (YPDA) after cooling the agar to approximately 50°C.

To estimate the antimicrobial activity of basidiomycete isolates against *S. cerevisiae* (Eukaryote), *E. coli* (Gram negative bacteria) and B. subtilis (Gram positive bacteria), a plug of fresh growing mycelium of these fungi was transferred to the centre of 90 mm petri dishes of appropriate medium and incubated for 5-7 days. These plates were then overlaid with 5-10 ml of microbial overlay medium, before secondary incubation. The temperature and incubation period of this further incubation depends on the challenging microorganism. The incubation conditions for *S. cerevisiae* and *B. subtilis* overlays were 48 hrs at 30°C, and for *E. coli* were 24 hours at 37°C. Both fungal growth and the clearing zone diameter were measured. This plate based bioassay was performed in 3 replicates for each tested basidiomycete, each growth medium and each challenging microbe.
